# Genotyping common, large structural variations in 5,202 genomes using pangenomes, the Giraffe mapper, and the vg toolkit

**DOI:** 10.1101/2020.12.04.412486

**Authors:** Jouni Sirén, Jean Monlong, Xian Chang, Adam M. Novak, Jordan M. Eizenga, Charles Markello, Jonas A. Sibbesen, Glenn Hickey, Pi-Chuan Chang, Andrew Carroll, Namrata Gupta, Stacey Gabriel, Thomas W. Blackwell, Aakrosh Ratan, Kent D. Taylor, Stephen S. Rich, Jerome I. Rotter, David Haussler, Erik Garrison, Benedict Paten

## Abstract

We introduce Giraffe, a pangenome short read mapper that can efficiently map to a collection of haplotypes threaded through a sequence graph. Giraffe, part of the variation graph toolkit (vg)^1^, maps reads to thousands of human genomes at around the same speed BWA-MEM^2^ maps reads to a single reference genome, while maintaining comparable accuracy to VG-MAP, vg’s original mapper. We have developed efficient genotyping pipelines using Giraffe. We demonstrate improvements in genotyping for single-nucleotide variants (SNVs), small insertions and deletions (indels) and structural variations (SVs) genome-wide. We use Giraffe to genotype about 167 thousand structural variants ascertained from long read studies in 5,202 human genomes sequenced with short reads, including the complete 1000 Genomes Project dataset, at an average cost of $1.50 per sample. We determine the frequency of these variations in diverse human populations, characterize their complex allelic variations and identify thousands of expression quantitative trait loci (eQTLs) driven by these variations.

## Introduction

Relative to existing linear reference genomes, pangenomes hold promise as reference structures for improving common genomic tasks, such as genotyping, by more fully capturing prior information about genetic variation^3–5^. They have been shown to improve the genotyping of structural variants (SVs, ≥ 50 bp)^6^, and may particularly improve the genotyping of all forms of variation within highly polymorphic and repetitive regions. SVs affect millions of bases within each human genome but are much more poorly characterized compared to single-nucleotide variants (SNVs) and short insertions/deletions (indels,<50 bp)^7^,^8^, largely because of reference bias created by mapping to single linear reference genomes. Similarly, genotyping in highly polymorphic or repetitive regions has proven challenging^9^ because information about orthologous haplotypic variation that can be used to disambiguate mappings is, by definition, not captured by existing linear reference genomes.

To date pangenomes, which encode information about many genomes, have been described by structures that can be formulated as sequence graphs^10^. A number of pangenome mapping algorithms have been developed for mapping reads to sequence graphs, but none has yet challenged the dominance of linear reference genome mapping algorithms within the community. The original VG-MAP^1^ algorithm is able to map to sequence graphs with arbitrary topology, including those containing cycles, as can be produced by duplication and complex rearrangement, but while accurate VG-MAP is at least an order of magnitude slower than popular linear genome mappers, such as BWA-MEM^2^, for mapping to human genomes. Given that mapping is frequently a bottleneck in genome analysis, the cost of VG-MAP has proven prohibitive. The pangenome mapper HISAT2^11^ is extremely fast, but is limited to mapping to graphs containing only SNVs and indels at relatively low density. As we will show, for genome mapping, rather than transcriptome mapping (where HISAT2 is well-used), it is also of limited accuracy. The recently developed GraphAligner^12^ is able to map to arbitrary sequence graphs but is designed for long reads and, in our experiments, performs poorly with short reads. The Seven Bridges GRAF software^13^ combines an integrated pangenome mapper and genotyping algorithm, but is closed-source, currently specialized to work only with a customized human pangenome that is not public, and is not available for general testing or customization.

Here we introduce a new pangenome mapper, Giraffe, that focuses on mapping to collections of aligned haplotypes. It was created with the objective of being the first open-source, practically fast mapper that can be used to map pangenomes containing a high density of all forms of variation. We show Giraffe improves genotyping of SNVs, indels and SVs genome-wide, and use it to cheaply and efficiently characterize SVs in thousands of genomes.

## Results

### Giraffe: Fast, Haplotype Aware Pangenome Mapping

Giraffe is a short read to graph mapper designed to map to haplotypes, producing alignments embedded within a sequence graph^3^ (Figure 1(A)). We take advantage of the fact that the vast majority of Illumina sequencing errors are substitutions, and we further assume that the common indels are already present in the haplotypes. Hence, we try to align the read without gaps before resorting to dynamic programming.

**Figure 1.**
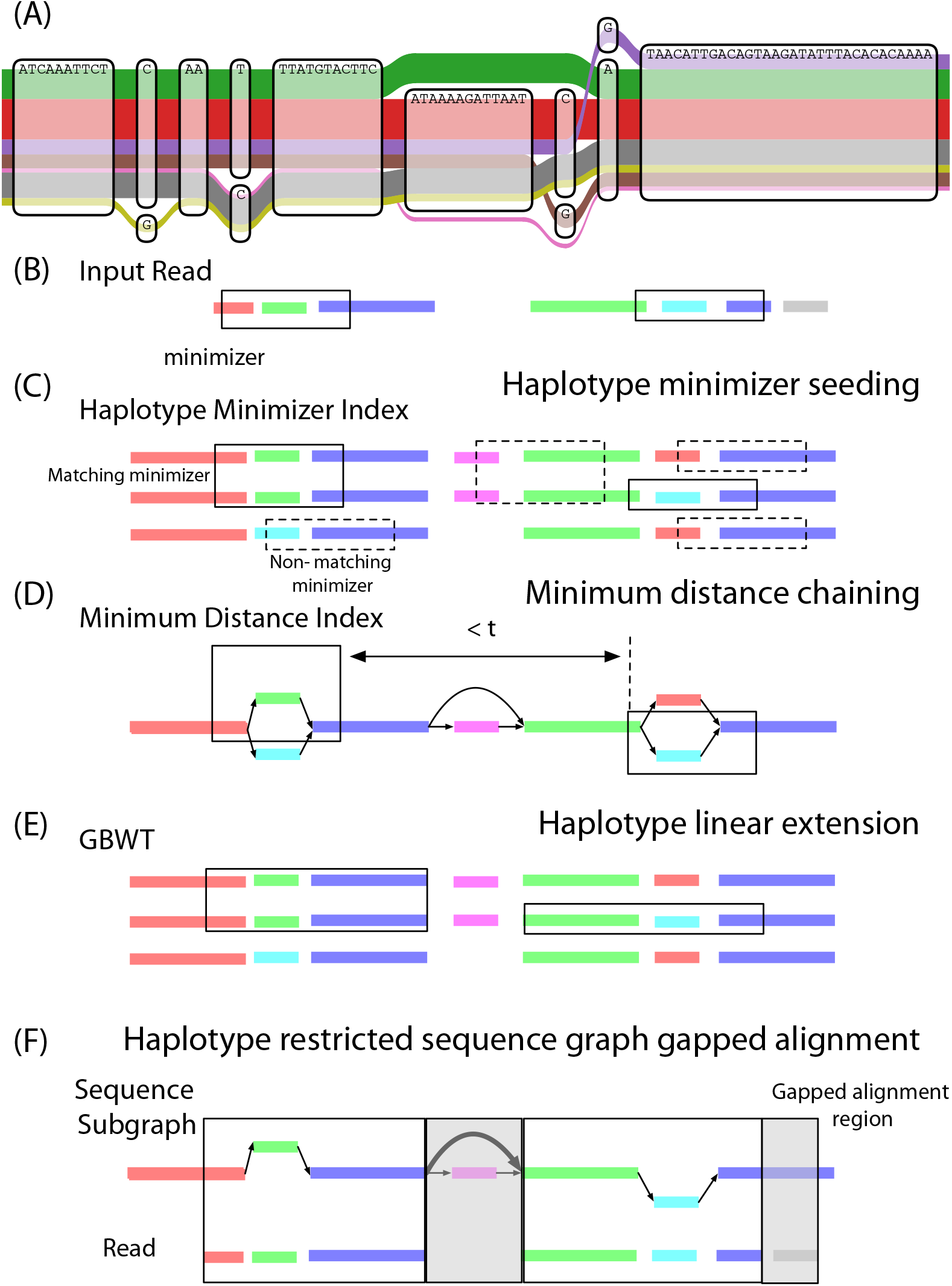
Haplotype mapping. (A) A region of the *CASP12* gene in the 1KG graph (see Methods), on path 11 from 104, 898, 041 bp to 104, 898, 120 bp, illustrating complex local variation. The number of possible paths through a sequence graph generally grows exponentially with size. However, common haplotypes (the colored ribbons of width log-proportional to population frequency) represent a tiny subset of these possible paths. Mapping to embedded haplotypes therefore vastly reduces the size of the mapping space. (B-F) An overview of Giraffe. (B) The input read is represented as a sequence of aligned, colored rectangles corresponding to the haplotypes, which are similarly represented below. (C) Haplotype minimizer seeding: The minimizers in the read are enumerated. The matching minimizers in the haplotypes are identified. (D) Minimum distance chaining: Minimizer instances in the graph are clustered by minimum distance. (E) Haplotype linear extension: Minimizers in high scoring clusters are extended linearly to form maximal gapless local alignments. (F) Haplotype restricted sequence graph gapped alignment: Any remaining gaps in the alignment between read and graph are resolved by gapped alignment.

We use the *GBWT index*^14^, a compressed and self-indexed representation for large numbers of haplotypes embedded in a graph, which recasts the graph as an alignment of those haplotypes. The graph describes which positions in the haplotypes are equivalent, while the haplotypes describe the subset of the possible paths in the graph that are worth considering. In Giraffe, we work in graph coordinates, and map reads to the graph, but we usually restrict our attention to paths that are consistent with the haplotypes. This approach allows us to deal effectively with complex graph regions, where the number of possible paths may be excessively large, but the vast majority of them represent rare or nonexistent sequences.

The main stages of the Giraffe algorithm, as illustrated in Figure 1(B-F), are:

1. Find seeds using a *minimizer index*^15^. Because we assume that sequencing errors are rare and the graph already contains common variants, we can use longer minimizers than in most other applications.
2. Cluster the seeds using a *distance index*^16^. The index is a tree based on a hierarchical decomposition^17^ of the graph. We can quickly compute the shortest distance between any two positions in the graph by traversing the tree.
3. Gaplessly extend the seeds in each cluster, allowing for a limited number of mismatches.
4. If we did not find enough full-length alignments, fully align the most promising clusters. The extended seeds from the previous stage serve as seeds for dynamic programming.

We present two parameterizations of Giraffe: *default Giraffe*, also written as just “Giraffe”, balances speed and accuracy, while *fast Giraffe* is optimized for speed at the expense of some accuracy.

### Simulation Study

Lacking a true gold standard for mapping evaluation, we undertook a comprehensive simulation study. We used read simulations for three different Illumina sequencing technologies, and mapped reads to three different graphs. We performed separate evaluations of single and paired end mapping.

#### Graphs

We evaluated Giraffe on two human genome reference graphs: one built with small variants (<50 bp) from the 1000 Genomes Project^18^ (the *1KG graph*) and one with SVs (≥50 bp) from the Human Genome Structural Variant Consortium^19^ (the *HGSVC graph*). Both were built by inducing graphs from phased variant call format (VCF) files (see Methods). The 1KG graph, which uses the GRCh37 assembly as a reference, contains data from 2,533 individuals, of which 2,503 had haplotypes on the autosomes and 2,532 had mitochondrial haplotypes. It represents 81,478,306 SNVs, 3,485,914 small indels (<50 bp), and 2,474 larger SVs (≥50 bp). The HGSVC graph contains data from three individuals sequenced with long-reads, HG00514, HG00733, and NA19240, and uses the GRCh38 assembly as a reference. The HGSVC graph contains 78,106 larger SVs (≥50 bp).

To evaluate Giraffe’s performance on more diverged, non-human data we used a Cactus multiple sequence alignment for five strains of the *S. cerevisiae* and *S. paradoxus* yeasts, without including any human sequences^6^. The resulting *yeast graph* is challenging to work with because it contains the cycles and duplications typical of graphs generated from genome-wide multiple sequence alignments of more divergent sequences. Although its complex topology can make categorizing the variants in the yeast graph ontologically challenging, it contains 4,833,776 nodes, 6,542,993 edges, and 14,926,050 bp of sequence. Using a graph decomposition technique^17^, we find it contains 1,459,769 variant sites, 90 of these sites are defined as complex, being not directed, acyclic, and free of internal source and sink nodes.

To allow more direct comparison of Giraffe to linear mappers, we created three *primary graphs* containing only the primary linear reference genome (GRCh37, GRCh38, or *S*.*c. S288C* assembly) with no variants or alternative loci, which we used as matched negative controls for the 1KG, HGSVC, and yeast graphs, respectively.

#### Reads

To evaluate Giraffe for mapping human data, we obtained paired-end sequencing reads from an Illumina NovaSeq 6000 for sample NA19239 (accession ERR3239454) and from Illumina HiSeq 2500 and HiSeq X Ten machines for sample NA19240 (accessions ERR309934 and SRR6691663, respectively). These samples are a parent and child in a pedigree, and were selected because the child NA19240 has known genotypes for HGSVC variants^19^, while the parent NA19239 has known genotypes for the 1KG variants^18^. Both samples were excluded from the 1KG graph (see Methods). For each sample and read technology, we simulated 1 million read pairs (2 million reads). To evaluate Giraffe for mapping non-human data we similarly simulated 500,000 read pairs using a model of Illumina HiSeq 2500 reads, simulating reads from a held out yeast strain, DBVPG6044, not included in the yeast graph (see Methods).

#### Evaluation

Read sets were mapped to the graphs using four genome graph mappers: Giraffe, VG-MAP^1^, HISAT2^11^, and GraphAligner^12^. HISAT2 and GraphAligner could not be run on all graphs. We were unable to build a HISAT2 index for the full 1KG graph, and so instead we mapped it to a subset of the 1KG data using the graph from the authors paper^11^ that contains approximately 1/6th of the variations. Similarly, neither HISAT2 nor GraphAligner could be made to map reads to the yeast graph; in the case of HISAT2, this was because it cannot map to graphs containing cycles. In addition, we mapped the read sets to the primary graphs using Giraffe, and to the linear reference assemblies using the linear sequence mappers BWA-MEM^2^, Bowtie2^20^, and Minimap2^21^.

Figure 2, Supplementary Figures S2, S3, and S4 and Supplementary Tables S1,S2,S3,S4,S5 and S6 compare the accuracy of single and paired-end mappings. At the highest reported mapping quality VG-MAP and default Giraffe consistently have either higher precision or recall than all other assessed mappers across all simulated read technologies and graphs, and are generally similar to each other. Relative to the linear mappers, the Giraffe and VG-MAP lead is larger for the HGSVC graph than the 1KG graph, and larger still for the more divergent, sibling-species-scale yeast graph, indicating that the gains to be had from using a genome graph are higher when the graph contains larger structural variants or more divergent sequences. GraphAligner, which was designed for long reads, appears poorly suited for mapping short reads. HISAT2 appears unable to stratify reads by mapping quality to produce a high-precision tranche.

**Figure 2.**
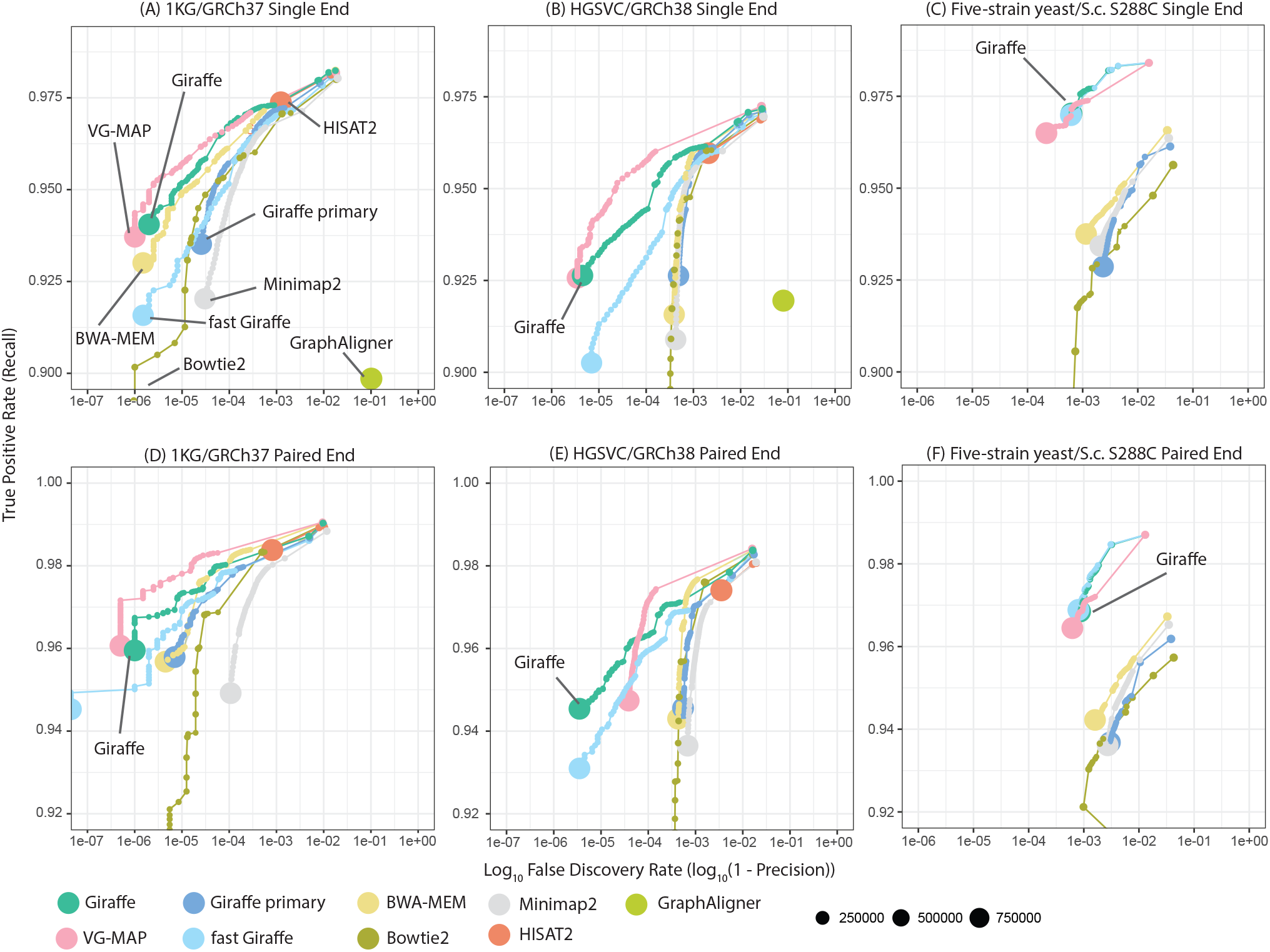
Simulated read mapping. Each panel shows recall vs. *FDR* (false discovery rate, or 1 minus precision) for a simulated read mapping experiment, comparing Giraffe to linear genome mappers (BWA-MEM, Bowtie2, Minimap2) and other genome graph mappers (VG-MAP, GraphAligner, HISAT2). Reads are simulated to match ∼ 150 bp Illumina NovaSeq (for human) or HiSeq 2500 (for yeast) reads, either as single-ended reads (A-C) or as paired-end reads (D-F) (see Methods). Results for each mapper are shown stratified by reported read mapping quality; the size of each point represents the number of reads with the given mapping quality. Three different mapping scenarios are assessed: (A,D) Comparing mapping to a graph derived from the 1KG data to mapping to the linear reference genome assembly upon which it is based (GRCh37). (B,E) Comparing mapping to a graph containing larger structural variations from the HGSVC project to mapping to the GRCh38 assembly upon which it is based. (C,F) Comparing mapping to a multiple-sequence-alignment-derived yeast graph to mapping to the single *S*.*c. S288C* linear reference, for reads from the DBVPG6044 strain. For mapping with Giraffe, we used the full GBWT containing 6 haplotpyes to map to the HGSVC graph and the downsampled GBWT containing 64x coverage of haplotypes to map to the 1KG graph. Giraffe primary are the results of mapping with Giraffe to the linear reference.

### Haplotype sampling improves read mapping

Having rare variants in the graph has been observed to reduce mapping accuracy by creating false positive mappings^22^. While Giraffe’s haplotype mapping approach mitigates issues with high-density variations potentially forming unlikely recombinant paths (Figure 1(A)), it is still possible that rare haplotypes, analogous to or even containing rare variations, could create false positive mappings. Additionally, mapping reads to regions with many distinct local haplotypes can be slow. To overcome these issues, Giraffe includes an optional mechanism for downsampling the haplotypes (see Methods). The mechanism can also create artificial haplotype paths, or *path covers*, in graph components not covered by input haplotypes (for example, decoy sequences or unlocalized contigs) to allow them to be mapped to.

To evaluate the effect of haplotype downsampling on read mapping, we generated GBWT indexes containing different numbers of haplotypes. The *full* 1KG GBWT index contains the 5,036 input haplotypes (real haplotypes from 2,533 samples, which were restricted to the autosomes, sex chromosomes, and mitochondrial genome on which 1000 Genomes performed genotyping) and an additional 16 path covers over connected components not containing input haplotypes. We downsampled these input haplotypes to create GBWT indexes containing between 1 and 128 haplotypes, and used the read mapping simulations described to compare these GBWT indexes to the full GBWT index containing all haplotypes and to a primary graph containing only the paths of the linear reference (Figure 3(A,C)). The mapping benefit of adding more haplotypes was found to plateau at 64 haplotypes, with higher accuracy than is achieved by mapping to the full haplotype set.

**Figure 3.**
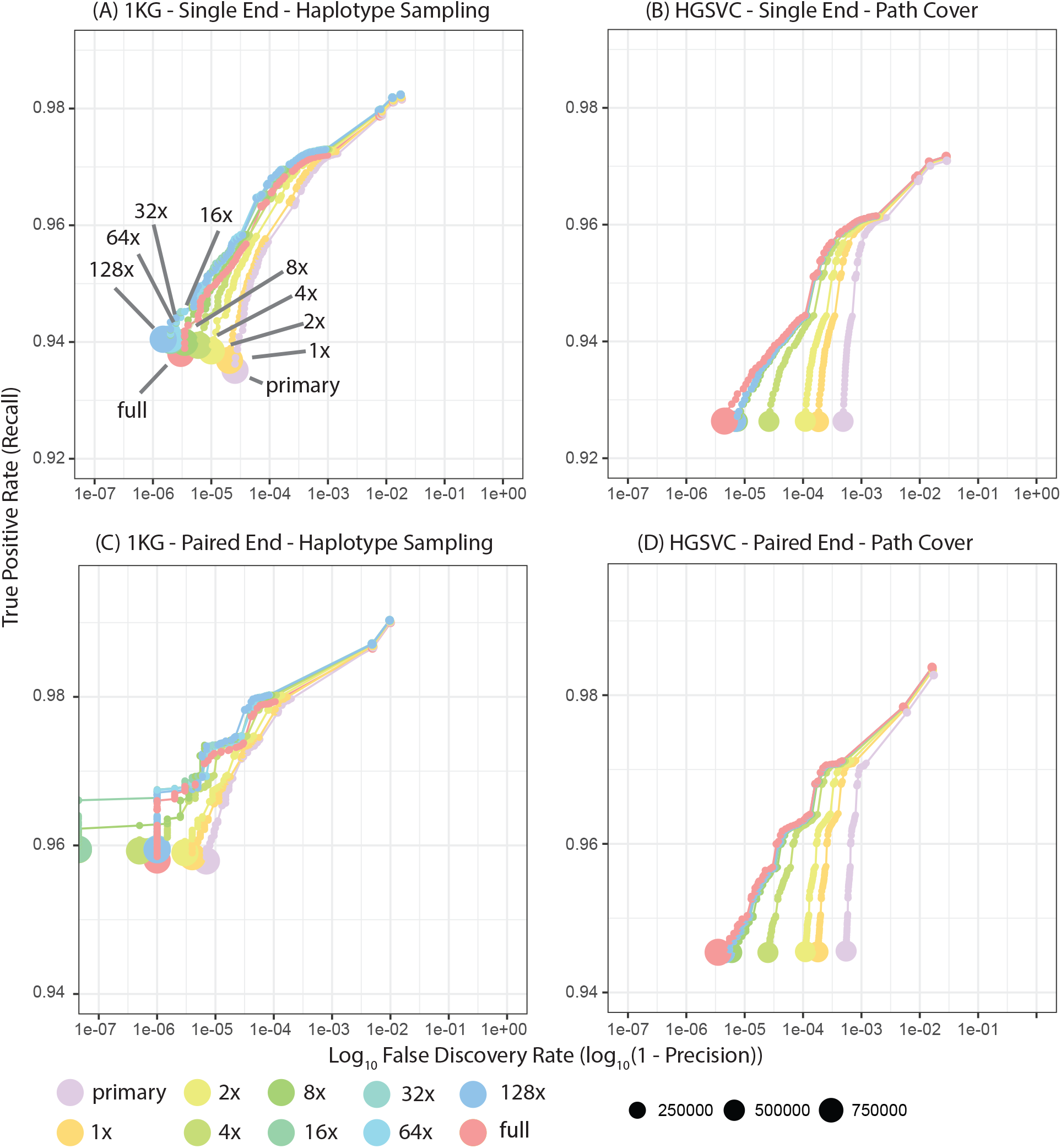
Assessing haplotype sampling and path cover. Each panel shows recall vs. FDR for a simulated read mapping experiment. (A,C) Haplotype sampling, for samplings from 1 to 128 using the 1KG derived graph. Includes for comparison a graph containing just the primary reference (here GRCh37) and, separately, the full 1KG GBWT. For both (A) single-ended and (C) paired-end mapping, performance saturates at around 64x coverage of sampled haplotypes, and exceeds that of mapping to the full GBWT containing all haplotypes. (B,D) Analogous path cover experiments, using the HGSVC graph. All GBWT indexes were evaluated with simulated NovaSeq 6000 reads.

In contrast to the 1KG graph, the full HGSVC GBWT index contains just 6 haplotypes for its three samples. We used this graph for an experiment on generating path covers without known haplotypes, to test how Giraffe, which requires a GBWT, might perform on graphs where there are no haplotypes available. Repeating the same read mapping analysis, we found that path covers alone did not outperform the full underlying haplotype set for the HGSVC graph, but came close to matching its performance (Figure 3(B,D)).

### Giraffe is over 10x faster than VG-MAP

We compared mapping speed and memory usage on an AWS EC2 i3.8xlarge node with 32 vCPUs and 244 GB of memory. To estimate real-world runtime and memory usage, we aligned a complete read set of 600 million NovaSeq 6000 reads from NA19239. We mapped reads to the 1KG graph and, for comparison, the GRCh37 linear reference Figure 4(A,C)) and, similarly to the HGSVC graph and the GRCh38 linear reference (Figure 4(B,D)).

**Figure 4.**
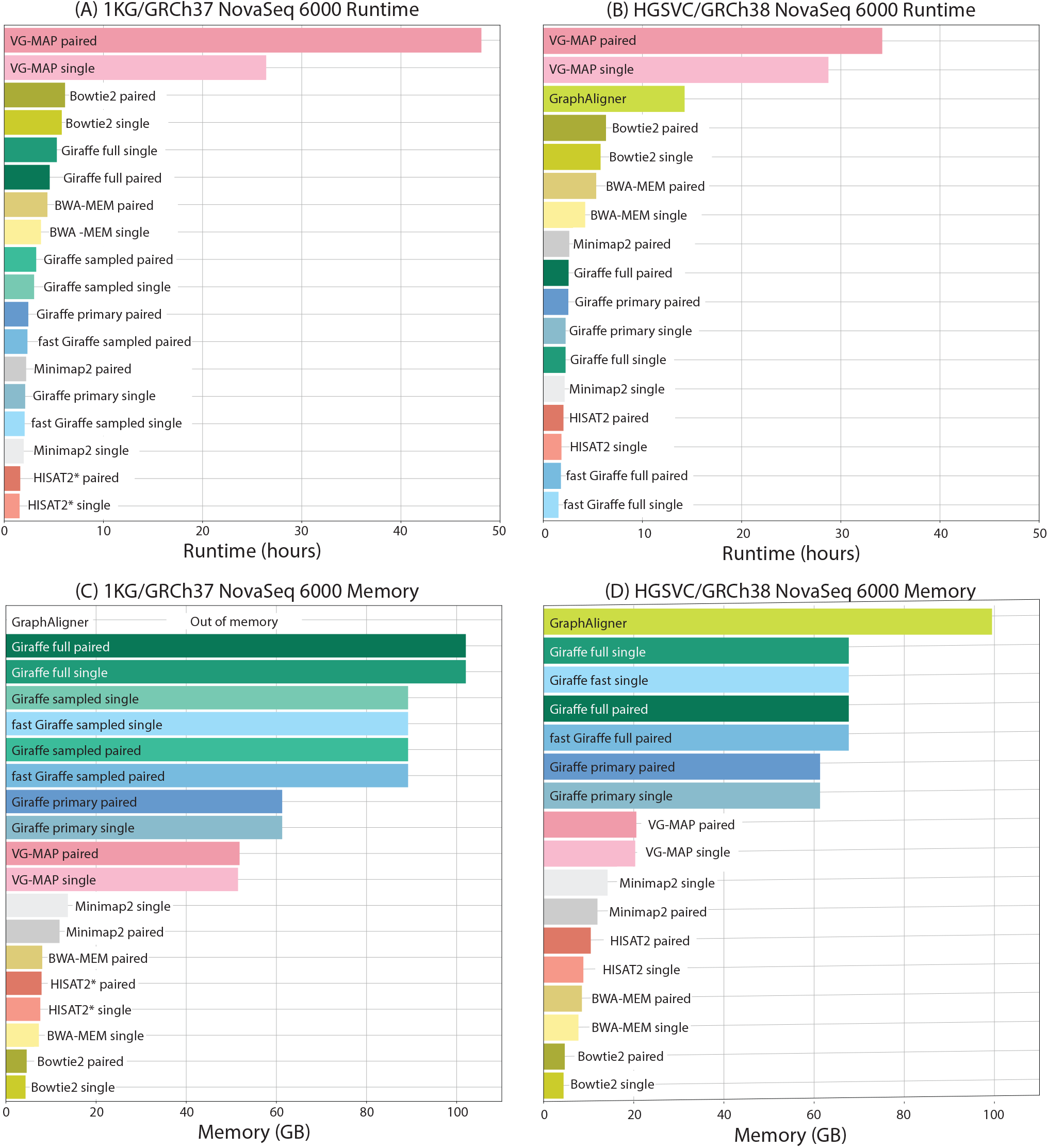
Runtime and memory usage. Total runtime (A,B) and peak memory use (C,D) for mapping ∼ 600 million NovaSeq 6000 reads. (A,C) Reads were mapped to the 1KG derived graph or (for linear mappers) the GRCH37 assembly, and (B, D) to the HGSVC graph or GRCh38 reference, respectively. HISAT2*: results are shown for the author’s reduced 1KG graph.

For each tool we also separately measured reads mapped per thread per second, ignoring the start-up time of the mapper (see Methods). This measure gives an estimate of speed that is invariant to read set size or core/thread count, except for the effects of long-running work batches and thread synchronization overhead. For this evaluation we mapped reads to the 1KG graph and GRCh37 linear reference, to the HGSVC graph and GRCh38 linear reference, and to the yeast graph and the *S*.*c. S288C* linear reference. We also compared mapping reads from other Illumina technologies and got highly consistent results (Supplementary Figures S5, S6).

Giraffe was found to be more than an order of magnitude faster than VG-MAP in all conditions, and was also always faster at aligning to human graphs with a sampled GBWT than Bowtie2 or BWA-MEM were at aligning to the corresponding linear reference. For example, default Giraffe using the 64 haplotype sampled GBWT took 3 hours and 13 minutes to align the complete read set to the 1KG graph while VG-MAP took 48 hours and 6 minutes, BWA-MEM took 3 hours and 42 minutes and Bowtie2 took 6 hours and 8 minutes. Both Minimap2 and HISAT2 were impressively fast, being generally faster than Giraffe in fast mode. For mapping paired-end NovaSeq 6000 reads to the 1KG graph/GRCh37 reference, Giraffe in fast mode using a 64 haplotype sampled GBWT took 2 hours 20 minutes, Minimap2 took 2 hours 12 minutes, and HISAT2 took 1 hour and 37 minutes. GraphAligner, when mapping single-ended reads against the HGSVC graph, was intermediate in speed between VG-MAP and Bowtie2. However, it lacks a paired-end mapping mode, and also could not map the full dataset to the 1KG graph without running out of memory.

Using the 64 haplotype sampled GBWT for mapping, instead of the full ∼ 5, 000 haplotype GBWT, was much faster in every case. For example, the 1KG paired-end mapping task took 4 hours and 35 minutes using the full GBWT, versus 2 hours and 20 minutes using the 64 haplotype sampled GBWT.

Due to the large size of the in-memory indexes it uses, Giraffe’s memory consumption is higher than the other mappers, except for GraphAligner. However, it is able to map to the 1KG graph with the full ∼ 5, 000 haplotype GBWT in ∼ 100 gigabytes of memory—much less than the memory available on most modern cluster nodes (Figure 4(C-D)).

### Giraffe reduces mapping bias relative to BWA-MEM

A major advantage of pangenome mapping is its potential to reduce reference bias: the tendency of linear mappers to align reads to the reference allele over alternate alleles. To assess reference allele mapping bias, we mapped 600 million real paired-end NovaSeq 6000 reads for NA19239 to the 1KG graph using default Giraffe and VG-MAP. For comparison, we mapped the same reads to GRCh37 with BWA-MEM. Replicating a previous approach^23^, we used `bcftools mpileup` and `bcftools call`^24^ to call variants, and filtered to high-confidence variants (root mean square read mapping quality >= 40 and depth >= 25) that were called as heterozygous for all mappers. For these variants, we found the fraction of reads supporting alternate vs. reference alleles at each indel length (Figure 5(A)). Giraffe and VG-MAP both show less bias towards the reference allele than BWA-MEM. This difference becomes more pronounced as indel length increases, particularly for larger insertions.

**Figure 5.**
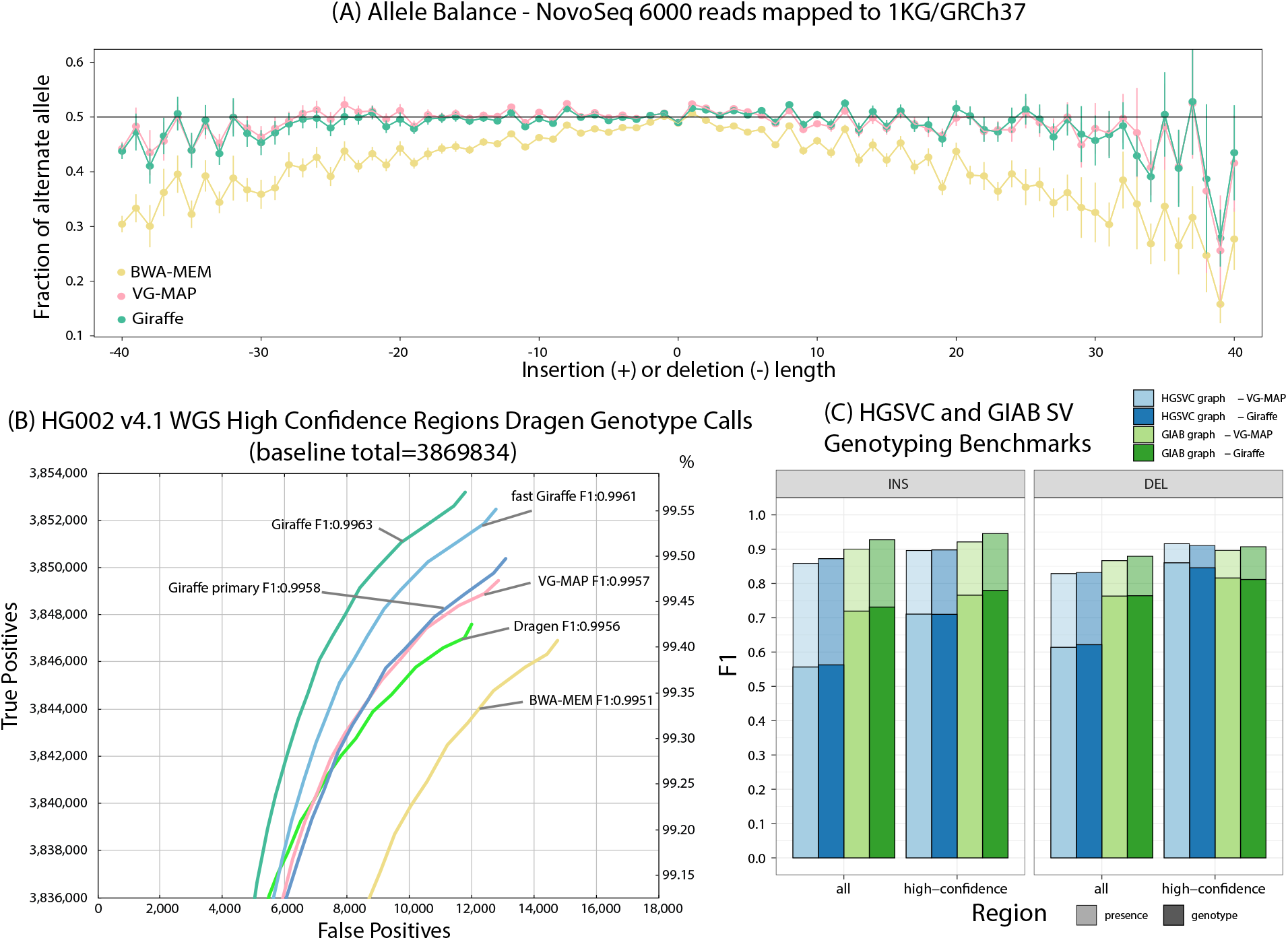
Evaluating Giraffe for genotyping. (A) The fraction of alternate alleles in reads detected for heterozygous variations in NA19239. Reads were mapped to the 1KG graph with Giraffe and VG-MAP and to GRCh37 with BWA-MEM, and the fraction of reads supporting reference or alternate alleles was found for each indel length.(B) Assessing true positive and false positive genotypes made using the Dragen genotyper with projected mappings from Giraffe and other mappers. (C) Comparing Giraffe to VG-MAP for typing large insertions and deletions. ‘Presence’ (lighter bars) evaluates the detection of SVs without regard to genotype, ‘genotype’ (darker bars) requires the SV to be detected and its genotype to agree with the truth genotype.

### Giraffe improves genotyping relative to existing best practices

We compared the performance of Giraffe, VG-MAP, Illumina’s Dragen platform, and BWA-MEM for genotyping SNVs and short indels (Figure 5(B)). To do this, we mapped ∼ 830 million paired-end, 150-bp-long reads (Precision FDA Challenge V2 Illumina ∼ 35x coverage) from the HG002 sample to the 1KG graph. We also separately evaluated mapping ∼ 1.08 billion 250-bp-long reads (∼ 40-50x coverage of Illumina HiSeq 2500) from HG002 to see if using longer reads and higher coverage would affect the results. For the genome graph mappers, we evaluated variant calling performance by using `vg surject` to produce BAM representations of our graph alignments projected onto the linear reference assembly, and then using Dragen version 3.4 to call variants against the `hs37d5` reference for each set of alignments (see Methods). Dragen was used as the primary variant caller because Illumina, who sells it, has found it to produce robust results^25–27^. Its speed was also useful for the purposes of rapid evaluation of whole genome alignments of real read data. No training or calibration was performed for any of the generated mappings other than those performed by default by Dragen itself. The variants produced by these pipelines were then compared against the Genome In a Bottle (GIAB) v4.1 HG002 high confidence variant calling benchmark^28^ using the RealTimeGenomics `vcfeval` tool^29^ and Illumina’s `hap.py` tool^30^ (Figure 5(B); Supplementary Tables S7 and S9). This benchmark set covers 94.1% of the GRCh37 sequence. Out of the examined pipelines, Giraffe produces the highest overall F1 score (harmonic mean of precision and recall) at 0.9963, higher even than Dragen working with the Dragen generated mappings (F1:0.9956) and substantially higher than the BWA-MEM generated mappings (F1:0.9951). Very similar but uniformly higher results were found using the higher coverage, 250bp reads (Supplementary Tables S8 and S10). Somewhat surprisingly, Giraffe with the 150bp read set has a higher F1 score (0.9963) than BWA-MEM (0.9962) with the higher coverage 250bp read set.

To both attempt to further improve genotyping performance and to create a methodology that does not require the Dragen hardware we developed a DeepVariant^31^ pipeline that uses Giraffe mappings (see Methods subsection DeepVariant calling). Using the default DeepVariant 1.0 trained model, we tested genotyping the HG002 sample on chromosome 20 (which was not used in training the model). The DeepVariant/Giraffe pipeline (Chr20 F1: 0.99762) outperforms both Giraffe/Dragen (Chr20 F1: 0.99733) and BWA MEM/Deep Variant (Chr20 F1: 0.99732) (Supplementary Table S11). It is encouraging that Giraffe/DeepVariant outperforms BWA MEM/DeepVariant despite the model not having been trained with Giraffe mappings.

In a previous study, where VG-MAP had been used to map reads to SV pangenomes, we showed that vg performed better than state of the art methods for SV genotyping^6^. We replicated the SV genotyping evaluation on the HGSVC and GIAB datasets^32,33^ to confirm that the quality of the SV genotypes was preserved when using Giraffe (see Methods). We observe similar SV genotyping accuracy across SV types, genomic regions, and datasets (Figure 5C).

### Population Scale Structural Variant Genotyping

Building on our work to type SVs^6^, we demonstrate Giraffe by performing population-scale analysis of an expanded compendium of SVs in large cohorts of samples sequenced with short reads.

#### Building a comprehensive structural variant pangenome from long-read sequencing studies

We combined variants from three catalogs of SVs discovered using long read sequencing: HGSVC^32^, GIAB^33^, and SVPOP^19^. The combined catalog represents 16 samples from diverse human populations and should cover the majority of common insertions and deletions in the human population^19^. An *SV graph* was constructed from this data (note, this SV graph is a distinct graph from the benchmark graphs used above for read mapping and genotyping evaluation) (see Methods). Briefly, near-duplicate variants, i.e. variants present in multiple catalogs with slightly different breakpoints, were first filtered using a re-mapping experiment to keep the version most supported by short reads from diverse samples. We identified 34,010 clusters of near-duplicates SVs using an 80% reciprocal overlap criterion, each with on average 2.6 SVs. For each cluster, we extracted short reads in the region from a diverse group of 17 individuals from the 1000 Genomes Project and HGSVC, pooled them, mapped them to a graph representing the SV cluster, and computed the read support for each variant. A subset of SV clusters didn’t have any SV-supporting reads, because they originally couldn’t be mapped to the linear reference, or were too difficult to map in general, or the variants were rare enough that none of the 17 individuals carried them. Out of the 23,929 clusters with read support, 43.1% had one variant supported by more than 50% of the SV-supporting reads. For each cluster, we kept the best supported variant and filtered out the other near-duplicates. This filtering step removed 55,330 near-duplicate SVs.

While most near-duplicates were removed, many SVs still shared sequence content. To minimize the amount of redundancy in the variation graph, we iteratively aligned and augmented the graph with the SVs in each cluster. We focused on 18,286 clusters of SVs shorter than 1 Kbp, each with an average of 3.24 SVs. A graph was constructed for each cluster by aligning and integrating SV-containing sequences flanked by 5 Kbp of reference sequence, one SV at a time.

The final variation graphs was constructed from 123,785 SVs from the original catalogs: 53,663 deletions and 70,122 insertions. Overall, the variation graph contained 26.2 Mbp of non-reference sequences in the form of insertions. Using a graph decomposition^17^, we identified 228,405 subgraphs representing SV sites and some contributing small variants. These could further be combined into 96,644 non-nested subgraphs, each representing a site not overlapping any other.

#### Efficient genotyping of structural variants across 5,202 samples

Our workflow to genotype SVs using pangenomes is implemented in multiple languages: Workflow Description Language (WDL), Toil, and Snakemake. We first ran the WDL workflow across samples from the Trans-Omics for Precision Medicine (TOPMed) program using the BioData Catalyst ecosystem^34^ (Figure 6(A)). Two thousand samples from the Multi-Ethnic Study of Atherosclerosis (MESA) cohort were selected, using a criterion to maximize the sample diversity (see Methods). The MESA project is a longitudinal cohort study consisting of 6,814 participants at baseline (2000-2002) of white (38%), African-American (28%), Hispanic/Latino (22%) and Asian/Chinese (12%) ancestry, aged 45-84 years at entry, ascertained from 6 sites in the USA. Briefly, the WDL workflow was deposited in Dockstore^35^, TOPMed data was accessed through Gen3, and the analysis was run on Terra. Using the graph described above, it took around 4 days to genotype 2,000 samples from the MESA cohort. In around 6 days, we also genotyped the 3,202 samples from the high-coverage 1000GP dataset, also available on Terra (see Code and Data Availability). On average, genotyping a sample took 194.4 cpu-hours of compute and cost between $1.11 and $1.56 (Figure 6(A) and Supplementary Table S12). The reads were mapped with Giraffe fast mode. The sequencing data was downsampled to ∼ 20x to reduce the computing cost. Downsampling only had a minimal impact on the genotyping accuracy (Figure S8).

**Figure 6.**
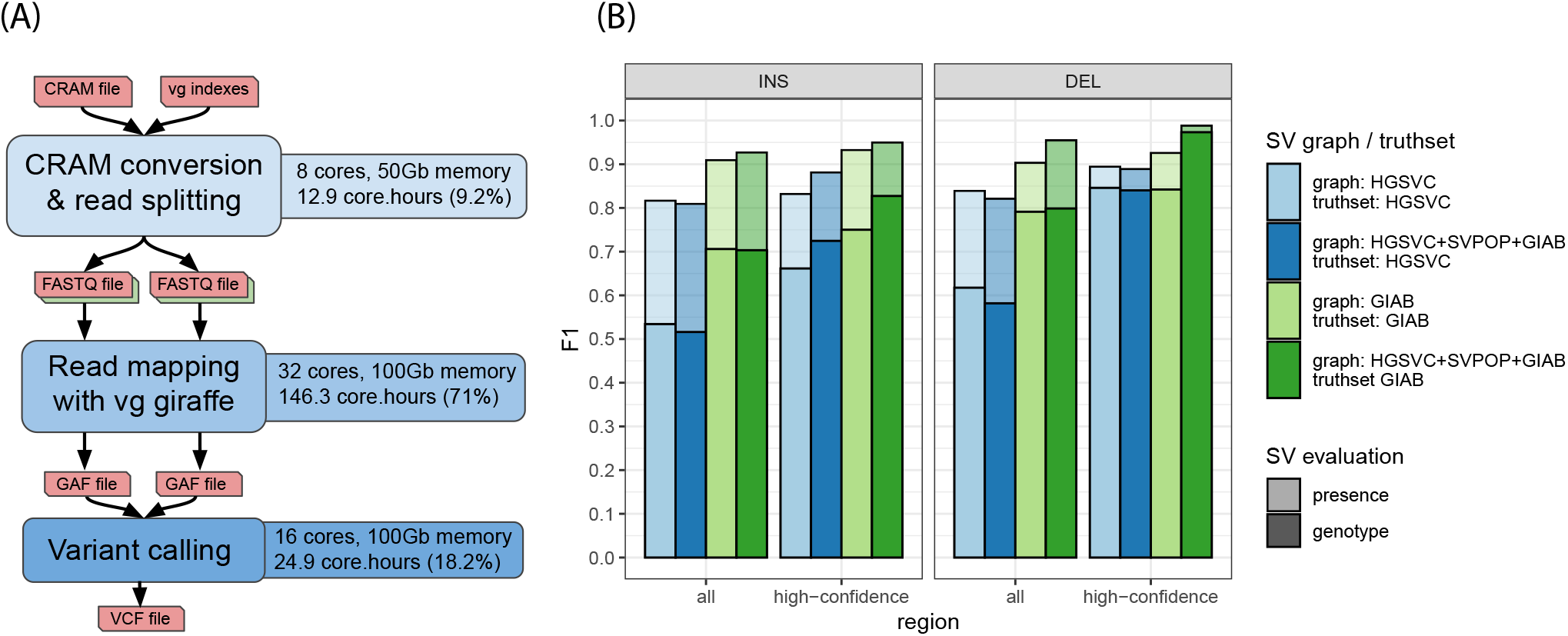
Structural variants genotyping workflow. (A) Workflow overview with average resources used for each task for a typical sample. Note: the core hours estimates include the time spent downloading/uploading files from/to the computing instances (Supplementary Table S13). (B) SV genotyping performance for the SV graph combining HGSVC, SVPOP and GIAB, and individual SV graphs containing only HGSVC or GIAB as used in Hickey et al.^6^.

Compared to Hickey et al.^6^, we used a graph containing more SVs and a more recent version of the vg toolkit, including the new Giraffe mapper described above. For this reason, we reproduced the benchmarking experiment to ensure the quality of the SV genotypes using the HGSVC and GIAB datasets^32,33^ as truth sets. The SV genotypes were as accurate if not more (Figure 6(B)). As before, we evaluated the different SV types across the whole genome or in higher-confidence regions, and both in term of calling and genotyping accuracy (see Methods). Thanks to the new read mapper and improvements in the variant calling approach, the genotyping workflow used about 12 times less resources (core hours) on a sample sequenced at about 20x coverage.

#### Structural variant alleles cluster in sites and span the full spectrum of size and sequence context

Thanks to our construction approach, the SV graph includes the multiple alleles that were cataloged at each SV locus. Although the SV graph was built from 123,785 SVs, novel allelic combinations at SV sites are supported by our genotyping approach. This includes, for example, SNVs and indels within the sequence of an insertion or around a deletion’s breakpoints, or copy-number variations in Variable Number Tandem Repeat (VNTR) regions. The genotyped SVs were also deconstructed into canonical SVs to facilitate their description and grouping into SV sites (see Methods). Hence, the genotyped SVs potentially span more alleles and sites than in the original public SV catalogs. Indeed, we genotyped a total of about 1.7 million alleles clustered in 166,959 SV loci across the 2,000 MESA samples. Figure 7(A-B) shows the size and frequency distributions of the most common allele at each locus. SVs spanned the full size spectrum (50 bp up to 125 Kbp), with 89.4% shorter than 500 bp. For 83.9% of the SVs, simple repeats or low-complexity regions overlapped at least 50% of the SV region. The vast majority of the SV loci (∼ 151 thousand, 90%) contained SV alleles that differed by only small variations from each other (defined as differing by less than 20% when the alleles are aligned (see Methods)), while the rest of the loci showed significant size variation from polymorphic VNTR regions (Figure 7(C) and Supplementary Figures S9(A)). Examples of SV sites that illustrate these different profiles are given in Figure 7(D-E) and Supplementary Figure S10. We observed similar patterns in the 1000 Genomes Project dataset. Genotyping the 2,504 unrelated individuals identified 1.8 million alleles clustered in 166,199 SV loci, of which ∼ 149 thousand (90%) contained SV alleles that differed only by small variations. The size and frequency distribution was very similar to that of the MESA cohort (Supplementary Figure S13).

**Figure 7.**
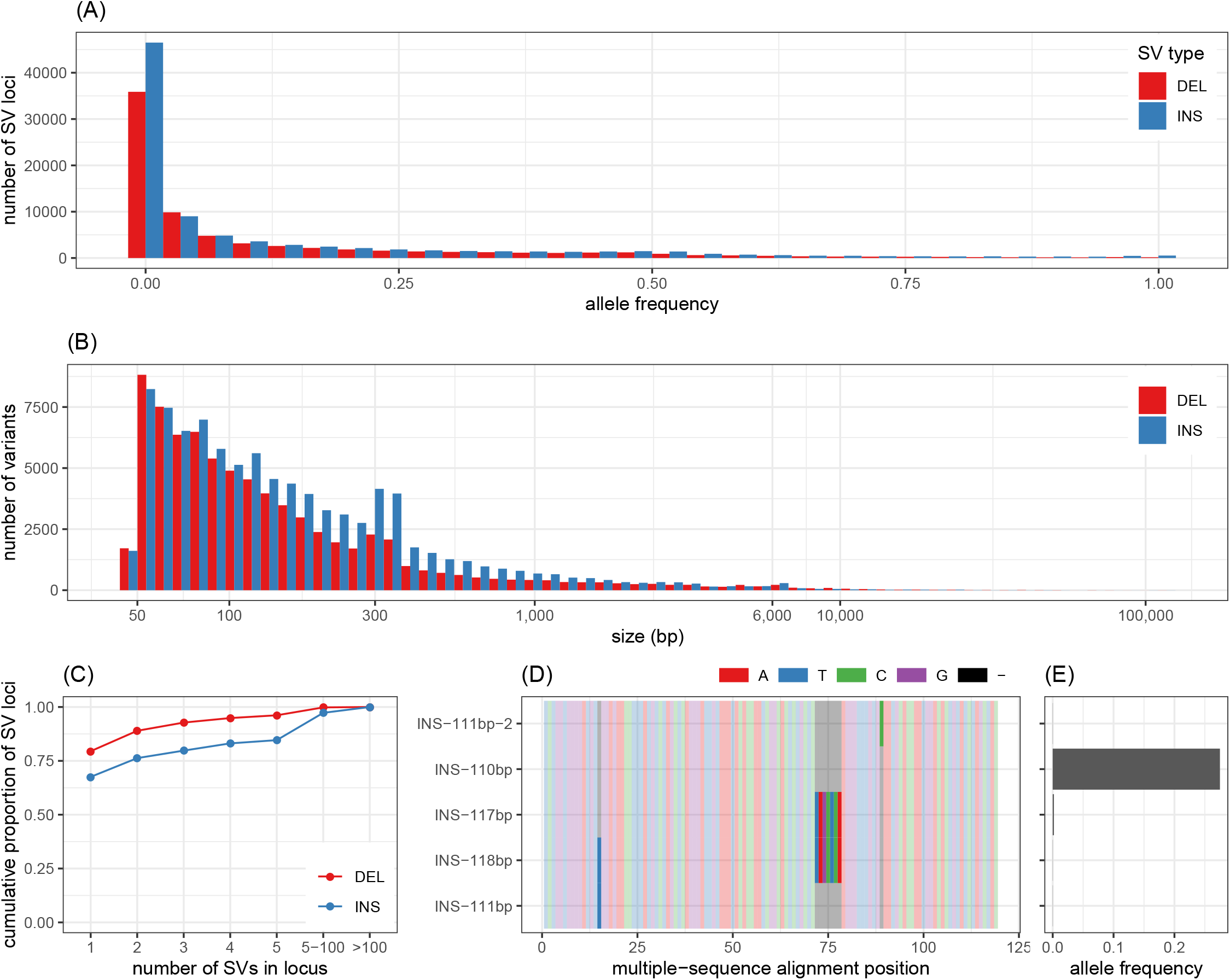
Structural variants in the MESA cohort. (A) Allele frequency distribution of the major allele for each SV loci. Size distribution of the major allele for each SV loci. (C) Cumulative proportion of SV loci depending on the maximum number of alleles (x-axis) in the locus. (D-E) illustrates an insertion site with 5 alleles. The alleles differ by 3 nested indels as shown by the multiple sequence alignment of the inserted sequences represented in (D). Only one allele is frequent in the population as highlighted by (E).

#### Population signatures in structural variant frequencies

Principal component analysis (PCA) of the allele counts at the 166,959 SV sites in the MESA cohort produced a low-dimensional embedding of the samples that appears quite similar to that derived by the TOPMed consortium’s PCA on their SNV genotype data across the entire TOPMed sample set (Pearson correlation: 0.96-0.99 for the top three components, Supplementary Figure S11). Since the SV sites that we genotyped and the sites genotyped by TOPMed come from the same samples which have undergone the same population history, this is the result we would expect to see if our SV genotyper were working properly.

After clustering samples based on PCA, and defining each cluster to be a *population*, we observed that the allele frequencies vary substantially across populations for thousands of SV sites (Supplementary Figure S12(A-C)). For example, we found 21,069 SV sites with strong inter-cluster frequency patterns, as defined by a frequency in any population differing by more than 10% from the median frequency across all populations (Supplementary Figure S12(D)). Using a permutation null model shuffling two-haplotype individuals among populations, we would expect only 85 SV sites with strong inter-cluster patterns (this null model is equivalent to assuming completely random mating for all of human history, although without separating the haplotypes of an individual). The existence of structural variants with significantly different frequencies in different populations supports the need to develop and test genomic tools and references across multiple populations.

Since there is a risk of circularity when using the same genotype data to define populations and look for patterns across them, we replicated these observations in the 1000 Genomes Project dataset, recently sequenced at high-depth (see Code and Data Availability). Here again, the PCA of the allele counts organized the samples in a way consistent with the known history of the 1000 Genomes Project “super population” groups (Figure 8(A-B)). In this analysis, we found 25,960 SV sites with strong inter-super-population frequency patterns, defined as for the MESA analysis, but with the 1000 Genomes super populations as the sample categories (Supplementary Figure S14). Under our permutation null model of completely random mating for all of human history, we would expect only 14 SV sites with strong inter-super-population frequency patterns. More than 17 thousand SV sites with strong inter-super-population frequency patterns were enriched or depleted in the AFR super population, followed by about 10 thousand sites enriched or depleted in the EAS super population. This suggests that people with African or east Asian ancestry might be more likely to be poorly served by genomic references or other tools that do not contemplate these SVs, especially near SV sites where globally-minor alleles are enriched.

**Figure 8.**
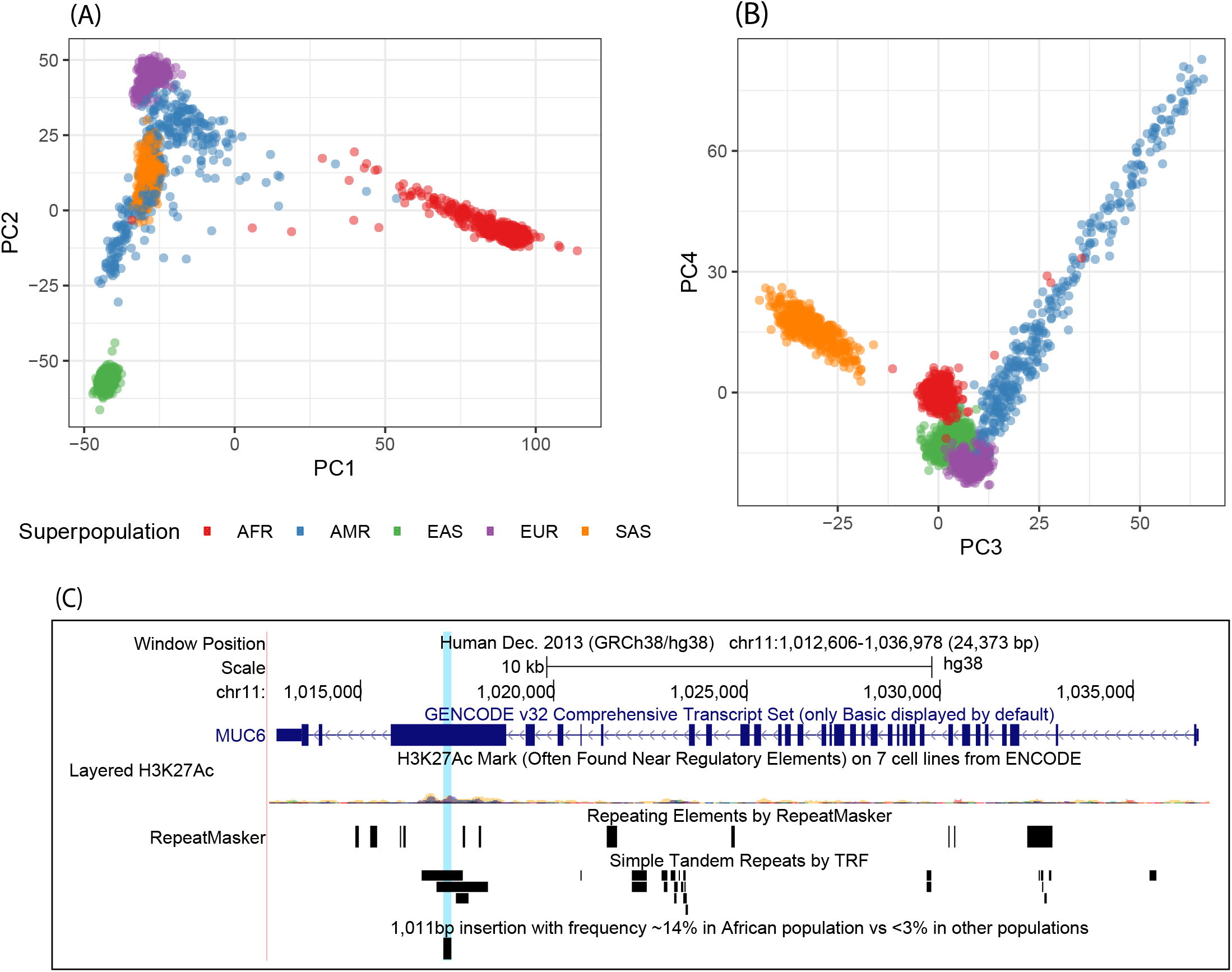
Structural variant genotypes from 2,504 unrelated individuals in the 1000 Genomes Project dataset. (A) shows the first and second principal components derived from the SV genotypes, (B) shows the third and fourth components. Example of an insertion at appreciable frequency (∼ 14%) in the AFR super population while very rare (*<* 3%) in the other super populations. The variant is a 1,011 bp expansion of a VNTR in the coding sequence of the *MUC6* gene.

#### A majority of structural variants are annotated for the first time with comprehensive frequency estimates

The genotyped SVs were originally discovered using long read sequencing technology and many are absent from large-scale SV catalogs. We found that 67-93% of the SV sites genotyped using our pangenome approach are missing from catalogs such as the 1000 Genomes Project SV catalog^36^ or the gnomAD-SV catalog^37^ (Supplementary Table S14). This is consistent with the amount of novel SVs described in the three studies from which our SV graph is derived^19^,32,33. For the first time, our results provide frequency estimates across a large and diverse cohorts for these SVs. Reassuringly, the frequency distribution resembled the allele frequency distributions in the 1000 Genomes Project SV catalog and gnomAD-SV (Supplementary Figure S15(A)). Also, the frequencies of variants present in both our catalog and these public catalogs were largely concordant (Supplementary Figure S15(B-C)). The subset of SVs for which we estimated a higher allele frequency might be due to vg being more sensitive than traditional SV genotypers^6^. Of note, the frequency distribution from SVPOP was clearly different from the other datasets (Supplementary Figure S15(A)). The frequencies of matched variants were also very different between SVPOP and this study or gnomAD-SV (Supplementary Figure S15(D,F)).

As an example of a newly annotated variant, a deletion of the *RAMACL* gene was genotyped with frequency 46.6% in the AFR super population, 4% in AMR, and less than 1% in other super populations. This deletion is not present in the 1000 Genomes Project SV catalog and was unresolved in version 2 of the gnomAD-SV catalog. It has been curated in gnomAD-SV v2.1 and shows similar population patterns, supporting our observation when reanalyzing the 1000 Genomes Project dataset. Such variants could be falsely identified as putatively pathogenic if analyzed only in European-ancestry populations like the EUR super population, where the frequency is low. Our results provide frequency information across an unprecedented number of common SVs in diverse populations that could be used, among other applications, to assist variant prioritization for genomics medicine. Our approach is often capable of genotyping repeat-rich variants such as short tandem repeat (STR) variation. For example, a large 1 Kbp expansion of an exonic STR in *MUC6* with a frequency of 14% in the AFR super population was observed only rarely outside of it: 2.3% in AMR and *<*1% in other super populations (Figure 8(C)). This repeat expansion is absent from gnomAD-SV and the SV catalog from the 1000 Genomes Project. The *MUC6* variable-number tandem repeat (VNTR) region is already known to vary in overall length, with a range of about 15 to 26 subunits of ∼ 507 bp each^38^. One study has associated *MUC6* VNTR lengths at the 66.6th percentile and above with more severe pathology in late-onset Alzheimer’s Disease^38^. This study identified the region as a candidate via a genome-wise scan in an exclusively European-ancestry cohort, from which “ethnic outliers” were removed by a PCA-based process, then followed up by measuring VNTR length in much smaller discovery and replication cohorts of unspecified ancestry^38^. Without catalogs of variants at appreciable frequency in any 1000 Genomes super population, to support large-cohort studies across ancestries, it will be extremely difficult to disentangle genetic and socioeconomic contributions to disease pathology.

#### The frequencies of alleles at a site help fine-tuning structural variants

SVs in the input catalogs may contain errors. When multiple alleles co-occur at a SV locus, we often observed one allele being frequently present in the cohort while other extremely similar alleles weren’t (Figure 7(D-E)). The other alleles at these loci are either extremely rare or they correspond to errors. In any case, it is useful to identify the major alleles and eventually discard the alleles that were not supported in the large-scale sequencing dataset. In 7,346 SV loci, one allele was called in more than 1% of the population while other alleles, present in the original catalogs, were not. Going further, the major allele was at least three times more frequent than the second most frequent allele in 5,936 of these SVs (Supplementary Figures S9(B)). Our results help fine-tune the sequence resolution of these SVs. More generally, our results identify one major allele for 37,667 multi-allelic SV loci.

#### More than 77 thousand structural variants around genes and their association with gene expression

1,561 and 1,599 SVs overlapped coding regions of 408 and 380 protein-coding genes in the MESA cohort and 1000 Genomes Project dataset respectively. In both datasets, these numbers increased to more than 77.8 thousand SVs and 7.6 thousand protein-coding genes when including promoter, intronic and untranslated regions. 10,617 of these SVs had shown strong inter-super-population frequency patterns in the 1000 Genomes Project dataset earlier.

We then tested for association between SVs and gene expression across 445 samples from the 1000 Genomes Project that have been RNA sequenced by the GEUVADIS consortium^39^. These samples span four European-ancestry populations (CEU, FIN, GBR, and TSI), and the Yoruba in Ibadan, Nigeria (YRI) population^39^. A pooled analysis identified 2,059 expression Quantitative Trait Loci (eQTLs) across 909 genes (false discovery rate of 1%, see Methods). 629 of those genes are protein-coding genes. We note that 46% of the SV-eQTLs are located within simple repeats or low-complexity regions. The distribution of the p-values across all tests showed the expected patterns for genome-wide association studies (Supplementary Figure S16). As expected, SV-eQTLs were enriched in coding, intronic, promoter, untranslated and regulatory regions (Supplementary Figure S17). Interestingly, SVs associated with decreased gene expression drove most of the enrichment in coding regions. These results show that the SV genotypes produced here can be used to test for phenotypic association.

Finally, separate analysis of the four European-ancestry populations together, and the YRI population alone, identified respectively 32 and 37 SVs where an association with the expression of protein-coding genes was detected only in the smaller analysis. Possible reasons for this include the 5% frequency cutoff imposed on SV-eQTL candidates (which produced different sets of eligible variants in the different groups), the limited statistical power of the test (as the absence of a finding of an SV-eQTL is not the same as the finding of an absence of an SV-eQTL), and interactions between the candidate SV-eQTLs and the frequencies of other alleles or environmental factors which vary between the populations. As expected, a number of these population-specific SV-eQTLs had shown strong inter-super-population frequency patterns in our earlier analysis: 16 out of the 32 CEU-FIN-GBR-TSI-specific SV-eQTLs and 8 out of the 37 YRI-specific SV-eQTLs showed such patterns.

## Discussion

Pangenome references hold great potential as a replacement for standard linear reference genomes. They are capable of representing variations from a population and have been shown to reduce the bias that arises from using a linear reference. However to date, due to the significant complexity of the task, mapping to pangenomes has been slow or not clearly better relative to linear genomes. Here we introduce Giraffe, a sequence graph mapper capable of mapping to pangenomes containing thousands of aligned haplotypes. Using embedded haplotypes to limit search, we have shown that Giraffe can map to graphs with complex topology as accurately as VG-MAP and as fast as BWA-MEM mapping to a single genome. Furthermore, we have shown for SNVs, indels and SVs that pangenomes can improve genotyping.

For SNVs and short indels our genotying results demonstrate improvements to the state-of-the-art at no additional time or cost, simply by substituting the pangenome for the linear reference during mapping and then projecting back on to a linear reference for the genotyping process itself. Such backward compatible integration provides a path by which pangenomes can be used today without substantially changing downstream tooling.

For SVs, and in particular insertions, we and others have shown the benefits of pangenomes for genotyping are not merely incremental, but transformative^6^,40,41. We developed the most complete SV catalog derived from long-read genomes to date and cheaply and efficiently used it to genotype these variations in over 5,000 short-read, whole genome samples. This approach allowed us to identify duplicate SVs, to refine the canonical definition of SVs and to establish the frequencies of these SVs in diverse human populations. The allele frequencies we established were concordant with previous studies, with the exception of SV-POP^19^, which used an order of magnitude fewer samples and appears to be an outlier. Using a diverse set of short read samples we were also able to reconstruct population substructure from SVs that is consistent with known relationships and those derived from point variations. Analysis of eQTLs show that thousands of the tested SVs have functional impact. We demonstrate further that many of the SVs are differentially distributed across human populations. It is vital in our view that future human pangenomes make the genotyping of these variants accurate and unbiased.

In the near future we expect pangenomes to be built from much larger collections of extremely high-quality long read, *de novo* assembled genomes, such as those being created by the Human Pangenome Reference Consortium. This will provide a human pangenome that will enable comprehensive genotyping of common complex variations, such as SVs, from existing catalogs of short-read sequence samples, putting the typing of such variations on a much more even footing with existing catalogs of variations. We expect the unlocking of this latent information will ultimately reveal new genotype and haplotype to disease associations.

## Methods

### Giraffe Algorithm

#### Definitions

We work in the *sequence graph* model, as defined in the vg toolkit^1^. In a sequence graph, each node *ν* is labeled by a nucleotide sequence (*ν*). A sequence graph is a bidirected graph, which means that each node *ν* has two *sides*: left side *ν*^−^ and right side *ν*^+^. Edges connect unordered pairs of sides of nodes. Nodes can be *visited* in either forward or reverse *orientation*. When we visit a node *ν* in forward orientation 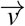, we enter it from the left side *ν*^−^, read the node’s sequence 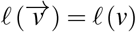, and exit from the right side *ν*^+^. For such a visit, *ν*^−^ is the *entry side*, and *ν*^+^ is the *exit side*. In a reverse orientation visit 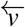, we enter from *ν*^+^, read the sequence 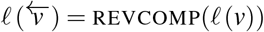, the reverse complement of the node’s sequence, and exit from *ν*^−^. See Supplementary Figure S1 (A) for an example. A *position* in a sequence graph is a pair (*x, i*), where *x* is a node in a specific orientation and *i* is an offset in sequence (*x*). We start counting sequence offsets from 0.

We calculate *alignment scores* with the following parameters: +1 for each matching character, −4 for each mismatch, − 6 for the first character in an insertion or a deletion, 1 for each additional character in the indel, and +5 at each end if the alignment matches or mismatches at that end of the read. Note that, in this scoring model, indels only occur between matches or mismatches; deletions abutting the ends of the read are always removable to get a higher score (and thus never occur), and insertions abutting the ends of the read are actually *softclips*, and are scored as 0 points.

The discussion below describes the general principles used in the Giraffe mapper. We skip many details and special cases.

#### Graph representation

Our graph representation is based on a bidirectional GBWT index^14^. The GBWT is a data structure, based on the FM-index^42^, that can efficiently compress and store a large number of similar paths (haplotypes) in a graph. The primary query in the GBWT is a counting query: as we traverse a path in the graph, we can always tell how many indexed paths contain the path traversed so far as a subpath.

The GBWT data model is based on directed graphs. However, we can accommodate a bidirected graph with a bidirectional GBWT index. For each sequence graph node *v*, we have separate GBWT nodes for visits 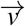 and 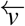. The outgoing edges of a GBWT node correspond to the graph edges on the exit side of that visit, and the incoming edges correspond to the graph edges on the entry side of the visit. See Supplementary Figure S1(B) for an example.

A GBWT index encodes the topology of the graph induced by the indexed haplotypes. This is not always identical to the original sequence graph, as some nodes and edges may not be visited by any haplotype. In order to support the *handle graph* interface^43^, we need some additional structures:

- Determining whether a node with a given identifier exists in the graph is relatively slow with the GBWT interface. Hence, we cache this information in a bitvector.
- We concatenate the sequences for all GBWT nodes (i.e. 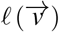 and 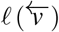 for each sequence graph node *ν*) and store them in a single character array.
- We use an integer array to store references into the concatenation that let us find the beginning of each sequence. When node sequences are requested, we use these references to provide clients with direct access to the stored sequence bytes, without needing to decompress the sequences or create temporary copies.

Accessing edges using the GBWT interface is somewhat slow. We first have to find the record corresponding to the GBWT node and then decompress the header of the record. As seed extension, dynamic programming, and pair rescue repeatedly access the edges in a small subgraph, we can speed them up by caching the decompressed records. We use an application-specific per-thread cache in Giraffe, where eviction consists of discarding the entire cache after each stage of processing for each cluster of aligner seed hits. The cache stores all records we have accessed during the current stage for the current cluster.

#### Downsampling the haplotypes

Mapping to a GBWT with too many haplotypes can be slow in regions with a high density of variations, and can be inaccurate due to rare variants creating false positive mappings. Conversely, a GBWT with too few haplotypes might miss regions that aren’t covered by the haplotypes, especially unplaced or unlocalized contigs, decoy contigs, and other contigs that might not appear in input VCF files. We therefore provide a mechanism for creating artificial haplotype paths for a graph, allowing us to downsample from a large GBWT, augment a sparse one, or create a new GBWT when haplotypes are not available.

We use a greedy algorithm, similar to that used by the Pan-genome Seed Index^44^, for generating paths. We create *n* paths in each weakly connected component and consider the frequencies of subpaths of length *k* = 4 nodes. By default, we use *n* = 64 if we have input haplotypes in any component, and *n* = 16 for graphs without any haplotypes at all. If the component is acyclic, we start from a source node and extend forward. Otherwise, we start from an arbitrary node and extend in both directions, until the length of the path reaches the size of the component.

If there are haplotypes in the component, we choose the extension that best preserves the proportionality of the sampling. Assume we are extending the path forward. Let *X* be the (*k* − 1)-node suffix of the path we are generating, *f*_*h*_(*P*) the frequency of path *P* in the full set of haplotypes, and *f*_*s*_(*P*) the frequency of *P* in the paths we have already generated. We extend the path with node *v* such that *f*_*h*_(*Xv*)*/*(*f*_*s*_(*Xv*) + 1) is maximized. If there are no haplotypes in the component, we assume that each path is equally likely and choose the extension with the lowest coverage so far.

#### Minimizer seeds

Giraffe maps reads using the common seed-and-extend approach. We use a minimizer index^15^ for finding seeds. We define a (*w, k*)-*minimizer* as the *k*-mer with the smallest hash value in a window of *w* consecutive *k*-mers and their reverse complements (in a *k* + *w* − 1 bp window). As we expect that sequencing errors are rare and that common variants are present in the haplotypes, we use longer than usual minimizers: *k* = 29 and *w* = 11 by default.

Our minimizer index is a hash table with 64-bit keys and 128-bit values. 62 bits of the key are used for encoding the *k*-mer, allowing us to support *k* ≤ 31 bp seeds. The value is either a single *hit* or a pointer to a sorted array of hits. We use the high bit in the key for indicating the type of the value. For each hit, we use 64 bits for the graph position (53 bits for node identifier, 1 bit for node orientation, and 10 bits for sequence offset) and reserve the other 64 bits for *payload*.

We index only unambiguous minimizers, skipping *k*-mers containing characters that represent multiple nucleotides. We only consider paths consistent with the haplotypes. Index construction is fast: typically 5–10 minutes for a human genome graph using 16 threads.

For mapping, we prioritize the most informative minimizers (those with the smallest number of hits) and always select more-informative minimizers before less-informative ones. For a minimizer with *n* hits, we define *minimizer score* as max(1 + ln*C*_*h*_ − ln *n*, 1), where *C*_*h*_ is the *hard hit cap* (default 500 hits). We try to select enough minimizers so that the sum of minimizer scores for the selected minimizers reaches the *score fraction* (default 0.9) of the total score for all minimizers available, but we never select any minimizers with more hits than the hard hit cap. To do this, we first select all minimizers that have a number of hits at or below the *soft hit cap C*_*s*_ (default 10 hits). Then, if we have not yet achieved the score fraction, we select progressively less-informative minimizers one by one. We stop when we reach the score fraction, run out of minimizers, or the next minimizer has more than *C*_*h*_ hits.

#### Minimum distance clustering

In order to avoid redundant work, we cluster the seeds into sets that should correspond to the same mapping. Two seeds are assigned to the same cluster if the minimum distance between them is sufficiently small. It must be no larger than the *distance limit* (default 200 bp), or, for longer reads, |*R*| + 50 bp, where |*R*| is the read length. ^16^

As computing distances between graph positions can be slow, we use a minimum distance index to speed up the clustering. The index is based on the *snarl decomposition* of the graph. A *snarl*^17^ *is a graph-aware concept of a genetic locus, or a “site”. It is a subgraph separated by two node sides from the rest of the graph. A chain* is a series of snarls arranged end to end, like beads on a string, with no connections to anything else except (possibly) at the ends. Any sequence graph can be recursively decomposed into a tree of snarls containing chains, and chains containing snarls. In order to find the distance between two graph positions, we determine their lowest common ancestor in this *snarl tree* and calculate the distance by traversing the tree and consulting precomputed tables at each level.

Non-informative seeds are often scattered around the graph. Finding them in the snarl tree causes cache misses for each seed. However, in a typical sequence graph, most positions are located close to the root of the snarl tree. For such positions, the amount of information we get from the distance index is small. Hence, we can cache the distance information for seeds near the root in the payload field of the minimizer index.

After clustering, we score the clusters by two criteria and pick the highest-scoring clusters for extension. The primary criterion is *read coverage*: the fraction of the read covered by the seeds. The secondary criterion, for breaking ties, is *cluster score*: the sum of minimizer scores for the minimizers present in the cluster.

#### Gapless haplotype-consistent seed extension

We need to turn seeds into *extensions*: gapless, haplotype-consistent partial alignments that may be longer than a minimizer and may contain some mismatches. We define a *mismatch bound* (default *d* = 4) that limits the number of mismatches in various parts of the extension.

Before extending, we first merge redundant seeds. A set of seeds is redundant if they all correspond to the same gapless alignment between the read and a node.

We then try to extend the remaining seeds into haplotype-consistent alignments without gaps. We extend up to 800 clusters (*extension bound*) but ignore those with read coverage or cluster score a certain threshold below the best cluster. For each cluster, we return a set of extensions. We also determine the *extension set score* for the cluster as an upper bound for the alignment score, assuming that we can chain the extensions arbitrarily over the read while ignoring graph topology.

To find the extensions, for each merged seed, we first gaplessly align the read to the entire node corresponding to the seed, and place that extension into a priority queue by alignment score. Then we process the extensions in the priority queue, highest-scoring first.

We define a *right extension* of an extension as the original extension, made longer on its right side in a haplotype-consistent way by extending the gapless alignment into another node in the graph. A right extension of an extension matches some nonempty subset of the haplotypes that match the original extension. Each haplotype in the original extension either continues on into some right extension, or ends. An extension is *right-maximal* if i) we have reached the end of the read; ii) it does not have right extensions for all haplotypes that currently match it; or iii) a right extension would have both more than *d* mismatches and more than *d/*2 mismatches in the right flank after the initial node. *Left extensions* and *left-maximality* are defined similarly. If the highest-scoring extension has not been found right-maximal, we find all its right extensions and place them in the priority queue. If, while attempting this process, we find that the extension is right-maximal, we mark it as such and return it to the priority queue. If the highest-scoring extension *has* been found right-maximal, we extend it to the left as above, marking it left-maximal if we find it to be so. At the end, we report the highest-scoring, both left-maximal and right-maximal extension we find for the seed.

We then look at the collection of highest-scoring, left-and-right-maximal extensions across all seeds. If a full-length alignment with at most *d* mismatches exists as an extension of a seed, we will have found it. In that case, we sort all the full-length alignments we have found in descending order by alignment score. Then we greedily select all alignments that do not overlap with any of the already selected alignments, where we consider two alignments *overlapping* if the fraction of (read, reference) base mapping pairs in common is greater than the *overlap threshold* (default 0.8). If there are no full-length alignments with at most *d* mismatches, on the other hand, we trim the partial extensions to maximize the alignment score and return the whole set of distinct extensions.

#### From extension sets to alignments

We transform at least two, but no more than 8 (the *alignment bound*) of the highest-scoring extension sets into alignments, and ignore those with scores further than the *extension set score threshold* (default 20 points) below that of the best set.

We try to find at least two alignments from each extension set we use. If an extension set consists exclusively of full-length gapless alignments, we can use them directly. We take the top two if there are two or more, or the only one otherwise. If there are partial extensions in the set, we also consider alignments computed from the best partial extensions in the set, to see if they beat out any full-length gapless alignments for those top two spots. Using each partial extension as a seed, we perform dynamic programming alignment of partial extensions above a score threshold, which is set dynamically such that we align at least one partial extension and end up with at least two alignments overall (including the full-length gapless ones).

Once these criteria are met, we do not align further partial extensions, unless we estimate that the extension could produce a better alignment than the best we have already seen in the extension set. We estimate the best possible alignment score for extension *X* by considering three types of alignments for each unaligned tail, without taking the graph into account:

1. An exact match of at most *k* + *w* − 2 bp (shorter than the minimizer window) followed by a gap to *X*.
2. A gap to another extension *Y*, an exact match up to the first mismatch in *Y*, and then another gap to *X*.
3. A gap to another extension *Y*, then *Y* or a part of it, and then another gap to *X*. In this case, we assume that all mismatches in *Y* are in the part we use.

Then we use the highest score across all cases as the estimate.

To actually perform dynamic programming alignment for an extension, we start from the GBWT search state for the extension. The search state corresponds to a set of haplotypes. For each unaligned tail of the read outside the partial extension, we build a tree-shaped graph by tracing out these haplotypes in the appropriate direction, splitting whenever two or more haplotypes in the set diverge from each other. We then align the tail to the tree using the dozeu library https://github.com/ocxtal/dozeu, a vectorized implementation of X-drop alignment.

After collecting up to two alignments from each extension set, we select the highest-scoring alignment as the *primary* for the read, while all others are *secondaries*.

#### Paired-end mapping

Paired-end mapping runs start with fragment length estimation, if a fragment length distribution has not been specified. We map the reads in single-ended mode using a single thread until we have 1000 uniquely mapping read pairs. We then estimate the mean and standard deviation of fragment length using the uniquely mapping pairs (see Fragment length estimation).

Once we have estimated the fragment length distribution, we continue mapping with multiple threads. We cluster the seeds for both reads in the pair at the same time and try to group clusters that are at the right distance from each other. Each *cluster group* can contain clusters from either mate pair that are closer than the *fragment distance limit* to each other (default: fragment length distribution mean + 2 standard deviations). We produce alignments within each cluster as is done for single-ended mapping, prioritizing cluster groups containing clusters from both reads. We then attempt to find pairs of alignments that originated from the same cluster group. Pairs of alignments are scored as the sum of the two alignment scores and the log likelihood of their fragment length. If we do not find paired alignments or if we have an unpaired alignment with score at least 0.9 times (the *rescue threshold*) the score of the best alignment, we try to *rescue* the pair. We attempt to rescue at most 15 unpaired alignments (the *rescue bound*).

To perform rescue, we use an aligned read to find an alignment for its unaligned mate. We use the distance index to find the *rescue subgraph* with all nodes within at most 4 standard deviations of the expected read location. We then find all minimizer hits in the subgraph, including those we ignored during mapping. Because the hits are sorted by node identifier in the index, we can find the ones in the rescue subgraph efficiently by doing list intersection with exponential search. We treat all the seeds as a single cluster and try to extend them without gaps. If we find a full-length alignment, we use it as the rescued alignment.

If there are no full-length extensions, we resort to dynamic programming. We now consider alignments to the entire subgraph, not just to the paths consistent with haplotypes. By default, we use the dozeu algorithm. If we had minimizer hits in the rescue subgraph, we use the best extension as the seed for dynamic programming. Otherwise, we find the best alignment for the first 15 bp of the read and use it as the seed. As a slower but more accurate alternative to dozeu, we also support using the GSSW alignment implementation^45^. While dozeu must extend a seed, GSSW always finds the best alignment in the subgraph.

#### Fragment length estimation

We use a method of moments approach to estimate the fragment length distribution The distribution is modeled as a normal distribution. To estimate the parameters of the distribution, we map pairs of reads without pairing constraints until obtaining *N* uniquely mapped pairs that can reach each other through some path in the graph (default *N* = 1000). To avoid producing a fragment length distribution that does not reflect expected lengths from paired-end sequencing machines, we do not consider distances that are greater than the *maximum fragment length* (default 2000 bp). We discard the largest and smallest (1 − *γ*)*/*2 distances that we find (default *γ* = 0.95). We use the remaining *γN* distance measurements as a sample from a truncated normal distribution. The truncated distribution has the same *µ* and *σ* ^2^ as the underlying normal distribution, and the truncation points can be calculated based on *µ, σ* ^2^, and *γ*. We can estimate the following parameters using the method of moments:

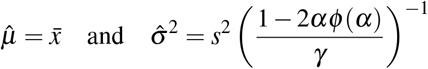

In the above, 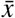 and *s*^2^ are the maximum likelihood estimates of the mean and variance among the retained *N* measurements, and *α* = Φ^−1^((1 − *γ*)*/*2) is the left truncation point on a standardized normal distribution.

#### Mapping quality estimation

Mapping uncertainty is conventionally summarized as a *mapping quality*: the Phred-scaled^46^ posterior probability of a mapping error. To compute mapping qualities, we take advantage of the interpretation of an alignment score as the log-likelihood of a Viterbi path through a pair hidden Markov model^47^. With this model, the posterior probability of the maximum likelihood alignment *Â* of a query sequence *Q* to the reference *R* can be expressed as

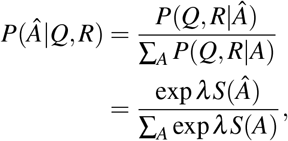

where *S* is the alignment score, and *λ* is a normalizing factor that can be numerically computed from the scoring parameters^48^. In practice, we do not score all possible alignments in order to evaluate this expression’s denominator. Rather, we approximate it by using the high-scoring secondary alignments that are discovered during the mapping process.

#### Mapping quality caps

We impose several *caps* on the mapping quality derived from the formula above, each of which models a source of error due to Giraffe’s heuristics. All mapping qualities are capped at 60 to account for some miscalibration and approximation in the mapping quality model. Another cap is computed based on the probability that all of the minimizers we explored were actually created by base errors. Finally, we employ additional mapping quality caps for paired-end mapping to account for uncertain pairing and error introduced by hard caps in the rescue algorithm.

#### Minimizer error mapping quality cap

The Giraffe algorithm can only find a correct mapping if the read contains instances of minimizers that exactly match minimizers in the true placement in the graph, which then form a cluster, which is then extended to produce an alignment. Consequently, the probability that all the minimizer instances in extended clusters were actually created by errors in the read is an upper bound on the mapping quality. We take a conservative approach to estimating this cap where we actually compute a lower bound on the probability of error. In particular, we make assumptions that restrict the set of errors we consider for the sake of computational efficiency. We consider only substitution errors, which are much more frequent than indel errors, and we restrict our attention to only the most probable way the minimizer instances could have been created, rather all possible ways.

Each minimizer instance in the read is minimal in one or more windows; we call the consecutive run of windows that produce a minimizer instance the minimizer instance’s *agglomeration*. The agglomeration can be divided into the minimizer itself, of length *k*, and the *flank* of zero or more bases beyond the minimizer on either or both ends. In order for all minimizer instances to be disrupted, it is necessary for each window in each agglomeration to contain at least one error. Mathematically, we say that for an agglomeration *i*, the minimizer occurs at bases in the set core(*i*) = {minstart(*i*) *…* minend(*i*)} in the read, and the agglomeration occurs at bases {aggstart(*i*) *…* aggend(*i*)}. We write the hash value of the agglomeration’s minimizer as hash(*i*).

We consider only the subset of minimizer instances that were explored, and thus contributed to the set of candidate alignments. Our goal is to find a lower bound on probability of disrupting all minimizers, via errors in their agglomerations. We take advantage of the fact that we are looking for a lower bound by restricting our considered cases to those where each base error disrupts all windows that it overlaps. We also employ an approximation, by treating as sufficient the disruption of any windows of an agglomeration with an error in the flank, instead of requiring errors in the flank to disrupt all windows of the agglomeration.

We consider the minimizers’ agglomerations to be sorted in the order they appear along the read. We approach this as a dynamic programming problem, with the dynamic programming table *c*, where *c*_*i*+1_ holds a lower bound on the probability that all agglomerations {0 *… i*} were disrupted. We use *I* to represent the number of the final agglomeration, and we number the bases in the read (or *columns*) as *j* running {0 *… J* .}

We imagine all the agglomerations arranged under the read, each on its own line (Figure 9(B)). We overlay the minimizers with *regions* left to right, where the region changes whenever an agglomeration begins or ends (Figure 9(C)). We number these regions as *r* running {0 *… R*}, and define their 0-based end-inclusive bounds in agglomerations as {top(*r*) *…* bottom(*r*)} and in columns as {left(*r*) *…* right(*r*)}.

**Figure 9.**
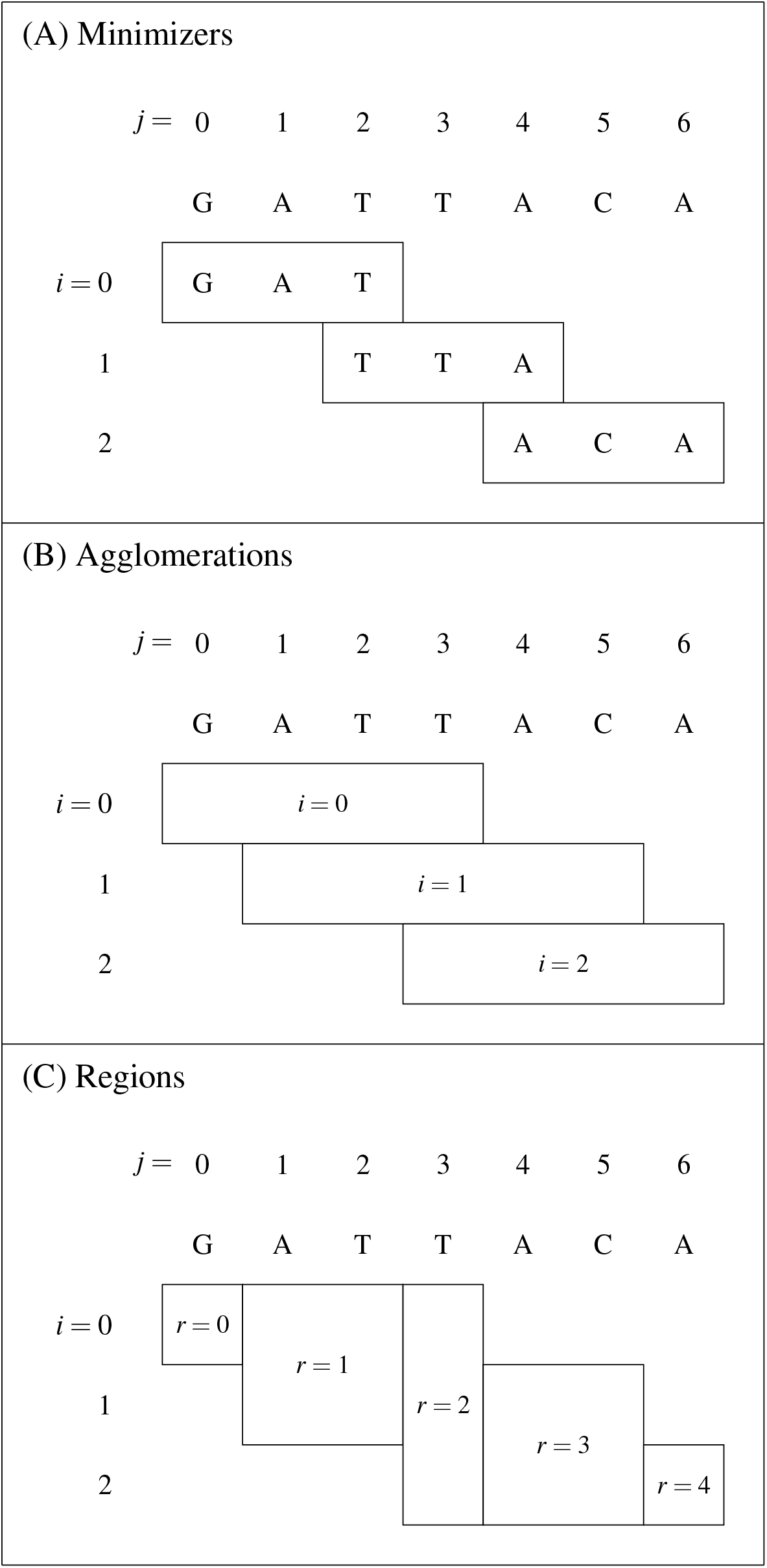
Diagram of minimizers, agglomerations, and regions. In this example, we use a minimizer length of *k* = 3, and a window size of *w* = 4. (A) shows where three example minimizers fall in the read. (B) shows the agglomerations of these minimizers, if the minimizers happen to be minimal for all windows they appear in. For agglomeration *i* = 1, we have aggstart(1) = 1, aggend(1) = 5, minstart(1) = 2, and minend(1) = 4. (C) shows the regions, which we use to decompose the part of the diagram covered by agglomerations, with breaks wherever an agglomeration begins or ends. For the region *r* = 3, we have top(3) = 1, bottom(3) = 2, left(3) = 4, and right(3) = 5.

We define the following events to structure our probability model:

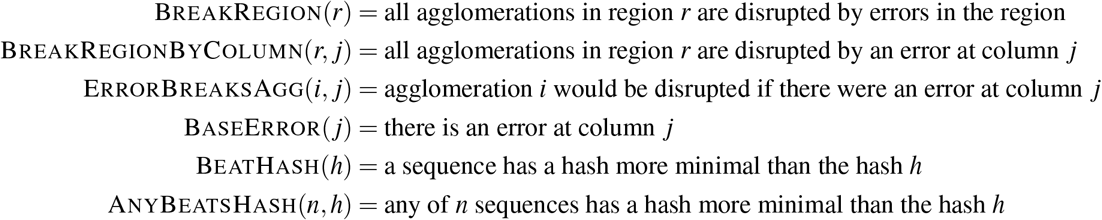

We implement the following recurrence relation on the dynamic programming table *c*:

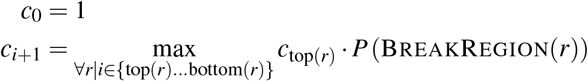

In addition to the minimizer and region information, we take the read quality string *q* as input, which defines *P* (BaseError(*j*)). Also, we approximate the output of hash(*i*) as a uniform random integer in {0, 1, *…*, 2^64^ − 1}. We can then evaluate the recurrence to fill the dynamic programming table using the following relationships:

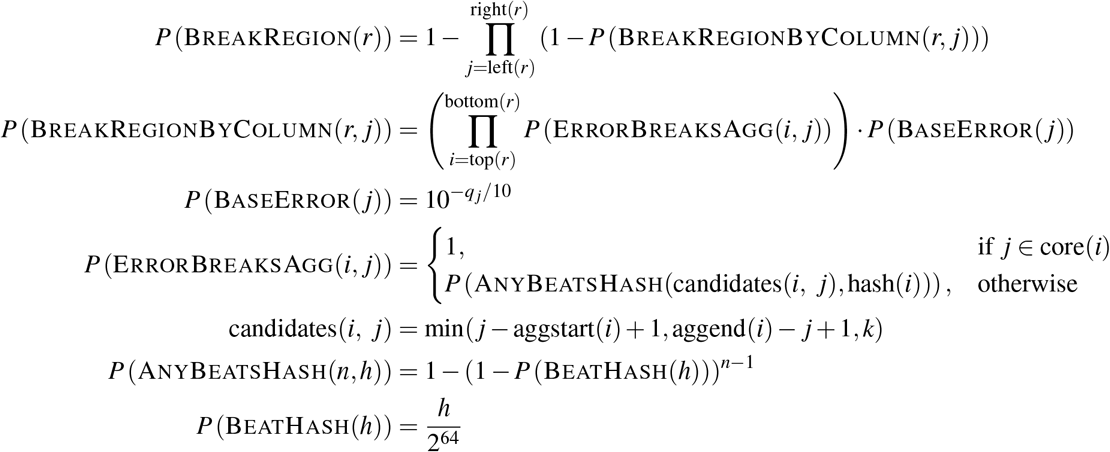

The final result is the bottom cell in the dynamic programming table, at *c*_*I*+1_: the probability of the highest-probability case in which all agglomerations in the region are disrupted. This probability is converted from the internal log10-probability representation used for computation into a Phred score.

The Phred score is then transformed by an empirically-developed heuristic to produce a mapping quality cap, based on how the computation of the initial mapping quality went for the read or read pair. The initial mapping quality can be *pegged* at the maximum value representable by a 32-bit signed integer, which occurs when the score gap between the top-scoring alignment or pair and any secondary is too great. Our heuristic is that, when the initial mapping quality is not pegged, the cap is used as-is, but if the initial mapping quality for the read is pegged to its maximum representable value, the cap is doubled.

#### Paired read mapping quality caps

We apply two different caps for paired reads based on the outcome of the pair rescue routine. If we didn’t find the best pair of alignments with rescue, we cap the mapping quality of each individual alignment to account for uncertain pairing. First, we find all alignments for the one read that were in a pair with the same alignment of the other read. We then calculate the mapping quality of the best pair based on the scores of these alignments and use the resulting value as a cap on the mapping quality of the other read. As a second cap, we treat all equally-scoring cluster groups as having an equal probability of producing a correct pair of alignments. Therefore, for the final alignment pair, we take the probability that we picked the incorrect alignment pair to be at most 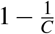, where *C* is the number of equivalent or better cluster groups, and cap the mapping quality accordingly.

If all pairs of alignments were found by rescue, then we adjust the mapping quality to model the effects of Giraffe’s hard limits on the number of rescue attempts. For the primary alignment pair, we estimate the number of equivalent pairs that would have been found without the limit. This estimate is the total number of unpaired alignments divided by the number of alignments that were actually rescued. When calculating the mapping quality of the primary alignment pair, we compute it as if the estimated number of pairs had been found.

### Mapping evaluation

We ran each of the mappers on a variety of different graphs and read sets to evaluate their accuracy (Figures 2 and 3 and Supplementary Figures S2, S3, and S4), speed (Supplementary Figure S5), and time and memory consumption (Figure 4).

#### Graph construction

Human graphs were constructed using the toil-vg tool, versions 71f484c through cfe1599. Construction and indexing required multiple retries of the workflow with successive improved versions of the tool. Some indexes, such as the multiple kinds of sampled GBWT indexes and their corresponding minimizer indexes, were generated with additional ad-hoc scripts around functionality in vg. To run GraphAligner, we also needed to convert our vg-format graphs to GFA format using vg.

The indexes used by HISAT2 for the mapping evaluation were constructed using version 2.2.0. Variants were first split into bi-allelic sites and normalized using bcftools version 1.9^24^. The variants were then preprocessed using `hisat2_extract_snps_haplotypes_VCF.py` (part of the HISAT2 package) with the option that allows variant ids not beginning with rs. Furthermore, `Y` bases in the reference genome were changed to `N` for the preprocessing only. The resulting variant and haplotype list was then used as input to `hisat2-build` to construct the index.

For the 1KG graph, VCF files from the 20130502 release were used, with sample genotypes included. Because we used reads simulated from NA19239 to evaluate mapping accuracy in the 1KG graph, we did not include any variants that were unique to NA19239 or their child NA19240 in the 1KG graph. These samples and their private variants were filtered out of the input VCF files before the graph was constructed. The other parent of the family, NA19238, was retained. This left the other 2,533 samples that were present in at least one of the VCFs to be included the base haplotype index.

For the HGSVC graph, VCF files from the HGSVC were used (see Hickey et al.^6^ and available here), together with a FASTA reference combining the GRCh38 no-alt analysis set (accession GCA_000001405.150) and the `hs38d1` decoy sequences (accession GCA_000786075.2). Although NA19240 was used as the simulation target for simulating reads, no haplotype data or private variants were held out, as the graph contained only three samples to begin with.

The same reference genomes and variant sets as above were used to build the HISAT2 indexes, except for the GRCh37 1KG set. HISAT2 was not able to create an index using this set specifically, likely due to its size and complexity. We tried to reduce the set to only variants with a frequency at or above 0.01, but this did not help. We therefore instead used their pre-made variant containing GRCh37 index, which is available at https://genome-idx.s3.amazonaws.com/hisat/grch37_snp.tar.gz. This index contains the common SNVs and indels from dbSNP 144, which includes the variants from the 1000 genomes project.

For the yeast experiments, a five-strain yeast multiple sequence alignment previously constructed^6^ was imported as a graph with `hal2vg` v2.1^6^, with ancestral genomes (internal nodes in the alignment tree) excluded.

#### Sample selection

For human evaluation, we decided to use real and simulated reads from NA19239 and NA19240. These two people are parent and child from a well-studied trio pedigree, sometimes described as “the Yoruba trio”, recruited from among people claiming four Yoruba grandparents in Ibadan, Nigeria^49,50^. After obtaining informed consent and collecting blood, the original collecting investigators gave donors “an equivalent of US $8.00 and multivitamins worth ∼ US $4.00 to compensate them for their time and travel”^51^. Because NA19239 had haplotypes available on the 1000 Genomes variants, and NA19340 had haplotypes available on the HGSVC variants (no sample has both), and real reads from a variety of Illumina instruments were publicly available for one or the other, these two samples were chosen to avoid unduly confounding graph choice and sample genome.

#### Real reads

For speed evaluation on human data, reads from accessions ERR3239454, ERR309934, and SRR6691663 were used. The reads were downloaded, shuffled as pairs, and subset to a set of 500 thousand pairs (1 million reads) and a set of 300 million pairs (600 million reads).

For speed evaluation on yeast data, real reads for each strain were used, from accessions SRR4074256, SRR4074257, SRR4074394, SRR4074384, SRR4074413, SRR4074358, and SRR4074383, shuffled and subset to 500 thousand pairs as with the human data.

#### Read simulation

To train vg’s read simulator for generating human reads, the truncated 500 thousand pair real read sets were used to train the simulator to match the read length and quality string characteristics of each of the three different Illumina instruments. Training reads for human were interpreted as pairs. To train the read simulator for the yeast experiments, the Illumina HiSeq 2500 reads from accession SRR4074257 were used as the training reads. These training reads were not interpreted as pairs, and the entire file was used.

For the human 1KG experiments, using `toil-vg construct`, the input VCFs were subset to just sample NA19239, and a pair of single-haplotype graphs were built. These graphs were used as input to `toil-vg sim`, trained as above for the different sequencer models, and configured for an average fragment length of 570 bp, a standard deviation of 165 bp, and an indel rate of 0.029%. Reads were annotated with path positions for touched reference paths. Reads touching the `hs37d5` general decoy sequence and the `NC_007605` viral decoy were excluded. Finally, we truncated to exactly 1 million pairs (2 million reads).

For the human HGSVC graph, we needed to annotate reads not only with positions along the reference paths, but also with positions along alt allele paths corresponding to large insertions, in order to assess our ability to correctly place reads that did not touch the primary reference. Since the way these paths are named by vg differs depending on the set of alt alleles in the input VCF, the single-haplotype graph approach could not be used. Instead, we simulated from the full HGSVC graph using `vg sim`, restricting the simulator to the haplotype segments representing sample NA19240 in the GBWT. The same training reads, fragment length distribution, and error parameters were used as in the 1KG case. Reads were annotated with path positions for all touched paths, including reference and alt allele paths. Because NA19240 is known to have an XX karyotype, we excluded reads annotated as touching paths in the connected component of the graph hosting the Y chromosome reference path. Reads touching paths containing “JTFH” or “KN70” in their names were also excluded, in order to remove reads simulated from decoy sequences. Finally, we truncated to exactly 1 million pairs (2 million reads).

For yeast, reads were simulated from the DBVPG6044 strain as contained in an *all-strain yeast graph*. This graph was generated using `hal2vg` v2.1 from a previously reported Cactus alignment of the five strains in the yeast graph and an additional seven strains^6^. The `vg sim` tool was used on this graph standalone, without toil-vg. The simulator was trained as described above, and used the same fragment length distribution and indel rate as for the human reads. Simulation was restricted to the paths in the graph belonging to DBVPG6044. Exactly 1 million reads (500,000 pairs) were simulated for the DBVPG6044 strain; no truncation was performed.

#### Read mapping

For our read mapping experiments on human data, we used vg release 1.27.1 to run Giraffe and VG-MAP. For mapping to yeast, we used commit 1d9385e. We tested this commit on a small number of cases with human data and saw no difference between it and the `1.27.1` version. For our simulated read mapping experiment on the 1KG graph (Figure 2 (A,D), we used a 64 haplotype sampled GBWT and for the HGSVC graph (Figure 2 (B,E), we used the full GBWT for mapping with Giraffe. We ran Bowtie2 version `2.4.1` with default settings for single-ended reads, and a maximum fragment length of 1065 for paired-end. BWA-MEM version `0.7.17-r1188` was run with default settings. GraphAligner version `1.0.11` was run with its variation graph parameter preset. HISAT2 version `2.2.1` was run with default parameters with maximum fragment length 1065 and no spliced alignment. We also ran HISAT2 with its sensitive and very sensitive settings. Minimap2 version `2.14-r883` was run with its short reads preset and no secondary alignments. A Docker container with these versions of each of the mappers is available on Docker Hub: xhchang/vg:giraffe-paper

#### Evaluating simulated read mapping

We used the path position annotations that were found during simulation to determine the correctness of read mappings. After mapping, each read was annotated again with the nearest reference position in the graph. For regions in the held out sample not included in the reference assemblies, other linear paths were also considered (such as the alt allele paths in the HGSVC graph). Any alignment that was within 100 bp of the true reference position was considered correct. For the linear mappers, we *injected* the alignments into the graph with `vg inject`, turning them into alignments to the graph, and then performed the same evaluation as for the graph mappers. Several tools allowed split read mapping and produced both primary and supplementary alignments for a single read. In these cases, we counted a read as correctly mapped if any of its alignments were correct.

#### Speed, runtime, and memory evaluation

For our speed, runtime, and memory evaluations we used an AWS EC2 i3.8xlarge node with 32 vCPUs and 244 GB of memory. We found the total runtime and memory use of each of the mappers using our 600-million-read NovSeq 6000 set. Runtime and memory were measured using `/usr/bin/time`. Each of the mappers was run using 16 threads. We were unable to map this read set with GraphAligner on the 1KG graph as we ran out of memory.

We found the speed of each mapper using our 1 million read real read sets for each sequencing technology. Each mapper except for Minimap2 was run on 16 threads; Minimap2 was run on 2 threads because it did not use all 16 threads when given. Speed was measured as reads per second per thread based on the time each tool reported as spending on mapping. We did not find the speed of GraphAligner because it logs mapping time per read, and dumping and aggregating the measurements without affecting the mapping speed would not have been feasible.

### Genotyping and evaluating short variants with Dragen

All pipelines evaluated for short variant calling performance had the same structure. First, we mapped reads to the appropriate GRCh37-based linear or graph reference using the pipeline’s mapper. Then, we genotyped the resulting alignments using Illumina’s “Dragen Bio-IT Platform” product, version 3.4, against an index generated from the `hs37d5` human genome reference. The mappers for each pipeline evaluated were Illumina’s Dragen internal aligner, HISAT2, BWA-MEM, VG-MAP, Giraffe, and Giraffe in fast mode. The Genome In a Bottle HG002 version 4.1 high-confidence VCFs were used as the truth sets for evaluating performance of variant calling^28^. The VCF was obtained from https://ftp-trace.ncbi.nlm.nih.gov/giab/ftp/release/AshkenazimTrio/HG002_NA24385_son/NISTv4.1/GRCh37/supplementaryFiles/HG002_GRCh37_CHROM1-22_v4.1_highconf.vcf.gz and the high confidence regions evaluated were based on the BED files obtained from https://ftp-trace.ncbi.nlm.nih.gov/giab/ftp/release/AshkenazimTrio/HG002_NA24385_son/NISTv4.1/GRCh37/supplementaryFiles/HG002_GRCh37_CHROM1-22_v4.1_highconf.bed. We used RTG’s `vcfeval`^29^ and Illuminas `hap.py` https://github.com/Illumina/hap.py to evaluate variant calling concordance with truth sets. All pipelines used the computational resources of the NIH HPC Biowulf cluster (http://hpc.nih.gov).

#### Graph construction

For the variant calling experiments, graph references, and their indexes for running vg mappers, were constructed using different versions of vg ranging from `1.20.0` to `1.27.0`, orchestrated by `toil-vg construct`. The graph references for the 2×250 bp read alignment experiments were constructed using vg version `v1.20.0-134-gc29c4a250`, as packaged in container quay.io/vgteam/vg:v1.20.0-134-gc29c4a250-t347-run, using the canonical `hs37d5.fa` FASTA reference file and the 1000 Genomes Project Phase 3 phased VCFs, as for the read mapping experiments, but this time selecting only variants with a minor allele frequency of greater than 1%. The graph references for the 2×150 bp read alignment experiments were constructed using vg version `v1.27.0-90-ga64b70c1f`, as packaged in container quay.io/vgteam/vg:ci-2351-a64b70c1f9345f0821e3f3a600eb8bbf4fe44bf2, using the same input files. *Primary* graphs and indexes were constructed using just the linear reference FASTA, while *snp1kg* graphs and indexes were constructed using both the FASTA and the VCFs.

Mapping with HISAT2 was done with two different reference indexes. One, obtained from https://genome-idx.s3.amazonaws.com/hisat/grch37_genome.tar.gz, was a pre-made index containing just the linear GRCh37 reference. The other, obtained from https://genome-idx.s3.amazonaws.com/hisat/grch37_snp.tar.gz, was a pre-made index containing the GRCh37 reference and the common SNVs and indels from dbSNP 144, which also includes the variants from the 1000 Genomes Project.

#### Read sets

Two different read sets were used to evaluate performance. One was the 2×150 bp paired-end FASTQ set from sample HG002 as obtained from the FDA Precision Challenge dataset. These reads are available as paired FASTQs at https://storage.googleapis.com/cmarkell-vg-wdl-dev/test_input_reads/HG002.NovaSeq.pcr-free.35x.R1.fastq.gz and https://storage.googleapis.com/cmarkell-vg-wdl-dev/test_input_reads/HG002.NovaSeq.pcr-free.35x.R2.fastq.gz. The other read set consisted of 2×250 bp paired-end FASTQ reads from sample HG002, obtained from the GIAB NovoAligned BAM at ftp://ftp-trace.ncbi.nlm.nih.gov/giab/ftp/data/AshkenazimTrio/HG002_NA24385_son/NIST_Illumina_2x250~bps/novoalign_bams/HG002.hs37d5. 2×250.bam. These reads in paired FASTQ format are available at https://storage.googleapis.com/cmarkell-vg-wdl-dev/test_input_reads/HG002_read_pair_1.fq.gz and https://storage.googleapis.com/cmarkell-vg-wdl-dev/test_input_reads/HG002_read_pair_2.fq.gz.

#### Read mapping

The HISAT2 mapping runs were done using version `2.1.0`. Only default settings, with the addition of the `--no-spliced-alignment` flag, were used during HISAT2 execution.

One pipeline used Illumina’s Dragen module for both mapping and alignment; both steps were run against the same index derived from the `hs37d5` human genome reference.

The BWA-MEM mapping runs were done using version `0.7.17-r1188` against the `hs37d5` human genome reference.

The vg alignment runs used different graph indexes depending on the mapper used (VG-MAP or Giraffe). Run details are given in Table 1. All runs used the `toil-vg map` workflow.

**Table 1.**
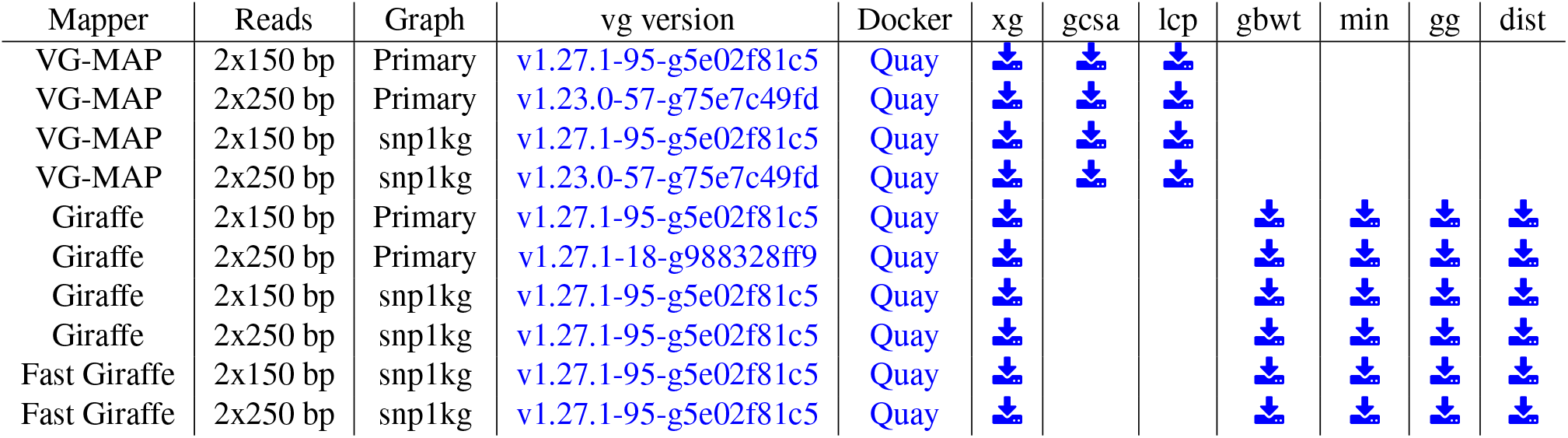
Table of parameters for genotyping experiment vg runs

| Mapper | Reads | Graph | vg version | Docker | xg | gcsa | lcp | gbwt | min | gg | dist |
| --- | --- | --- | --- | --- | --- | --- | --- | --- | --- | --- | --- |
| VG-MAP | 2x150 bp | Primary | v1.27.1-95-g5e02f81c5 | Quay |  |  |  |  |  |  |  |
| VG-MAP | 2x250 bp | Primary | v1.23.0-57-g75e7c49fd | Quay |  |  |  |  |  |  |  |
| VG-MAP | 2x150 bp | snp1kg | v1.27.1-95-g5e02f81c5 | Quay |  |  |  |  |  |  |  |
| VG-MAP | 2x250 bp | snp1kg | v1.23.0-57-g75e7c49fd | Quay |  |  |  |  |  |  |  |
| Giraffe | 2x150 bp | Primary | v1.27.1-95-g5e02f81c5 | Quay |  |  |  |  |  |  |  |
| Giraffe | 2x250 bp | Primary | v1.27.1-18-g988328ff9 | Quay |  |  |  |  |  |  |  |
| Giraffe | 2x150 bp | snp1kg | v1.27.1-95-g5e02f81c5 | Quay |  |  |  |  |  |  |  |
| Giraffe | 2x250 bp | snp1kg | v1.27.1-95-g5e02f81c5 | Quay |  |  |  |  |  |  |  |
| Fast Giraffe | 2x150 bp | snp1kg | v1.27.1-95-g5e02f81c5 | Quay |  |  |  |  |  |  |  |
| Fast Giraffe | 2x250 bp | snp1kg | v1.27.1-95-g5e02f81c5 | Quay |  |  |  |  |  |  |  |

#### DeepVariant calling

All pipelines for the DeepVariant experiment evaluation used BWA-MEM version `0.7.17-r1188` against the `hs37d5` human genome reference or VG Giraffe against the same SNP1KG graph as described in subsection Graph construction. Alignments were then deduplicated using Picard’s^52^ MarkDuplicates and indels were realigned using ABRA2^53^. The DeepVariant v1.0 code base was used for all experiments.

DeepVariant learns to call variants by training on input data from gold standard samples HG001-HG007. In all DeepVariant models, chromosome20 is fully withheld from training, allowing any sample to be evaluated in this region. Because the DeepVariant production models for Illumina are trained on samples mapped with BWA, DeepVariant likely optimizes for the mapping accuracy and quality profile of BWA. We used DeepVariant v1.0 production model, which still showed higher accuracy with Giraffe-mapped BAMs.

### Genotyping and evaluating structural variants

We replicated the evaluation performed in Hickey et al.^6^ using two SV graphs made from the HGSVC^32^ and GIAB v0.6^33^ SV catalogs, respectively. The graphs had originally been built from VCF files provided by the HGSVC and GIAB consortia. To evaluate Giraffe, we built the necessary indexes. For the minimizer index, we used the default minimizer length of 29 and window size of 11. The GBWT index was created using the greedy path-cover algorithm and *n* = 32 paths. We then mapped and genotyped the SVs in four samples: HG00514, HG00733 and NA19240 for the HGSVC graph; HG002 for the GIAB graph. The genotyped were compared to the truth-set using `sveval`^6^. We use non-repeat regions, i.e. regions of the genome that don’t overlap simple-repeats and segmental duplications, as high-confidence regions for the HGSVC graph and the combined SV graphs that are both on GRCh38. For the GIAB graph, which uses GRCh37, we use the high-confidence regions provided by the GIAB consortium^33^. The scripts to build the pangenomes, genotype the SVs and evaluate the genotyping performance are available in the repository (see Code and Data Availability).

### Building a SV pangenome combining catalogs from long-read sequencing studies

We combined variants from the three SV catalogs used previously in Hickey et al.^6^:HGSVC^32^, GIAB^33^, and SVPOP^19^. The three catalogs were first normalized and merging using bcftools version 1.9^24^.

In order to simplify the graph, we first identify near-duplicate variants that might contain error around their breakpoints. When multiple variant formed clusters of very similar near-duplicates, we use short-reads to test if one variant was clearly supported relative to the other near-duplicates. Our aim was to filter out the most obvious errors before starting to build the SV graph. We identified near-duplicates by overlapping the SVs in this combined catalog. At this stage, we defined near-duplicate SVs as deletions with a reciprocal overlap at least 80% or insertions located at less than 50 bp from each other and sizes at least 80% reciprocally similar. A network was build from this “near-duplicate” relationships to identify connected components of near-duplicate variants, which we referred to as SV clusters. For each SV cluster, we extracted the reference sequence and variants to create a local SV graph of the region. We then aligned short-reads pooled from 17 diverse individuals selected to span different populations in the 1000 Genomes Project and HGSVC. The reads aligning to the region of interest were extracted using SAMtools^24^ from the aligned reads publicly available. When reads supported one variant better than the other near-duplicates in the SV cluster, the near-duplicates were discarded.

Although many obvious duplicates had been removed, many SVs sharing sequence remained, for example if they were not covered by the short-reads used above or if they were too different to match the near-duplicate definition used above. Instead of building a pangenome naively from the variant definition, we used a hybrid approach using iterative graph augmentation. Again, the goal is to collapse as much as possible variants that share sequence content. The graph was built in chunks separated by at least 10 Kbp without any variants. For each chunk, we clustered SVs using a less stringent criterion as above: deletions with at least 30% reciprocal overlap; insertion located at less than 100bp from each other and with at least 30% reciprocally similar sizes. A graph was built for the chunk using one variant per SV cluster. One by one, the other variants were then added by aligning them to the graph and augmenting the graph. We used `vg mpmap` to align large sequences representing the SV variation flanked by 5 Kbp of sequence context. The alignment was added to the graph using `vg augment`. The chunked graphs was merged using `vg concat`.

Finally, we added unplaced and unlocated contigs from the GRCh38 reference assembly to the pangenome. We also added the `hs38d1` decoy sequences (accession GCA_000786075.2).

The scripts used to build this pangenomes are documented in the repository (see Code and Data Availability).

### Population scale analysis of SVs

#### Whole-genome sequencing datasets

The MESA whole-genome sequencing dataset was obtained from NHLBI’s TOPMed program (www.nhlbiwgs.org), freeze 5b. Paired-end 150bp reads had been sequenced on Illumina HiSeq X Ten instruments and achieved a mean depth of at least 30x. All sequencing had used PCR-free library preparation kits.

For the 1000 Genomes Project samples, we used public whole-genome sequencing data generated at the New York Genome Center (see Code and Data Availability). Samples had been sequenced to a minimum of 30x mean genome coverage using PCR-free sequencing libraries on the Illumina NovaSeq 6000 sequencing instrument, with 2×150bp reads. All cell lines had been obtained from the Coriell Institute for Medical Research and had been consented for full public release of genomic data. Of note, the reads were downsampled to 20x in both datasets to reduce the computing cost.

#### Pangenome genotyping workflow

A genotyping pipeline was implemented in WDL and ran using the BioData Catalyst ecosystem^34^ providing access to the TOPMed and 1000 Genomes Project datasets. The WDL pipeline was deposited on Dockstore^35^ and ran on Terra. The estimated cost per sample was estimated from Google Billing Reports. Resources used for each sample or step in the workflow were extracted using `terra-utils` notebooks from the BioData Catalyst Collection on Terra.

#### Selection of diverse individuals in the MESA cohort

The MESA cohort is a medical research study involving white, African American, Hispanic/Latino, and Chinese adults from 6 U.S. communities. We selected a subset of 2 thousand individuals in the MESA cohort from the TOPMed freeze 5b. The individuals were selected to cover as uniformly as possible the space defined by the top 11 principal components provided by the TOPMed consortium. These principal components (PCs) were derived from SNVs genotypes across the full TOPMed dataset. Briefly, the selection process started with the two individuals that located the furthest from each other in the PC space. New individuals were then selected, one by one, to maximize their distance to already selected individuals.

#### Combining SV genotypes across samples

SVs were merged using bcftools version 1.9^24^. For each sample, we split multi-allelic variants, normalize them and sort the output VCF. The output of `vg call` is derived from paths in the graph (like haplotypes) that traverse the SV site. For large SV sites, it means that one called variant may contain multiple smaller canonical SVs. In addition, this path-based reporting of the variants can sometimes separate homozygous canonical variants into two heterozygous variants embedded within larger paths. Hence, during normalization, we aligned the reference and alternate alleles in the VCF file to break down each variant into canonical SVs. We also combined heterozygous variants that matched exactly into homozygous calls. These SVs were then left-aligned and sorted using bcftools. Finally, the VCF files were combined using `bcftools merge` using the `-0 -m none` parameters to avoid multi-allelic records. The corresponding WDL workflow has been deposited on Dockstore^54^.

#### Clustering of SV alleles into SV sites

Deletions were matched if their reciprocal overlap was at least 80%. Insertions were matched if located at less than 20 and their sizes were reciprocally similar at least 80%. Annotated simple repeats were also used to extend the range used to match SVs because similar variants might be placed in different position of an annotated VNTR. We built a network using these matches to identify connected component representing SV sites where similar alleles cluster. If the component formed a clique, i.e. all SVs matched all other SVs, the SV site contained alleles that differed by small variants (relative to the SV size). If not, the SV site contained alleles with significant differences, usually due to important repeat expansion or contraction at a VNTR site. For most analysis at the SV site level, we used the most frequent allele’s information, such as boundaries or size. The corresponding scripts are documented in the repository (see Code and Data Availability).

#### Allele frequency patterns across populations

We performed a principal component analysis of the individuals using the allele counts across all SV sites. The allele counts represent the number of non-reference alleles called in each individual at each SV locus.

For the MESA cohort, we compared the principal components derived from the SV allele counts to the principal components produced by the TOPMed consortium using short variants. Population information was not available for these individuals so we used the top three principal components from our SV analysis to cluster them and explore population patterns further. We used hierarchical clustering (`ward.D` method in R’s `hclust` function) and cut the tree to obtain 6 clusters.

For the 1000 Genomes Project dataset, we used public information about the samples’ super populations to color the PCA graphs and explore population patterns.

For each SV site, we compared the allele frequencies between the clusters/super populations. To illustrate global population differences, we computed the range of allele frequencies across clusters/super populations. This distribution was compared to an empirical null distribution produced by shuffling the population assignments. As population-centric metric, we contrasted the allele frequency at each SV site in a super population versus the median allele frequency across all super populations. Again, we used the null dataset to illustrate the enrichment in SVs whose frequency in the super population of interest is different from the global allele frequency. We highlight SVs with a frequency differing by more than 10% from the median frequency, a threshold at which only a few SVs are expected to show population patterns based on the permuted dataset. The corresponding scripts are documented in the repository (see Code and Data Availability).

#### Comparison with existing large-scale SV catalogs

We compared the variants genotyped in this study with three SV catalogs that screened large populations:

1. the 1000 Genomes Project phase 3^36^ (2,504 individuals) downloaded from http://ftp.1000genomes.ebi.ac.uk/vol1/ftp/phase3/integrated_sv_map/supporting/GRCh38_positions/ALL.wgs.mergedSV.v8.20130502.svs.genotypes.GRCh38.vcf.gz
2. gnomAD-SV^37^ v2 (more than 14 thousand individuals), lifted over to GRCh38
3. SVPOP^19^ (440 individuals) downloaded from http://ftp.1000genomes.ebi.ac.uk/vol1/ftp/data_collections/hgsv_sv_discovery/working/20181025_EEE_SV-Pop_1/VariantCalls_EEE_SV-Pop_1/EEE_SV-Pop_1.ALL.genotypes.20181204.vcf.gz.

These three public catalogs provide allele frequency estimate that we compared with the frequency of matched variants in our datasets. Variants were matched with weaker matching criteria than before to avoid over-estimating the amount of novel variants. We matched deletions with at least 30% reciprocal overlap and insertions located at less than 200 bp from each other and at least 30% reciprocally similar sizes. Annotated simple repeats were also used to extend the range used to match SVs. The corresponding scripts are documented in the repository (see Code and Data Availability).

#### Annotation of the SVs

The RepeatMasker and SimpleRepeat annotation downloaded from the UCSC Genome Browser FTP server. We considered a SV to overlap simple repeats if at least 50% of its region overlapped with the SimpleRepeat track or repeats from the *Simple_repeat* class in the RepeatMasker annotation. Low-complexity regions were extracted from the RepeatMasker annotation. The SVs were overlapped with gene annotation from Genecode v35. Promoter regions were defined from 2 Kbp up-stream to 200 bp downstream of the transcription start site. We annotated SVs that overlapped coding regions, UTRs, promoters or introns of protein-coding genes, in this order. This means that a SVs spanning a full gene is annotated as *coding*, or, for example, that a *promoter* SV overlapped neither coding nor untranslated regions. Because the gene expression in the GEUVADIS is from lymphoblastoid cell lines, we used candidate regulatory regions for GM12878 from the ENCODE project^55^,56 (ENCFF590IMH) when annotating SV-eQTLs. As before, a SV was preferentially annotated using the following order: coding, UTR, promoter, intron, regulatory region, intergenic region.

#### Expression Quantitative Trait Locus discovery

We used gene expression provided by the GEUVADIS consortium^39^. We tested for cis-eQTLs using a 1 Mbp range around the gene’s transcription start site using Matrix eQTL^57^. First, all 445 samples with gene expression information were analyzed jointly. Additionally, we also analyzed the CEU, FIN, GBR, and TSI samples together, and the YRI population separately. In each analysis, we only tested SVs with a minor allele frequencies of at least 5%. The gene expression had been corrected for batch effects by the GEUVADIS consortium using PEER^39^. Covariates of the model included gender and the top four principal components derived from the SV genotypes. The p-values were corrected for multiple-testing using Benjamini-Hochberg correction. We report eQTLs with an adjusted p-value (false discovery rate) of 1%.

We annotated each SV-eQTL using the gene annotation (see above). SVs were split between Svs associated with increased or decreased gene expression. As baseline for the functional overlaps, we selected genes with similar sizes as the genes involved in eQTLs. We then selected all common SVs (allele frequency ≥ 5%) and located at less than 1 Mbp of these genes.

## Code and Data Availability

An overview of the data generated for this paper is available at https://cglgenomics.ucsc.edu/giraffe-data/. The dataset has IPFS hash QmNNjnJXARUeaaLhjQzQbX5XJRuKEjmUcF5yThyuo3zc7D.

The latest version of vg, including the Giraffe implementation, is available at https://github.com/vgteam/vg. The commands and scripts used for the analysis presented in this study have been deposited at https://github.com/vgteam/giraffe-sv-paper. This repository also contains paths to download the public datasets and annotation used in this study.

Data used in the Giraffe read mapping experiments, including the 1KG, HGSVC, and yeast target graphs, the linear control graphs, the graphs used to simulate reads, and the simulated reads themselves, can be found at https://cgl.gi.ucsc.edu/data/giraffe/mapping/.

The SV pangenomes and SV catalogs annotated with allele frequencies are hosted at https://cgl.gi.ucsc.edu/data/giraffe/calling/. This includes SVs with strong inter-super-population frequency patterns (Supplementary Table S15), SV-eQTLs (Supplementary Table S16), and SVs overlapping protein-coding genes (Supplementary Table S17).

The public high-coverage sequencing dataset from the 1000 Genomes Project is available at https://www.internationalgenomeorg/data-portal/data-collection/30x-grch38, including ENA projects PRJEB31736 and PRJEB36890. The gene expression data was download from ArrayExpress E-GEUV-1 (GD462.GeneQuantRPKM.50FN.samplename.resk10.txt.gz We downloaded the call sets from the ENCODE portal (Sloan et al. 2016) (https://www.encodeproject.org/) with the following identifiers: ENCFF590IMH.

Individual whole-genome sequence data for TOPMed whole genomes are available through dbGaP. The dbGaP accession numbers for the Multi-Ethnic Study of Atherosclerosis (MESA) is phs001416. Data in dbGaP can be downloaded by controlled access with an approved application submitted through their website: https://www.ncbi.nlm.nih.gov/gap.

## Acknowledgments

The high coverage sequencing data for the 1000 Genomes Project were generated at the New York Genome Center with funds provided by NHGRI Grant 3UM1HG008901-03S1.

MESA and the MESA SHARe projects are conducted and supported by the National Heart, Lung, and Blood Institute (NHLBI) in collaboration with MESA investigators. Support for MESA is provided by contracts 75N92020D00001, HHSN268201500003I, N01-HC-95159, 75N92020D00005, N01-HC-95160, 75N92020D00002, N01-HC-95161, 75N92020D00003, N01-HC-95162, 75N92020D00006, N01-HC-95163, 75N92020D00004, N01-HC-95164, 75N92020D00007, N01-HC-95165, N01-HC-95166, N01-HC-95167, N01-HC-95168, N01-HC-95169, UL1-TR-000040, UL1-TR-001079, UL1-TR-001420. Funding for SHARe genotyping was provided by NHLBI Contract N02-HL-64278. Genotyping was performed at Affymetrix (Santa Clara, California, USA) and the Broad Institute of Harvard and MIT (Boston, Massachusetts, USA) using the Affymetrix Genome-Wide Human SNP Array 6.0. Also supported in part by the National Center for Advancing Translational Sciences, CTSI grant UL1TR001881, and the National Institute of Diabetes and Digestive and Kidney Disease Diabetes Research Center (DRC) grant DK063491 to the Southern California Diabetes Endocrinology Research Center. Whole genome sequencing (WGS) for the Trans-Omics in Precision Medicine (TOPMed) program was supported by the National Heart, Lung and Blood Institute (NHLBI). WGS for “NHLBI TOPMed: Multi-Ethnic Study of Atherosclerosis (MESA)” (phs001416) was performed at the Broad Institute of MIT and Harvard (3U54HG003067-13S1 and HHSN268201500014C). Core support including centralized genomic read mapping and genotype calling, along with variant quality metrics and filtering were provided by the TOPMed Informatics Research Center (3R01HL-117626-02S1; contract HHSN268201800002I). Core support including phenotype harmonization, data management, sample-identity QC, and general program coordination were provided by the TOPMed Data Coordinating Center (R01HL-120393; U01HL-120393; contract HHSN268201800001I). We gratefully acknowledge the studies and participants who provided biological samples and data for TOPMed. The views expressed in this manuscript are those of the authors and do not necessarily represent the views of the National Heart, Lung, and Blood Institute, the National Institutes of Health or the U.S. Department of Health and Human Services.

Research reported in this publication was supported by the National Human Genome Research Institute of the National Institutes of Health under Award Numbers U41HG010972, 1R01HG008742, U01HG010961, R01HG009737 and 2U41HG007234. Research reported in this publication was supported by the National Heart, Lung, And Blood Institute of the National Institutes of Health under Award Number U01HL137183. Research reported in this publication was supported by the BioData Catalyst Fellows Program of the National Institutes of Health through the University of North Carolina at Chapel Hill, under Award Number 1 OT3 HL147154. The content is solely the responsibility of the authors and does not necessarily represent the official views of the National Institutes of Health. JAS was supported by the Carlsberg Foundation.

