## Supplementary Material for "Genotyping common, large structural variations in 5,202 genomes using pangenomes, the Giraffe mapper, and the vg toolkit"

### Supplementary Tables

| Pairing | Mapper | % Correct | % MAPQ 60 | % Incorrect and MAPQ 60 |
|---|---|---|---|---|
| single | Giraffe primary | 98.16 | 93.52 | 0.00255 |
| | VG-MAP primary | 98.24 | 94.04 | 0.00050 |
| | Bowtie2 | 98.04 | 84.90 | 0.00005 |
| | BWA-MEM | 98.24 | 93.01 | 0.00015 |
| | Minimap2 | 98.03 | 92.03 | 0.00305 |
| | Giraffe sampled | 98.24 | 94.05 | 0.00020 |
| | fast Giraffe sampled | 98.10 | 91.58 | 0.00015 |
| | VG-MAP | 98.19 | 93.71 | 0.00010 |
| | GraphAligner | 89.85 | - | - |
| | Hisat2 default | 98.16 | 97.50 | 0.12345 |
| | Hisat2 sensitive | 98.24 | 97.60 | 0.14365 |
| | Hisat2 very sensitive | 98.31 | 97.72 | 0.20930 |
| paired | Giraffe primary | 99.01 | 95.79 | 0.00070 |
| | VG-MAP primary | 99.05 | 96.19 | 0.00060 |
| | Bowtie2 | 99.00 | 90.87 | 0.00045 |
| | BWA-MEM | 99.05 | 95.68 | 0.00045 |
| | Minimap2 | 98.84 | 94.93 | 0.01075 |
| | Giraffe sampled | 99.03 | 95.95 | 0.00010 |
| | fast Giraffe sampled | 99.01 | 94.53 | 0.00000 |
| | VG-MAP | 99.05 | 96.07 | 0.00005 |
| | Hisat2 default | 98.93 | 98.46 | 0.08120 |
| | Hisat2 sensitive | 99.10 | 98.63 | 0.07915 |
| | Hisat2 very sensitive | 99.14 | 98.64 | 0.07615 |

**Table S1. Mapping accuracy on NovaSeq 6000 reads mapped to the 1KG graph/GRCh37 reference** Each mapper was run on 2 million simulated reads and assessed for the percent of reads that were mapped correctly, the percent of reads that were assigned mapping quality 60, and the percent of reads that were incorrect and assigned mapping quality 60. *Bowtie2 had a maximum mapping quality of 42. GraphAligner did not assign mapping quality

| Pairing | Mapper | % Correct | % MAPQ 60 | % Incorrect and MAPQ 60 |
|---|---|---|---|---|
| single | Giraffe primary | 98.13 | 93.31 | 0.00230 |
| | VG-MAP primary | 98.21 | 93.97 | 0.00075 |
| | Bowtie2 | 97.92 | 85.62 | 0.00010 |
| | BWA-MEM | 98.21 | 92.92 | 0.00040 |
| | Minimap2 | 97.98 | 91.91 | 0.00455 |
| | Giraffe sampled | 98.21 | 93.86 | 0.00030 |
| | fast Giraffe sampled | 98.04 | 91.68 | 0.00060 |
| | VG-MAP | 98.16 | 93.64 | 0.00025 |
| | GraphAligner | 89.60 | - | - |
| | Hisat2 default | 97.94 | 97.36 | 0.18050 |
| | Hisat2 sensitive | 98.07 | 97.52 | 0.21685 |
| | Hisat2 very sensitive | 98.15 | 97.60 | 0.24970 |
| paired | Giraffe primary | 98.99 | 95.83 | 0.00065 |
| | VG-MAP primary | 99.03 | 96.16 | 0.00075 |
| | Bowtie2 | 98.96 | 91.05 | 0.00035 |
| | BWA-MEM | 99.02 | 95.64 | 0.00065 |
| | Minimap2 | 98.82 | 94.84 | 0.01195 |
| | Giraffe sampled | 99.02 | 95.98 | 0.00005 |
| | fast Giraffe sampled | 99.00 | 94.72 | 0.00005 |
| | VG-MAP | 99.04 | 96.03 | 0.00015 |
| | Hisat2 default | 98.72 | 98.29 | 0.12050 |
| | Hisat2 sensitive | 98.97 | 98.55 | 0.12315 |
| | Hisat2 very sensitive | 99.11 | 98.62 | 0.09495 |

**Table S2. Mapping accuracy on HiSeq X Ten reads mapped to the 1KG graph/GRCh37 reference** Each mapper was run on 2 million simulated reads and assessed for the percent of reads that were mapped correctly, the percent of reads that were assigned mapping quality 60, and the percent of reads that were incorrect and assigned mapping quality 60. *Bowtie2 had a maximum mapping quality of 42. GraphAligner did not assign mapping quality

| Pairing | Mapper | % Correct | % MAPQ 60 | % Incorrect and MAPQ 60 |
|---|---|---|---|---|
| single | Giraffe primary | 98.74 | 96.46 | 0.00505 |
| | VG-MAP primary | 98.82 | 95.71 | 0.00180 |
| | Bowtie2 | 97.44 | 86.32 | 0.00005 |
| | BWA-MEM | 98.76 | 94.44 | 0.05190 |
| | Minimap2 | 98.62 | 94.81 | 0.05270 |
| | Giraffe sampled | 98.80 | 96.58 | 0.00250 |
| | fast Giraffe sampled | 98.69 | 95.66 | 0.00225 |
| | VG-MAP | 98.82 | 95.57 | 0.00130 |
| | GraphAligner | 93.70 | - | - |
| | Hisat2 default | 96.20 | 96.09 | 0.33515 |
| | Hisat2 sensitive | 97.25 | 97.08 | 0.31725 |
| | Hisat2 very sensitive | 97.56 | 97.37 | 0.36845 |
| paired | Giraffe primary | 99.21 | 97.14 | 0.00275 |
| | VG-MAP primary | 99.25 | 97.11 | 0.00220 |
| | Bowtie2 | 98.00 | 89.77 | 0.00005 |
| | BWA-MEM | 99.20 | 96.29 | 0.05535 |
| | Minimap2 | 99.02 | 94.21 | 0.06435 |
| | Giraffe sampled | 99.23 | 97.18 | 0.00210 |
| | fast Giraffe sampled | 99.21 | 96.64 | 0.00185 |
| | VG-MAP | 99.26 | 97.05 | 0.00155 |
| | Hisat2 default | 96.57 | 96.54 | 0.38540 |
| | Hisat2 sensitive | 97.77 | 97.72 | 0.34875 |
| | Hisat2 very sensitive | 98.16 | 97.93 | 0.23270 |

**Table S3. Mapping accuracy on HiSeq 2500 mapped to the 1KG graph/GRCh37 reference** Each mapper was run on 2 million simulated reads and assessed for the percent of reads that were mapped correctly, the percent of reads that were assigned mapping quality 60, and the percent of reads that were incorrect and assigned mapping quality 60. *Bowtie2 had a maximum mapping quality of 42. GraphAligner did not assign mapping quality

| Pairing | Mapper | % Correct | % MAPQ 60 | % Incorrect and MAPQ 60 |
|---|---|---|---|---|
| single | Giraffe primary | 97.09 | 92.68 | 0.04850 |
| | VG-MAP primary | 97.17 | 92.62 | 0.04685 |
| | Bowtie2 | 97.02 | 84.20 | 0.02645 |
| | BWA-MEM | 97.16 | 91.61 | 0.03925 |
| | Minimap2 | 96.96 | 90.93 | 0.04220 |
| | Giraffe full | 97.17 | 92.64 | 0.00045 |
| | fast Giraffe full | 97.02 | 90.26 | 0.00070 |
| | VG-MAP | 97.24 | 92.59 | 0.00035 |
| | GraphAligner | 91.95 | - | - |
| | Hisat2 default | 96.92 | 96.17 | 0.20935 |
| | Hisat2 sensitive | 96.98 | 96.27 | 0.24655 |
| | Hisat2 very sensitive | 97.05 | 96.40 | 0.30310 |
| paired | Giraffe primary | 98.27 | 94.61 | 0.05465 |
| | VG-MAP primary | 98.32 | 94.79 | 0.04850 |
| | Bowtie2 | 98.28 | 89.54 | 0.03220 |
| | BWA-MEM | 98.38 | 94.34 | 0.04275 |
| | Minimap2 | 98.09 | 93.72 | 0.06800 |
| | Giraffe full | 98.38 | 94.54 | 0.00035 |
| | fast Giraffe full | 98.34 | 93.10 | 0.00035 |
| | VG-MAP | 98.42 | 94.74 | 0.00385 |
| | Hisat2 default | 98.05 | 97.76 | 0.35310 |
| | Hisat2 sensitive | 98.18 | 97.98 | 0.44865 |
| | Hisat2 very sensitive | 98.23 | 97.97 | 0.38985 |

**Table S4. Mapping accuracy on NovaSeq 6000 reads mapped to the HGSVC graph/GRCh38 reference** Eaach mapper was run on 2 million simulated reads and assessed for the percent of reads that were mapped correctly, the percent of reads that were assigned mapping quality 60, and the percent of reads that were incorrect and assigned mapping quality 60. *Bowtie2 had a maximum mapping quality of 42. GraphAligner did not assign mapping quality

| Pairing | Mapper | % Correct | % MAPQ 60 | % Incorrect and MAPQ 60 |
|---|---|---|---|---|
| single | Giraffe primary | 97.10 | 92.57 | 0.0487 |
| | VG-MAP primary | 97.20 | 92.63 | 0.04585 |
| | Bowtie2 | 96.94 | 85.15 | 0.02865 |
| | BWA-MEM | 97.19 | 91.62 | 0.03885 |
| | Minimap2 | 96.96 | 90.86 | 0.04140 |
| | Giraffe full | 97.12 | 92.53 | 0.00090 |
| | fast Giraffe full | 97.01 | 90.44 | 0.00125 |
| | VG-MAP | 97.27 | 92.58 | 0.00035 |
| | GraphAligner | 91.89 | - | - |
| | Hisat2 default | 96.74 | 96.14 | 0.30020 |
| | Hisat2 sensitive | 96.84 | 96.27 | 0.34115 |
| | Hisat2 very sensitive | 96.92 | 96.34 | 0.34120 |
| paired | Giraffe primary | 98.27 | 94.72 | 0.05405 |
| | VG-MAP primary | 98.35 | 94.84 | 0.04750 |
| | Bowtie2 | 98.25 | 89.98 | 0.03325 |
| | BWA-MEM | 98.40 | 94.36 | 0.04280 |
| | Minimap2 | 98.07 | 93.69 | 0.06815 |
| | Giraffe full | 98.37 | 94.65 | 0.00060 |
| | fast Giraffe full | 98.34 | 93.39 | 0.00060 |
| | VG-MAP | 98.43 | 94.79 | 0.00420 |
| | Hisat2 default | 97.85 | 97.64 | 0.40260 |
| | Hisat2 sensitive | 98.04 | 97.94 | 0.51215 |
| | Hisat2 very sensitive | 98.18 | 97.98 | 0.42855 |

**Table S5. Mapping accuracy on HiSeq X Ten reads mapped to the HGSVC graph/GRCh38 reference** Each mapper was run on 2 million simulated reads and assessed for the percent of reads that were mapped correctly, the percent of reads that were assigned mapping quality 60, and the percent of reads that were incorrect and assigned mapping quality 60. *Bowtie2 had a maximum mapping quality of 42. GraphAligner did not assign mapping quality

| Pairing | Mapper | % Correct | % MAPQ 60 | % Incorrect and MAPQ 60 |
|---|---|---|---|---|
| single | Giraffe primary | 97.89 | 95.34 | 0.0698 |
| | VG-MAP primary | 97.99 | 94.38 | 0.05715 |
| | Bowtie2 | 96.56 | 85.74 | 0.03405 |
| | BWA-MEM | 97.93 | 93.08 | 0.09025 |
| | Minimap2 | 97.78 | 93.59 | 0.09525 |
| | Giraffe full | 97.98 | 95.30 | 0.00330 |
| | fast Giraffe full | 97.88 | 94.35 | 0.00260 |
| | VG-MAP | 98.0 | 94.35 | 0.00170 |
| | GraphAligner | 94.87 | - | - |
| | Hisat2 default | 95.14 | 95.29 | 0.69075 |
| | Hisat2 sensitive | 96.17 | 96.23 | 0.66070 |
| | Hisat2 very sensitive | 96.54 | 96.40 | 0.54060 |
| paired | Giraffe primary | 98.68 | 96.09 | 0.0636 |
| | VG-MAP primary | 98.79 | 95.93 | 0.05845 |
| | Bowtie2 | 97.33 | 88.71 | 0.03860 |
| | BWA-MEM | 98.75 | 94.97 | 0.09730 |
| | Minimap2 | 98.44 | 93.02 | 0.12805 |
| | Giraffe full | 98.79 | 96.04 | 0.00235 |
| | fast Giraffe full | 98.76 | 95.43 | 0.00260 |
| | VG-MAP | 98.89 | 95.90 | 0.00280 |
| | Hisat2 default | 95.68 | 96.00 | 0.75595 |
| | Hisat2 sensitive | 96.90 | 97.32 | 0.82530 |
| | Hisat2 very sensitive | 97.37 | 97.53 | 0.63680 |

**Table S6. Mapping accuracy on HiSeq 2500 reads mapped to the HGSVC graph/GRCh38 reference** Each mapper was run on 2 million simulated reads and assessed for the percent of reads that were mapped correctly, the percent of reads that were assigned mapping quality 60, and the percent of reads that were incorrect and assigned mapping quality 60. *Bowtie2 had a maximum mapping quality of 42. GraphAligner did not assign mapping quality

| Pipeline | TP | FP | FN | Precision | Sensitivity | F-measure |
|---|---|---|---|---|---|---|
| BWA-MEM | 3,846,860 | 14,769 | 22,934 | 0.9962 | 0.9941 | 0.9951 |
| HISAT2 grch37_snp index | 3,825,654 | 42,120 | 44,218 | 0.9891 | 0.9886 | 0.9888 |
| HISAT2 grch37_genome index | 3,813,621 | 47,070 | 56,190 | 0.9878 | 0.9855 | 0.9866 |
| DRAGEN | 3,847,559 | 11,992 | 22,233 | 0.9969 | 0.9943 | 0.9956 |
| VG-MAP PRIMARY | 3,846,702 | 12,337 | 23,091 | 0.9968 | 0.9940 | 0.9954 |
| VG-MAP SNP1KG | 3,849,411 | 12,873 | 20,383 | 0.9967 | 0.9947 | 0.9957 |
| VG GIRAFFE PRIMARY | 3,850,325 | 13,094 | 19,474 | 0.9966 | 0.9950 | 0.9958 |
| VG GIRAFFE SNP1KG | **3,853,165** | **11,786** | **16,635** | **0.9970** | **0.9957** | **0.9963** |
| VG GIRAFFE FAST SNP1KG | 3,852,436 | 12,775 | 17,365 | 0.9967 | 0.9955 | 0.9961 |

**Table S7.** VCFeval performance of linear and graph-based pipelines using 2x150bp reads with respect to HG002 GIAB v4.1 truth variant call sets in high confident regions. Best values in each column are highlighted in bold text.

| Pipeline | TP | FP | FN | Precision | Sensitivity | F-measure |
|---|---|---|---|---|---|---|
| BWA-MEM | 3,850,507 | 10,305 | 19,290 | 0.9973 | 0.9950 | 0.9962 |
| DRAGEN | 3,851,275 | 8,419 | 18,522 | **0.9978** | 0.9952 | 0.9965 |
| VG-MAP PRIMARY | 3,850,358 | **8,404** | 19,439 | **0.9978** | 0.9950 | 0.9964 |
| VG-MAP SNP1KG | 3,852,917 | 8,917 | 16,891 | 0.9977 | 0.9956 | 0.9967 |
| VG GIRAFFE PRIMARY | 3,854,046 | 9,281 | 15,765 | 0.9976 | 0.9959 | 0.9968 |
| VG GIRAFFE SNP1KG | **3,855,804** | 8,748 | **14,007** | 0.9977 | **0.9964** | **0.9971** |
| VG GIRAFFE FAST SNP1KG | 3,855,317 | 9,128 | 14,493 | 0.9976 | 0.9963 | 0.9969 |

**Table S8.** VCFeval performance of linear and graph-based pipelines using 2x250bp reads with respect to HG002 GIAB v4.1 truth variant call sets in high confident regions. Best values in each column are highlighted in bold text.

| Pipeline | Var Type | TP | FN | FP | Recall | Precision | F1 |
|---|---|---|---|---|---|---|---|
| BWA-MEM | INDELS | 519,816 | 2,927 | 2,264 | 0.994401 | 0.995842 | 0.995121 |
| | SNPS | 3,327,183 | 19,977 | 12,542 | 0.994032 | 0.996246 | 0.995138 |
| HISAT2 grch37_snp index | INDELS | 505,484 | 17,259 | 6,534 | 0.966984 | 0.987733 | 0.977248 |
| | SNPS | 3,320,214 | 26,946 | 35,664 | 0.991950 | 0.989376 | 0.990661 |
| HISAT2 grch37_genome index | INDELS | 505,740 | 17,003 | 6,372 | 0.967474 | 0.988054 | 0.977655 |
| | SNPS | 3,307,986 | 39,174 | 40,766 | 0.988296 | 0.987830 | 0.988063 |
| DRAGEN | INDELS | 519,843 | 2,900 | 2,310 | 0.994452 | 0.995758 | 0.995105 |
| | SNPS | 3,327,856 | 19,304 | 9,732 | 0.994233 | 0.997085 | 0.995657 |
| VG-MAP PRIMARY | INDELS | 519,261 | 3,482 | **2,150** | 0.993339 | **0.996046** | 0.994691 |
| | SNPS | 3,327,576 | 19,584 | 10,230 | 0.994149 | 0.996936 | 0.995541 |
| VG-MAP SNP1KG | INDELS | 519,281 | 3,462 | 2,245 | 0.993377 | 0.995872 | 0.994623 |
| | SNPS | 3,330,264 | 16,896 | 10,673 | 0.994952 | 0.996806 | 0.995878 |
| VG GIRAFFE PRIMARY | INDELS | 519,436 | 3,307 | 2,371 | 0.993674 | 0.995643 | 0.994657 |
| | SNPS | 3,331,023 | 16,137 | 10,763 | 0.995179 | 0.996780 | 0.995979 |
| VG GIRAFFE SNP1KG | INDELS | **519,893** | **2,850** | 2,166 | **0.994548** | 0.996022 | **0.995284** |
| | SNPS | **3,333,400** | **13,760** | **9,672** | **0.995889** | **0.997108** | **0.996498** |
| VG GIRAFFE FAST SNP1KG | INDELS | 519,877 | 2,866 | 2,206 | 0.994517 | 0.995948 | 0.995232 |
| | SNPS | 3,332,687 | 14,473 | 10,619 | 0.995676 | 0.996825 | 0.996250 |

**Table S9.** Hap.py performance of linear and graph-based pipelines using 2x150bp reads with respect to HG002 GIAB v4.1 truth variant call sets in high confident regions. Best values in each column are highlighted in bold text.

| Pipeline | Var Type | TP | FN | FP | Recall | Precision | F1 |
|---|---|---|---|---|---|---|---|
| BWA-MEM | INDELS | **519,995** | **2,748** | 1,748 | **0.994743** | 0.996787 | **0.995764** |
| | SNPS | 3,330,639 | 16,521 | 8,606 | 0.995064 | 0.997424 | 0.996243 |
| DRAGEN | INDELS | 519,961 | 2,782 | 1,783 | 0.994678 | 0.996722 | 0.995699 |
| | SNPS | 3,331,439 | 15,721 | **6,686** | 0.995303 | **0.997998** | 0.996649 |
| VG-MAP PRIMARY | INDELS | 519,398 | 3,345 | **1,673** | 0.993601 | **0.996920** | 0.995258 |
| | SNPS | 3,331,084 | 16,076 | 6,780 | 0.995197 | 0.997969 | 0.996581 |
| VG-MAP SNP1KG | INDELS | 519,328 | 3,415 | 1,864 | 0.993467 | 0.996569 | 0.995016 |
| | SNPS | 3,333,698 | 13,462 | 7,105 | 0.995978 | 0.997874 | 0.996925 |
| VG GIRAFFE PRIMARY | INDELS | 519,365 | 3,378 | 1,872 | 0.993538 | 0.996555 | 0.995044 |
| | SNPS | 3,334,790 | 12,370 | 7,452 | 0.996304 | 0.997771 | 0.997037 |
| VG GIRAFFE SNP1KG | INDELS | 519,763 | 2,980 | 1,814 | 0.994299 | 0.996664 | 0.995480 |
| | SNPS | **3,336,153** | **11,007** | 6,984 | **0.996712** | 0.997912 | **0.997311** |
| VG GIRAFFE FAST SNP1KG | INDELS | 519,736 | 3,007 | 1,827 | 0.994248 | 0.996640 | 0.995442 |
| | SNPS | 3,335,695 | 11,465 | 7,352 | 0.996575 | 0.997802 | 0.997188 |

**Table S10.** Hap.py performance of linear and graph-based pipelines using 2x250bp reads with respect to HG002 GIAB v4.1 truth variant call sets in high confident regions. Best values in each column are highlighted in bold text.

| Pipeline | Var Type | TP | FN | FP | Recall | Precision | F1 |
|---|---|---|---|---|---|---|---|
| BWA-MEM + Dragen | INDELS | 11,222 | 50 | 29 | 0.995564 | 0.997519 | 0.996541 |
| | SNPS | 70,536 | 329 | 121 | 0.995357 | 0.998288 | 0.996821 |
| VG GIRAFFE + Dragen | INDELS | 11,231 | 41 | 24 | 0.996363 | 0.997948 | 0.997155 |
| | SNPS | **70,604** | **261** | 92 | **0.996317** | 0.998699 | 0.997507 |
| Dragen MAP + Dragen | INDELS | 11,230 | 42 | 21 | 0.996274 | 0.998204 | 0.997238 |
| | SNPS | 70,535 | 330 | 82 | 0.995343 | 0.998839 | 0.997088 |
| BWA-MEM + DeepVariant | INDELS | 11,229 | 43 | **19** | 0.996185 | 0.998374 | 0.997278 |
| | SNPS | 70,509 | 356 | **18** | 0.994976 | **0.999745** | 0.997355 |
| VG GIRAFFE + DeepVariant | INDELS | **11,239** | **33** | **19** | **0.997072** | **0.998375** | **0.997723** |
| | SNPS | 70,547 | 318 | 32 | 0.995513 | 0.999547 | **0.997526** |

**Table S11.** Hap.py performance between DeepVariant and Dragen variant callers on linear and giraffe snp1kg mappers using 2x150bp reads with respect to HG002 GIAB v4.1 truth variant call sets in high confident regions on chromosome 20. Best values in each column are highlighted in bold text.

| Pre-empted jobs | Dataset | Workflows | Core.hour |
|---|---|---|---|
| 0 | all | 2910 | 194.388 |
| 0 | MESA | 1221 | 197.981 |
| 0 | 1000GP | 1689 | 191.791 |
| 1 | all | 1358 | 220.046 |
| 1 | MESA | 517 | 222.205 |
| 1 | 1000GP | 841 | 218.718 |
| 2 | all | 439 | 240.992 |
| 2 | MESA | 172 | 248.246 |
| 2 | 1000GP | 267 | 236.318 |
| 3+ | all | 209 | 270.351 |
| 3+ | MESA | 88 | 282.007 |
| 3+ | 1000GP | 121 | 261.874 |

**Table S12. Computing resources used when genotyping 2,000 samples from the MESA cohort and 3,202 samples from the 1000 Genomes Project dataset.** Pre-emptible instances on Google Cloud were used for all jobs. As expected, the average resource required was higher for workflows where some jobs were pre-empted. Note: The sequencing reads were down-sampled to ∼20x coverage.

| Task | CPU | Memory (Gb) | Dataset | Core.hour |
|---|---|---|---|---|
| CRAM conversion | 8 | 50 | all | 12.867 (7.6%) |
| CRAM conversion | 8 | 50 | MESA | 13.22 (7.6%) |
| CRAM conversion | 8 | 50 | 1000GP | 12.636 (7.6%) |
| mapping | 32 | 100 | all | 146.261 (75.2%) |
| mapping | 32 | 100 | MESA | 157.66 (77.1%) |
| mapping | 32 | 100 | 1000GP | 138.782 (73.9%) |
| genotyping | 16 | 100 | all | 24.891 (17.2%) |
| genotyping | 16 | 100 | MESA | 27.112 (15.4%) |
| genotyping | 16 | 100 | 1000GP | 23.433 (18.5%) |

**Table S13. Computing resources for each task in the WDL genotyping workflow.** The table shows numbers for the workflows where no jobs were pre-empted.

| Public SV catalog | Dataset | Novel | Prop. of SVs in public catalog covered | Prop. of common SVs in public catalog covered |
|---|---|---|---|---|
| 1000GP | MESA | 0.932 | 0.160 | 0.844 |
| | 1000GP | 0.932 | 0.160 | 0.844 |
| gnomAD-SV | MESA | 0.673 | 0.088 | 0.628 |
| | 1000GP | 0.670 | 0.087 | 0.627 |

**Table S14. Comparing SVs genotyped in the two datasets (MESA: 2,000 samples; 1000GP: 2,504 samples) with two public SV catalogs.** The table shows the proportion of novel SVs (not present in the public SV catalogs), and the proportion of SVs in the public catalog that are present in our results, either consering all variants or common variants (frequency greater or equal to 5%). Deletions were matched if their reciprocal overlap was at least 30%, insertions if located at less than 200 bp from each other and their size at least 30% similar. Annotated simple repeats were used to extend the matching.

http://cgl.gi.ucsc.edu/data/giraffe/calling/vggiraffe-sv-superpop-af-diff-med10.csv.gz

**Table S15. SVs with strong inter-super-population frequency patterns in the 1000 Genomes Project dataset.** The CSV file contains information about each SV-super-population pair where the allele frequency in the super-population was more than 10% shifted from the median allele frequency across all super-populations.

http://cgl.gi.ucsc.edu/data/giraffe/calling/vggiraffe-sv-eqtl-geuvadis.FDR01.csv

**Table S16. SV-eQTLs in the GEUVADIS dataset.** The CSV file contains information about each SV-gene pair that passed the FDR 1% association threshold.

http://cgl.gi.ucsc.edu/data/giraffe/calling/vggiraffe-sv-2504kgp-pcgenes.csv.gz

**Table S17. SVs overlapping protein-coding genes in the 1000 Genomes Project dataset.** The CSV file contains information about each SVs overlapping coding, untranslated, promoter or introns of protein-coding genes.

### Supplementary Figures

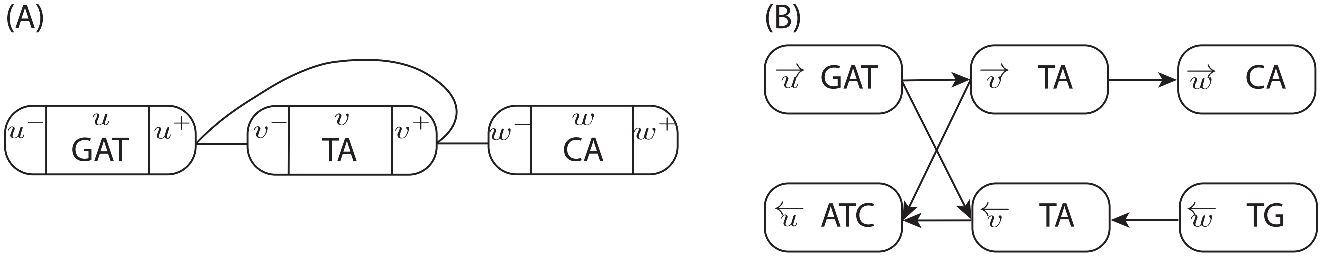

**Figure S1. Sequence graph example.** (A) A sequence graph with three nodes $u$, $v$, and $w$, with labels $\ell(u) = $ GAT, $\ell(v) = $ TA, and $\ell(w) = $ CA. Node sides $x^-$ and $x^+$ are marked for each node $x$. (B) The same graph as a directed graph with each visit $\overrightarrow{x}$ and $\overleftarrow{x}$ a separate node.

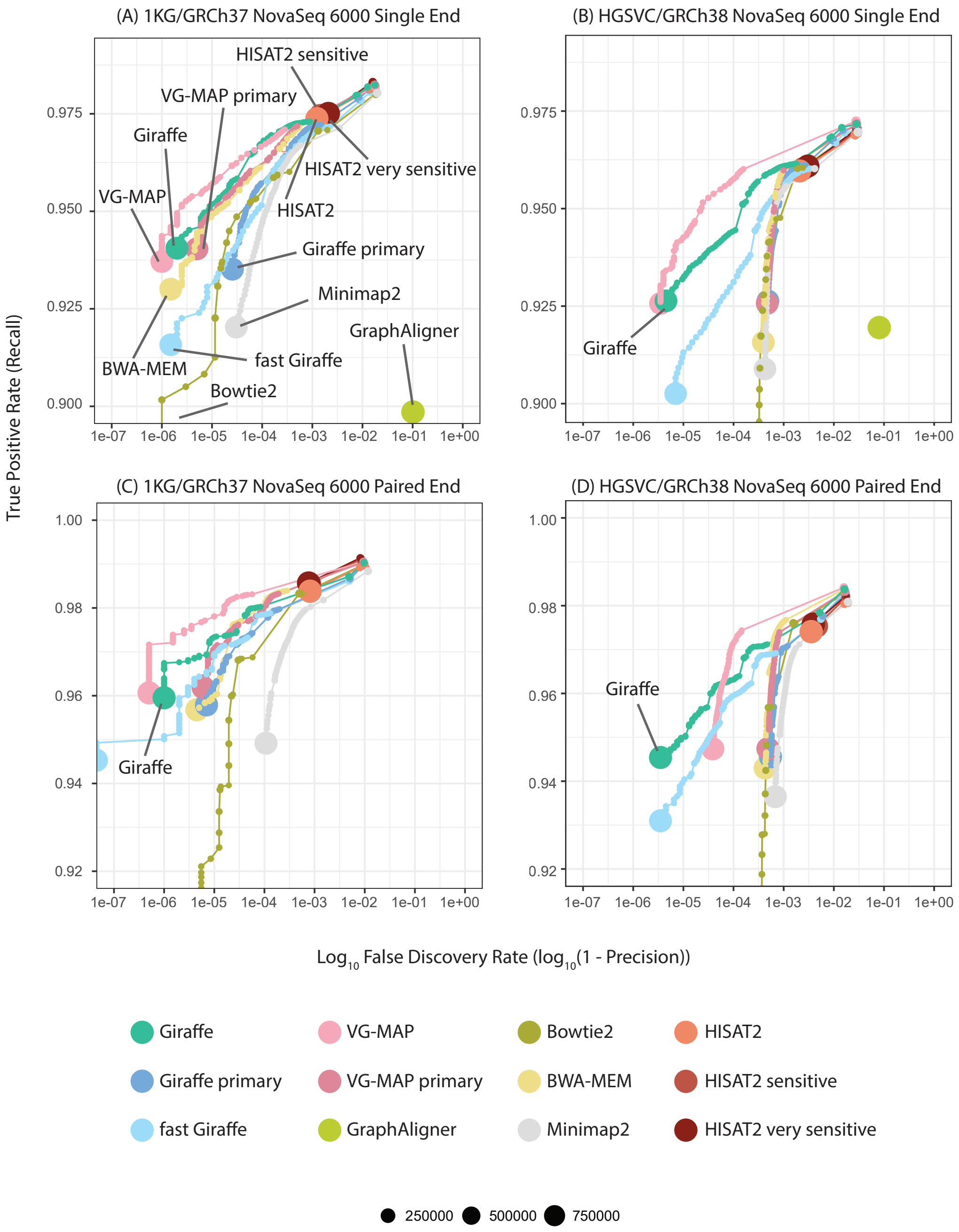

**Figure S2. Simulated read mapping with NovaSeq 6000 reads**. Each panel shows 1-precision (FDR) vs. recall for a simulated read mapping experiment. Reads were simulated to match 150bp Illumina NovaSeq 6000 reads and were mapped either as single ended reads (A,B) or as paired end reads (B,D). We compared graph mappers (Giraffe, VG-MAP, GraphAligner, HISAT2) and linear mappers (BWA-MEM, Bowtie2, Minimap2) mapping to either the 1KG graph and GRCh37 reference (A,C) or the HGSVC graph and GRCh38 reference (B,D). For Giraffe, we mapped to a 64 haplotype sampled GBWT for the 1KG graph and to the full GBWT for the HGSVC graph. In addition to aligning to the 1KG and HGSVC graphs, we also used Giraffe and VG-MAP to align to primary graphs containing only the GRCh37 or GRCh38 reference. We also ran Giraffe and HISAT2 using different pre-defined parametrizations in addition to the default settings: fast Giraffe and HISAT2 sensitive and very sensitive. Reads were stratified by their mapping quality. Each point in the plot represents to a mapping quality value and the size of a point corresponds the number of reads it represents. GraphAligner did not assign mapping qualities, so it is represented as a single point. For the remaining mappers, the mapping quality values varied from 0 to 60, with the majority of reads categorized as mapping quality 60.

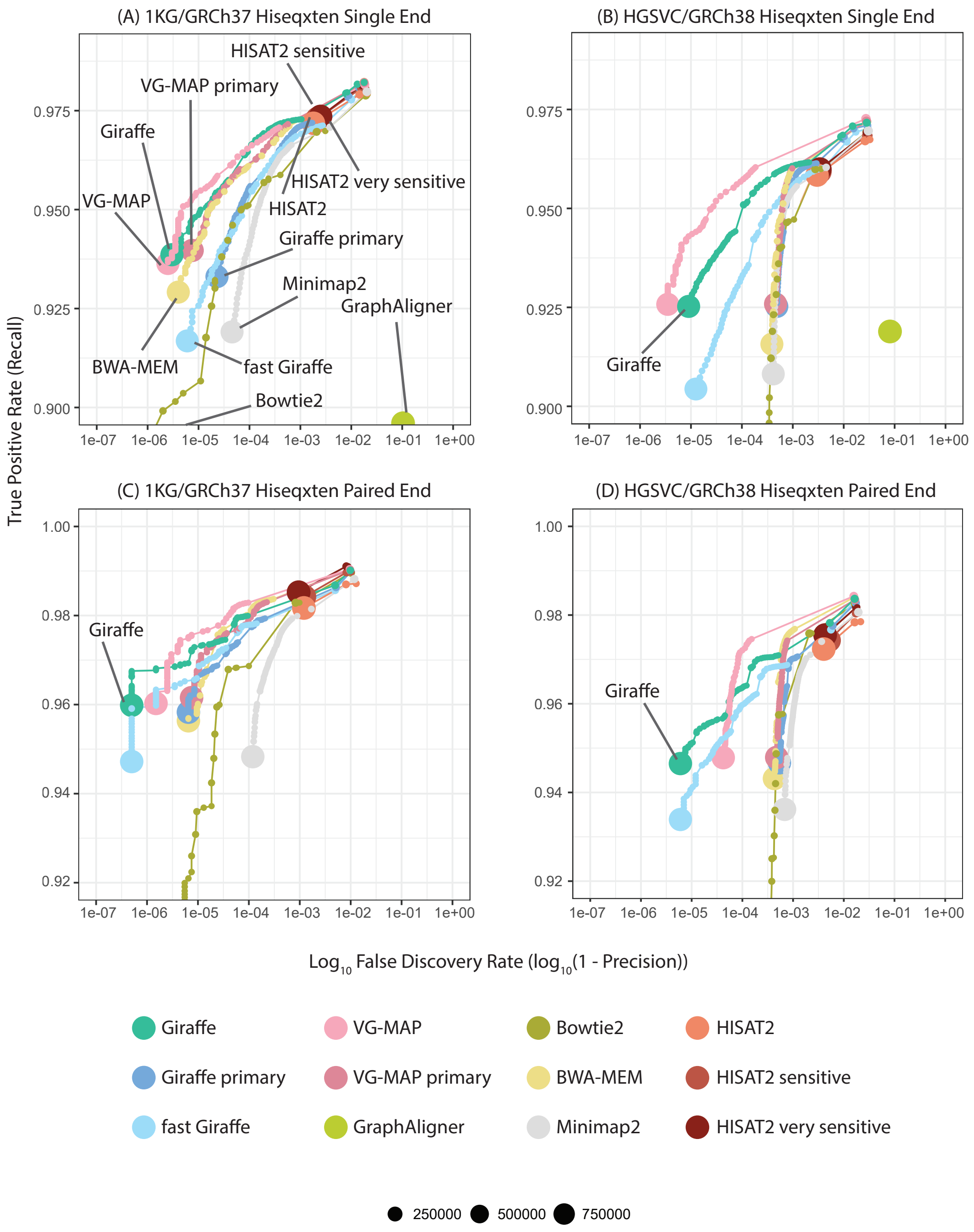

**Figure S3. Simulated read mapping with HiSeq X Ten reads.** Each panel shows 1-precision (FDR) vs. recall for a simulated read mapping experiment. Reads were mapped as single-end (A, B) or paired-end (C, D) to the 1KGP graph and GRCh37 reference (A, C) or the HGSVC graph and GRCh38 reference (B, D).

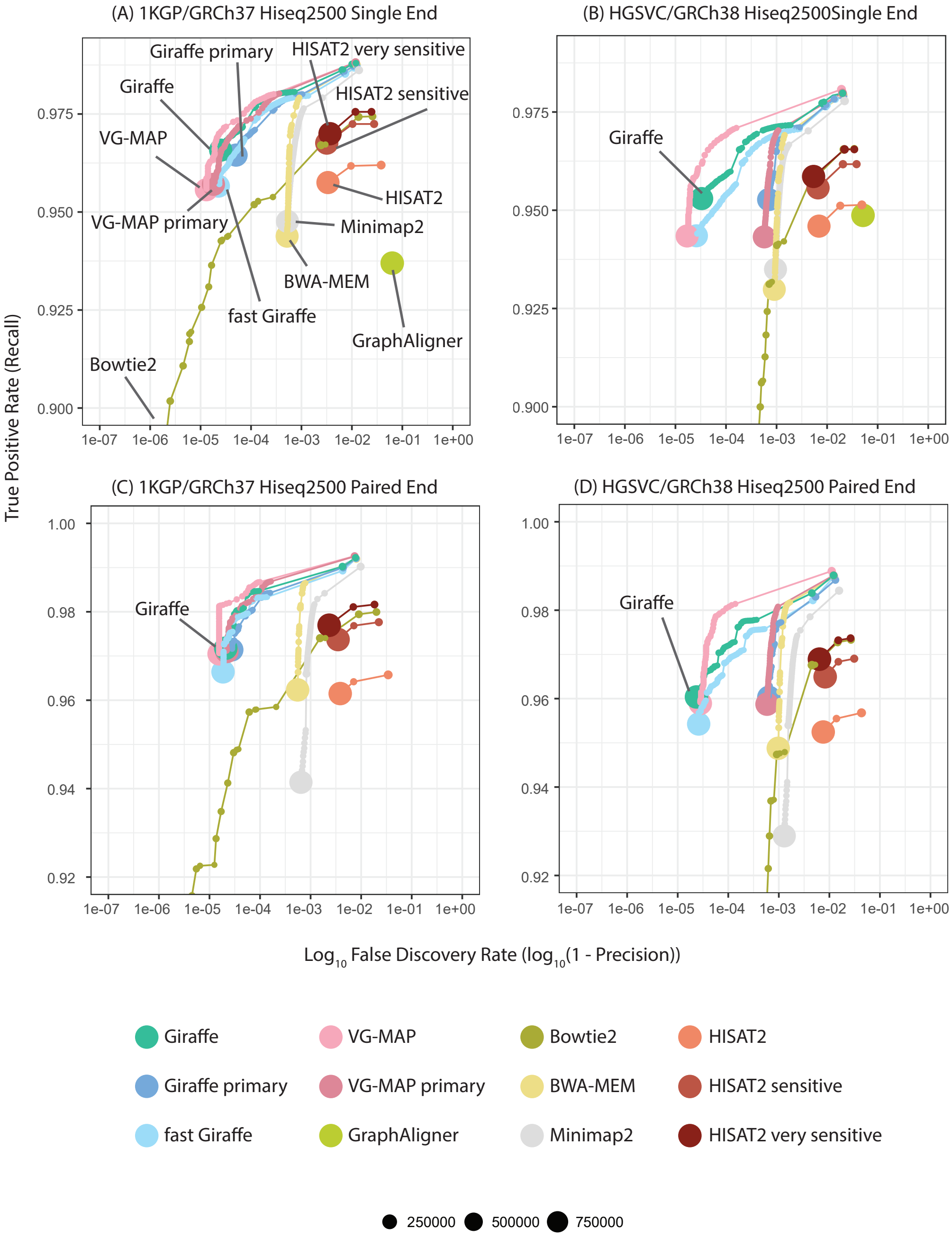

**Figure S4. Simulated read mapping with HiSeq 2500 reads.** Each panel shows 1-precision (FDR) vs. recall for a simulated read mapping experiment. Reads were mapped as single-end (A, B) or paired-end (C, D) to the 1KGP graph and GRCh37 reference (A, C) or the HGSVC graph and GRCh38 reference (B, D).

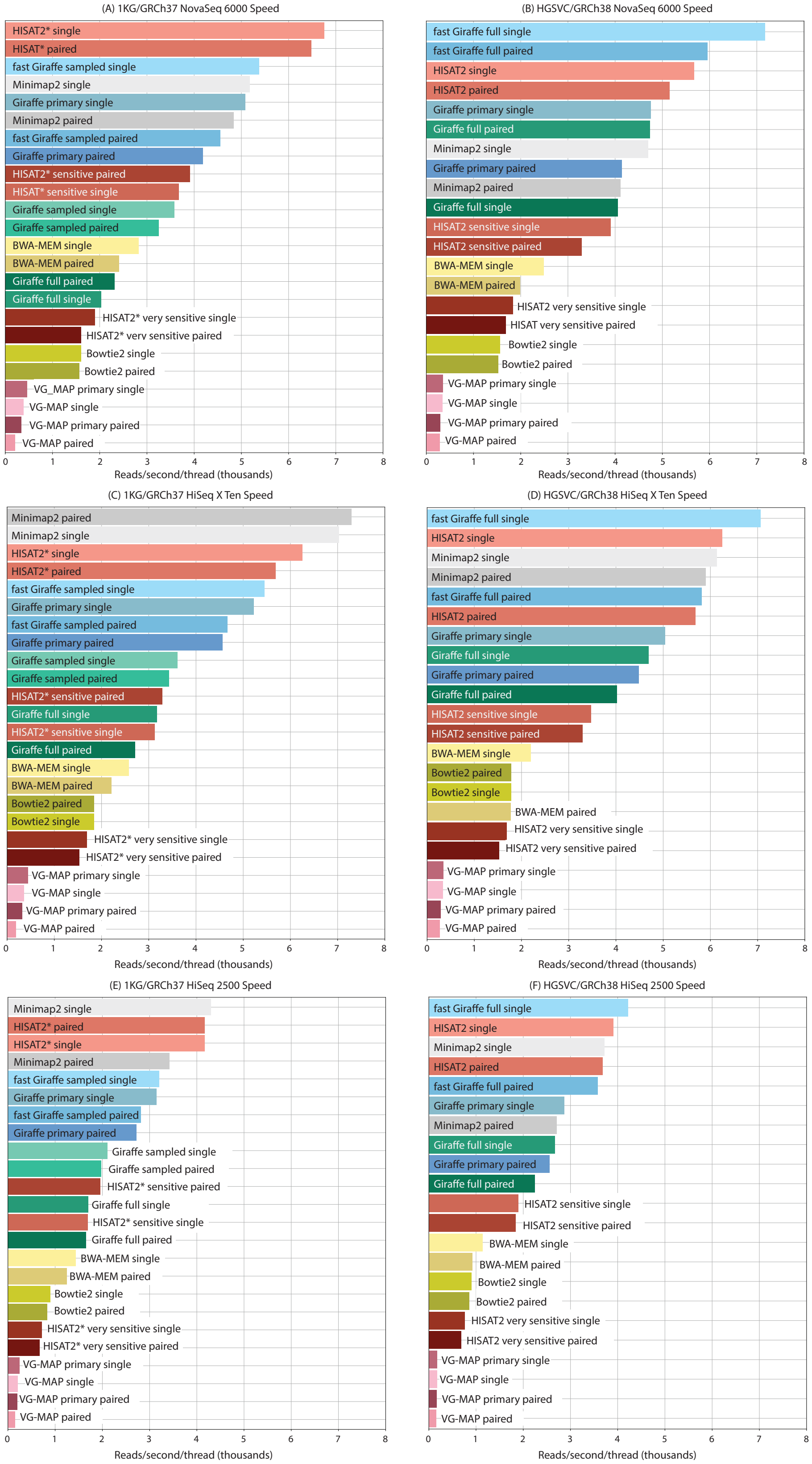

**Figure S5. Mapping speed on human data.** Each mapper was run on a dataset of 1 million real NovaSeq 6000 (A,B), HiSeq X Ten (C,D), or HiSeq 2500 (E,F) reads on a AWS EC2 i3.8xlarge node with 32 vCPUs and 244GB of memory. The speed of mapping in reads per second per thread was determined using the total time spend mapping as reported by each tool. Each tool except Minimap2 was run on 16 threads; Minimap2 was run on 2 threads because it did not use all 16 threads.

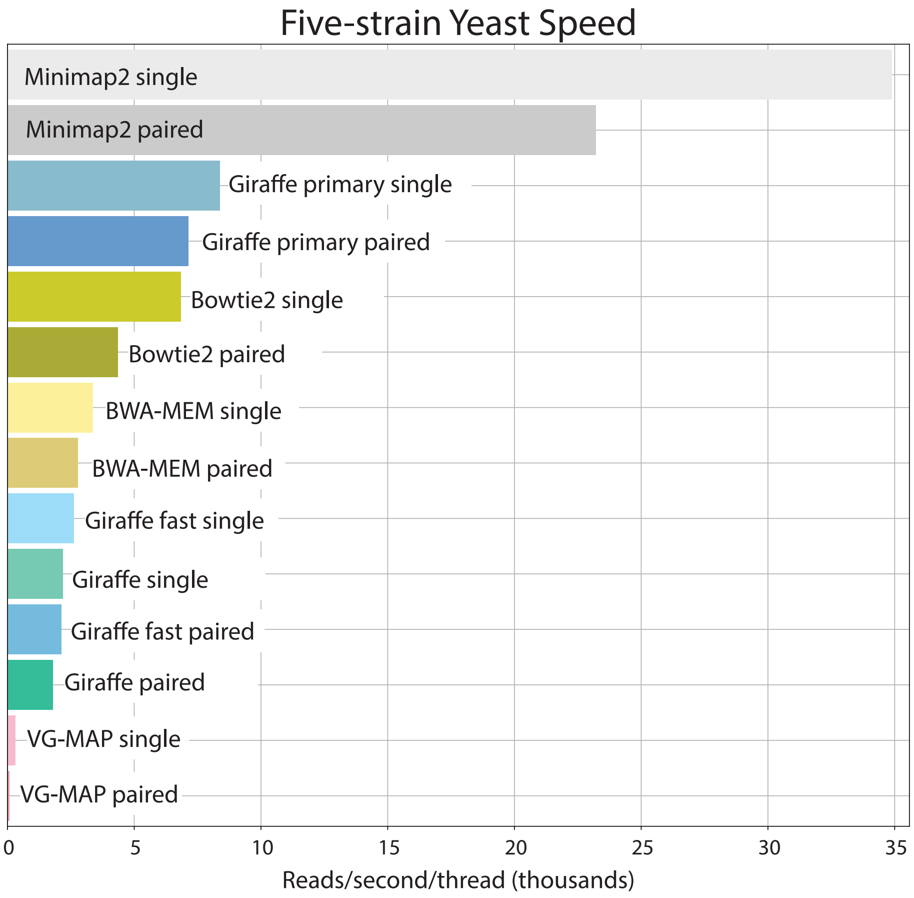

**Figure S6. Mapping speed on yeast data.** Each mapper was run on a dataset of 1 million real HiSeq 2500 reads from the DBVPG6044 strain on a AWS EC2 i3.8xlarge node with 32 vCPUs and 244GB of memory. The speed of mapping in reads per second per thread was determined using the total time spend mapping as reported by each tool. Each tool except Minimap2 was run on 16 threads; Minimap2 was run on 2 threads because it did not use all 16 threads.

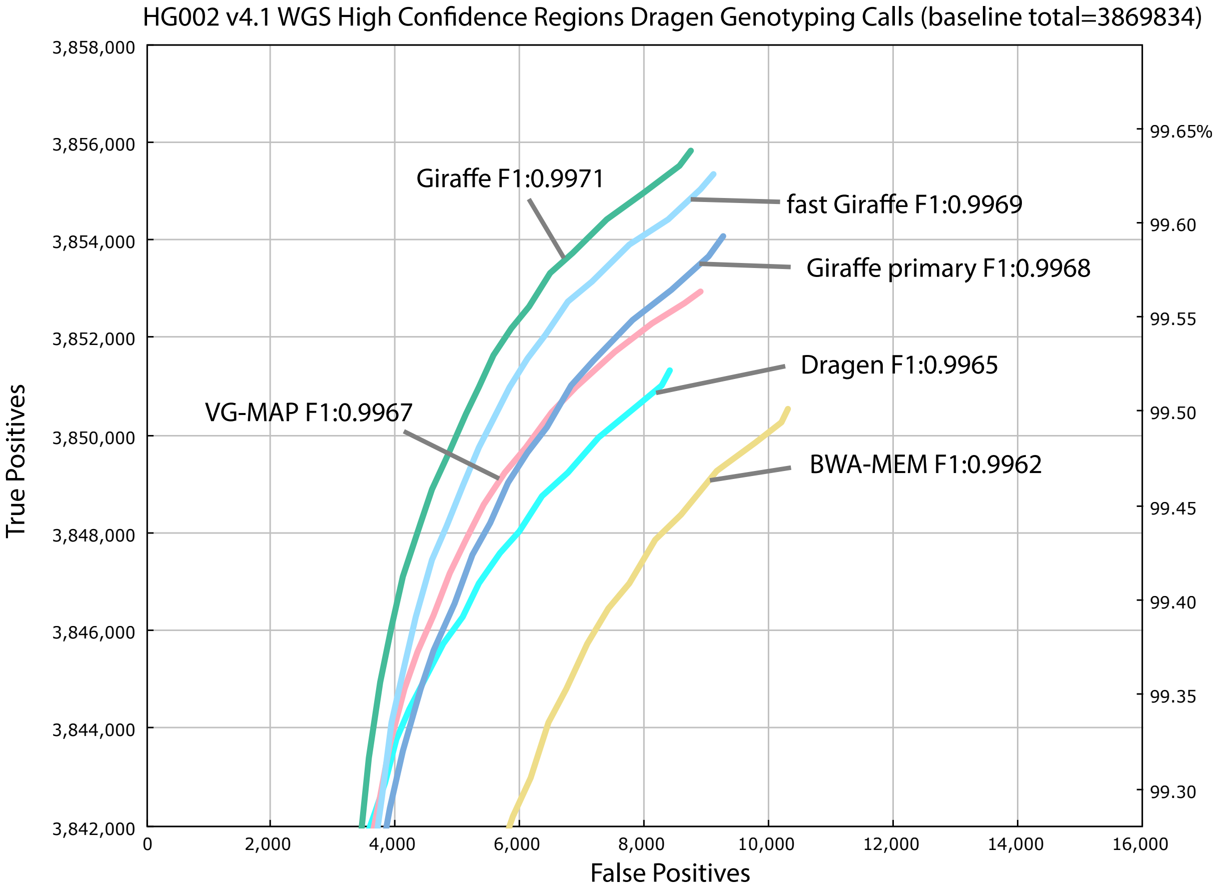

**Figure S7.** True positive and false positive genotypes made using the Dragen genotyper with projected mappings from Giraffe and other mappers, using 2x 250bp reads from the HG002 GIAB sample and evaluated against the HG002 GIAB v4.1 truth variant call sets in high confident regions. The ROC curve discrimination threshold is based on variant call quality.

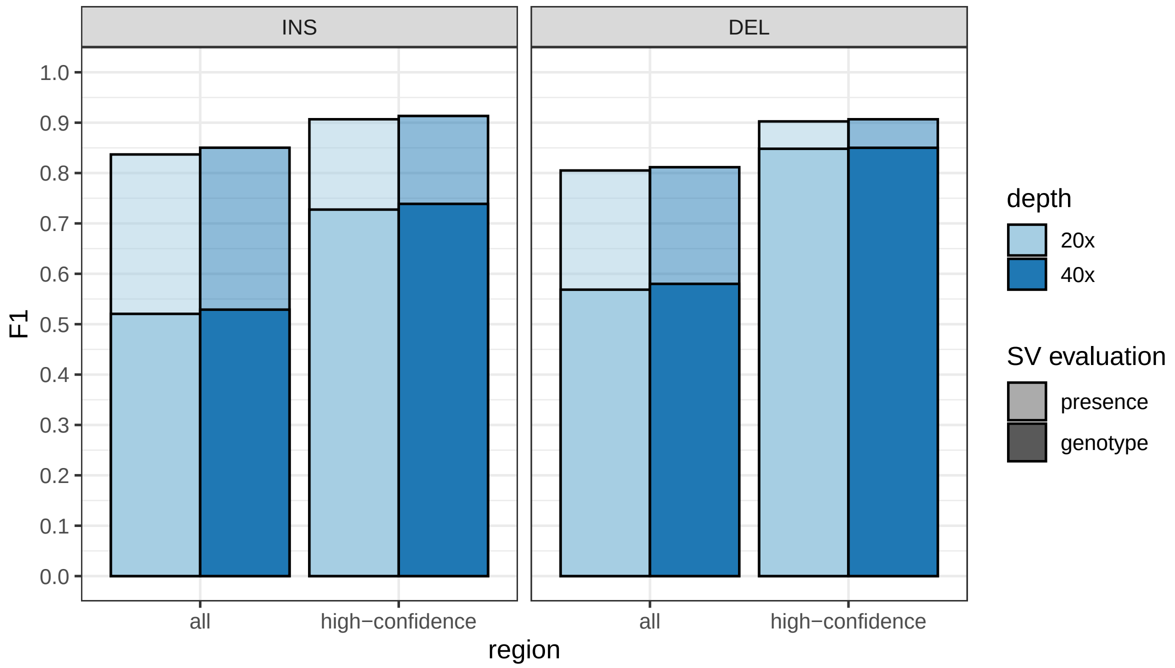

**Figure S8.** **Minimal effect of down-sampling reads to 20x depth on the SV genotyping performance.** HG00514 was genotyped using the SV graph combining HGSVC, SVPOP and GIAB.

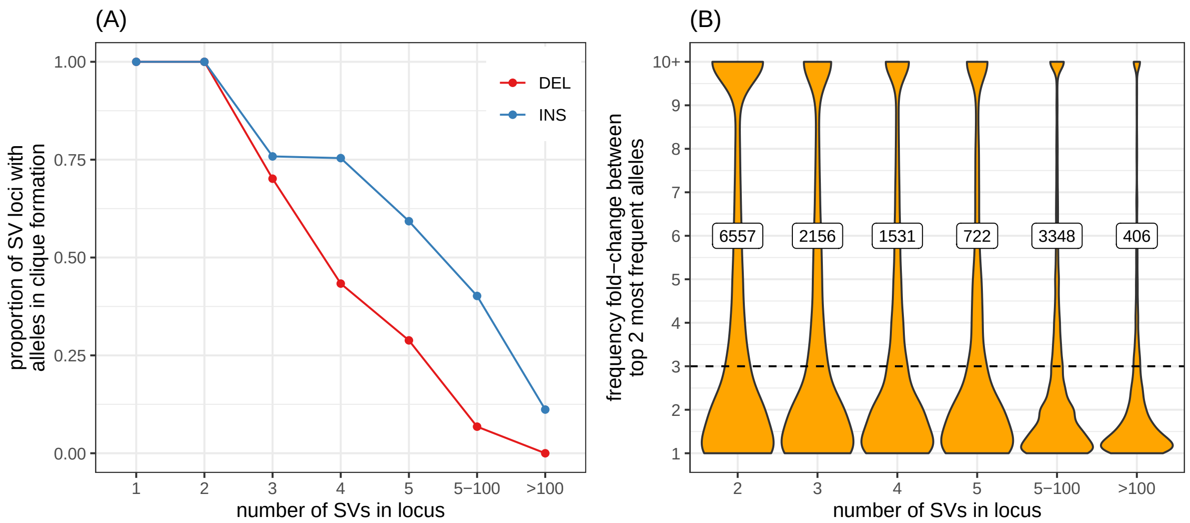

**Figure S9.** **SV alleles and SV sites in the MESA cohort.** (A) Proportion of SV loci in clique formation, i.e. with alleles different by only small variants (SNVs, indels) as opposed to large allelic variation such as VNTRs. The more alleles in a SV site the more likely it is to show VNTR-like allelic patterns. (B) Fold-change between the major allele and the second most frequent allele in a SV loci (y-axis), grouped by the number of alleles in the locus (x-axis). The number of loci with a fold-change higher than 3 (dotted line) are labeled.

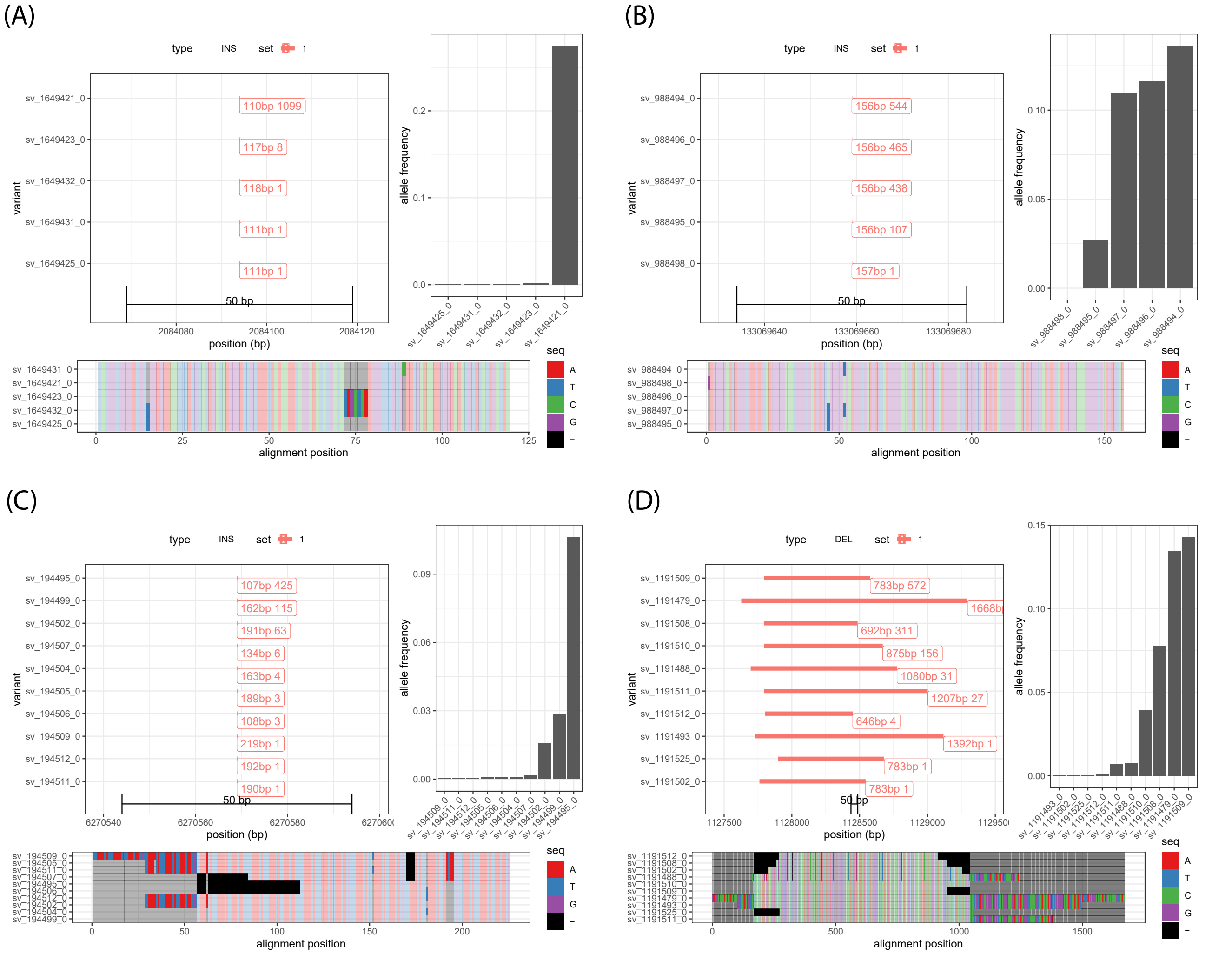

**Figure S10. Four examples of SV sites in the MESA cohort.** For each site, the top left panel shows a representation of their configuration in the genomic region, the top right panel shows the distribution of allele frequencies across the 2000 MESA samples, and the bottom panel shows the result of a multiple-sequence alignment of the inserted/deleted sequences. In (A) and (B), the insertions' alleles are only different due to small variants while (C) and (D) shows an insertion and deletion site with significant size variation between the alleles. In (A), there is one major allele while multiple alleles of (B) are frequent in the population.

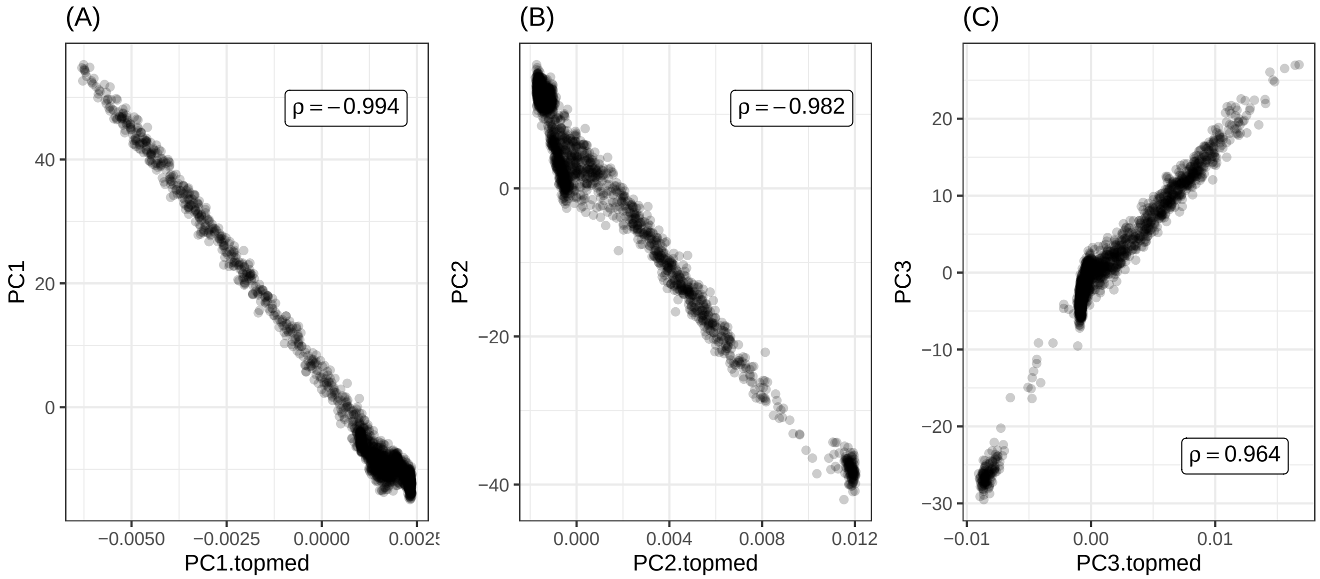

**Figure S11. Principal component analysis using the SV genotypes (y-axis) in the MESA dataset compared to the principal components derived from TOPMed-wide SNVs (x-axis).** (A), (B), (C) compare the first, second and third components.

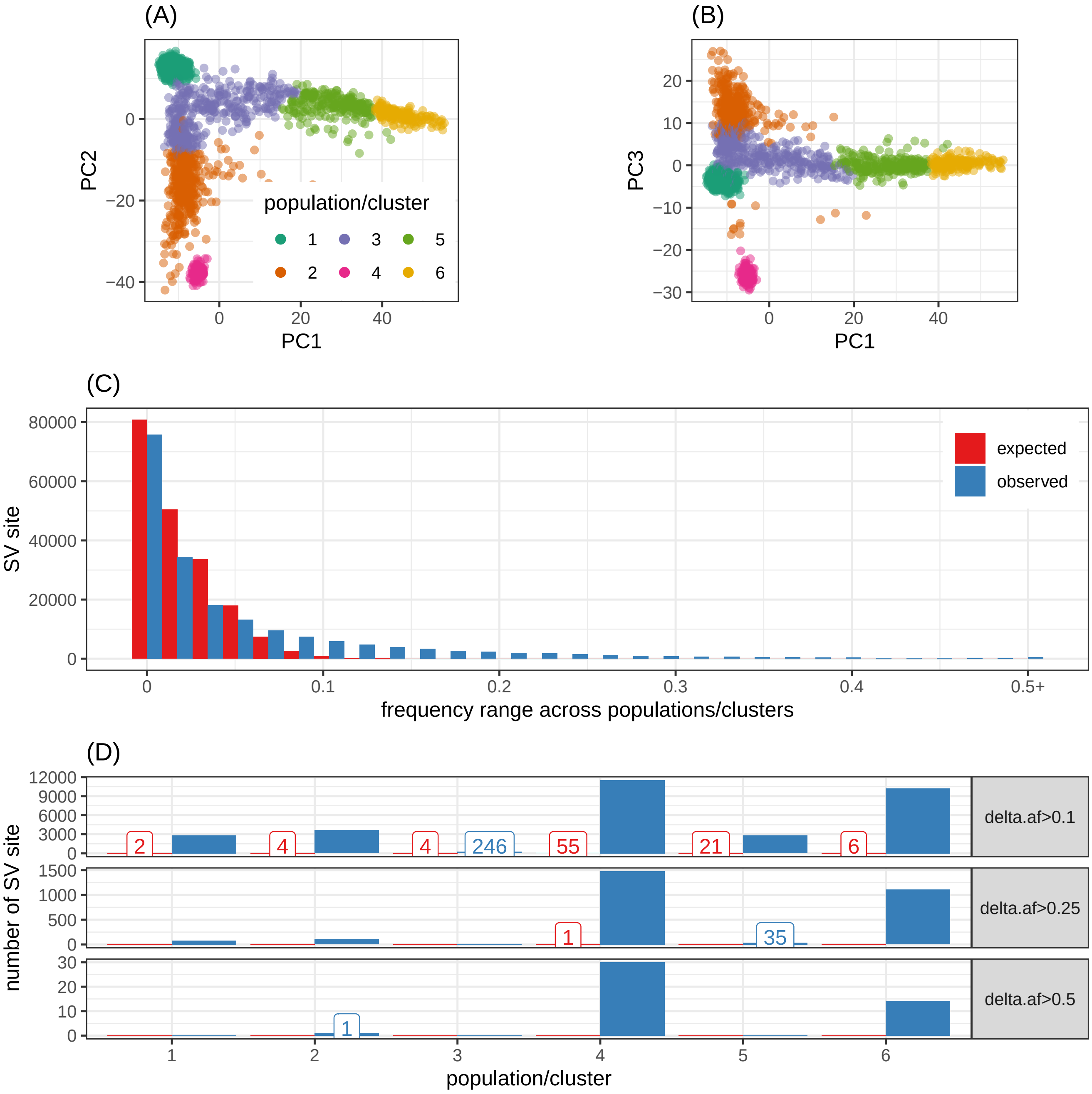

**Figure S12. SV sites with population specific frequency patterns in the MESA cohort.** The two thousand MESA samples were clustered in 6 clusters based on the top 3 principal components (A-B). (C) The range of frequency across the sample clusters was larger than expected (permutation) for thousands of SV sites. (D) Two clusters (x-axis) show high number of population-specific SV sites (y-axis), defined as deviating from the median allele frequency by at least 10%, 25%, or 50% (panels).

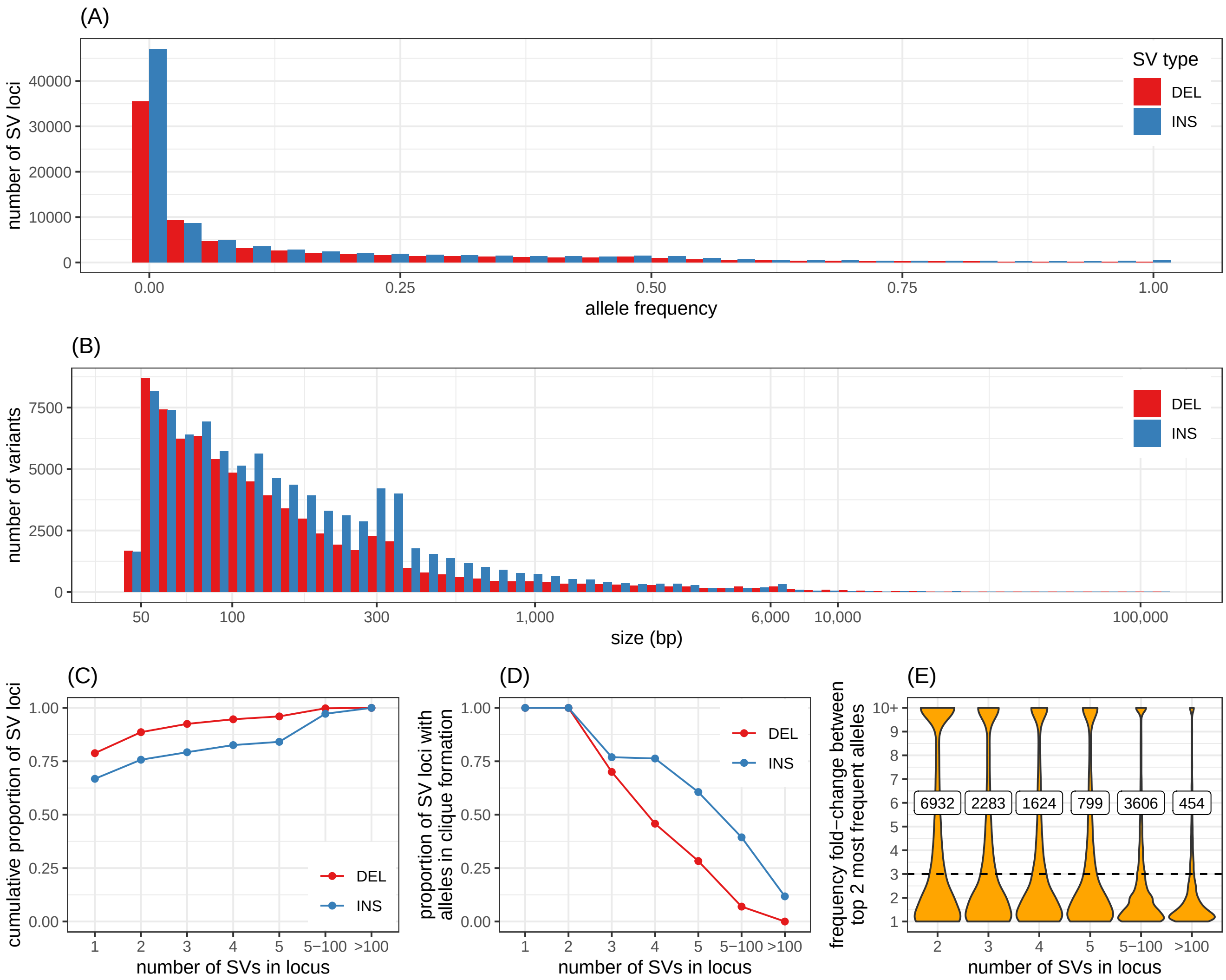

**Figure S13.** **Structural variants in 2,504 unrelated individuals from the 1000 Genomes Project.** (A) Allele frequency distribution of the major allele for each SV loci. (B) Size distribution of the major allele for each SV loci. (C) Cumulative proportion of SV loci depending on the maximum number of alleles (x-axis) in the locus. (D) Proportion of SV loci in clique formation, i.e. with alleles different by only small variants (SNVs, indels) as opposed to large allelic variation such as VNTRs. The more alleles in a SV site the more likely it is to show VNTR-like allelic patterns. (E) Fold-change between the major allele and the second most frequent allele in a SV loci (y-axis), grouped by the number of alleles in the locus (x-axis). The number of loci with a fold-change higher than 3 (dotted line) are labeled.

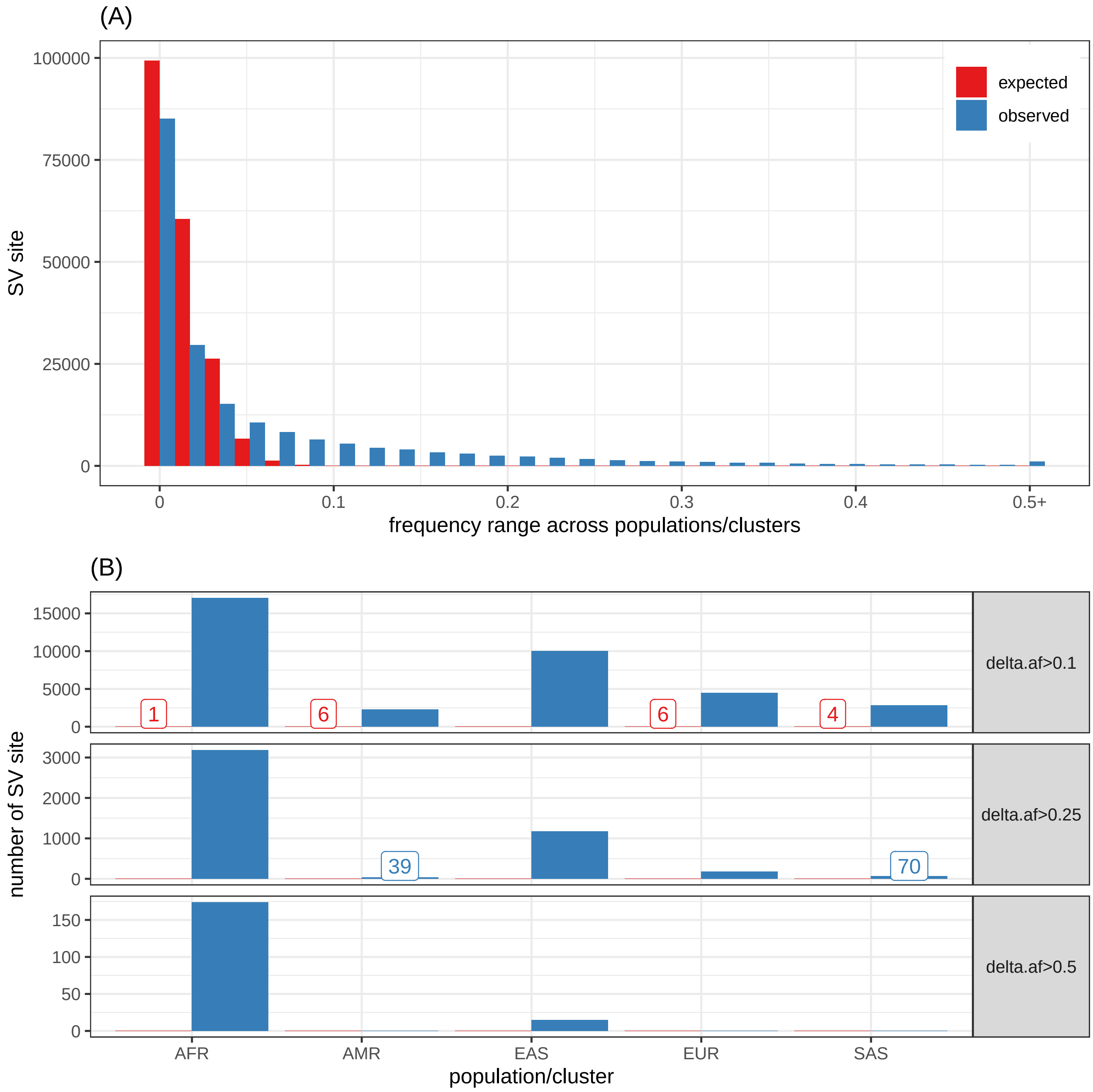

**Figure S14. SV sites with population specific frequency patterns in the 2,504 individuals from the 1000 Genomes Project.** (A) The range of frequency across the super-population was larger than expected (permutation) for thousands of SV sites. (B) Super-populations (x-axis) show high number of population-specific SV sites (y-axis), defined as deviating from the median allele frequency by at least 10%, 25%, or 50% (panels).

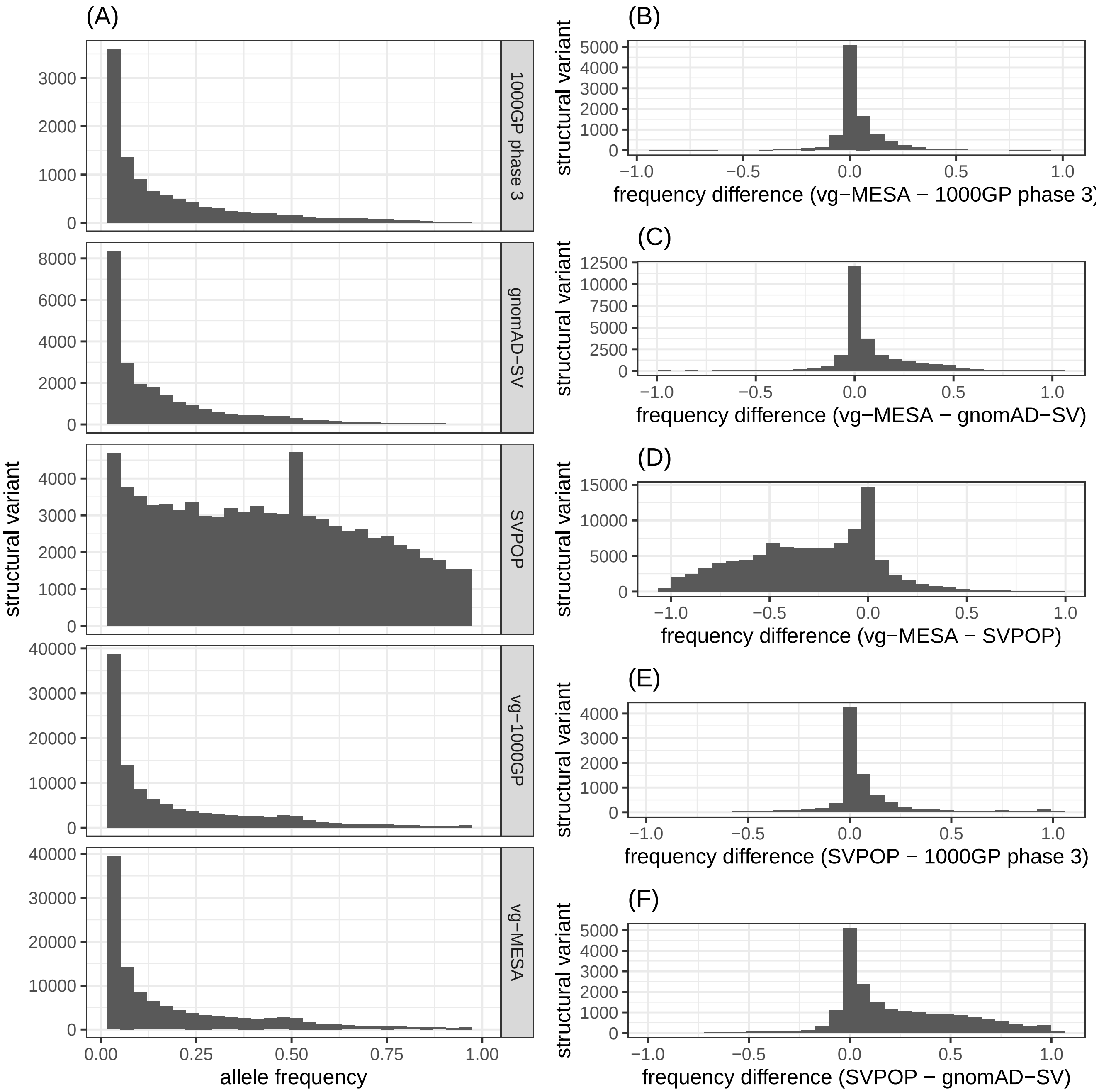

**Figure S15. Allele frequency of SVs in this study and public studies.** (A) Allele frequency distribution of common variants (frequency >1%) in each dataset. (B-D) show the difference in frequency between our results in the MESA cohort and each of the three public catalogs (1000 Genome Project phase 3, gnomAD-SV, SVPOP). (E-F) show the difference in frequency between SVPOP and the two other public catalogs.

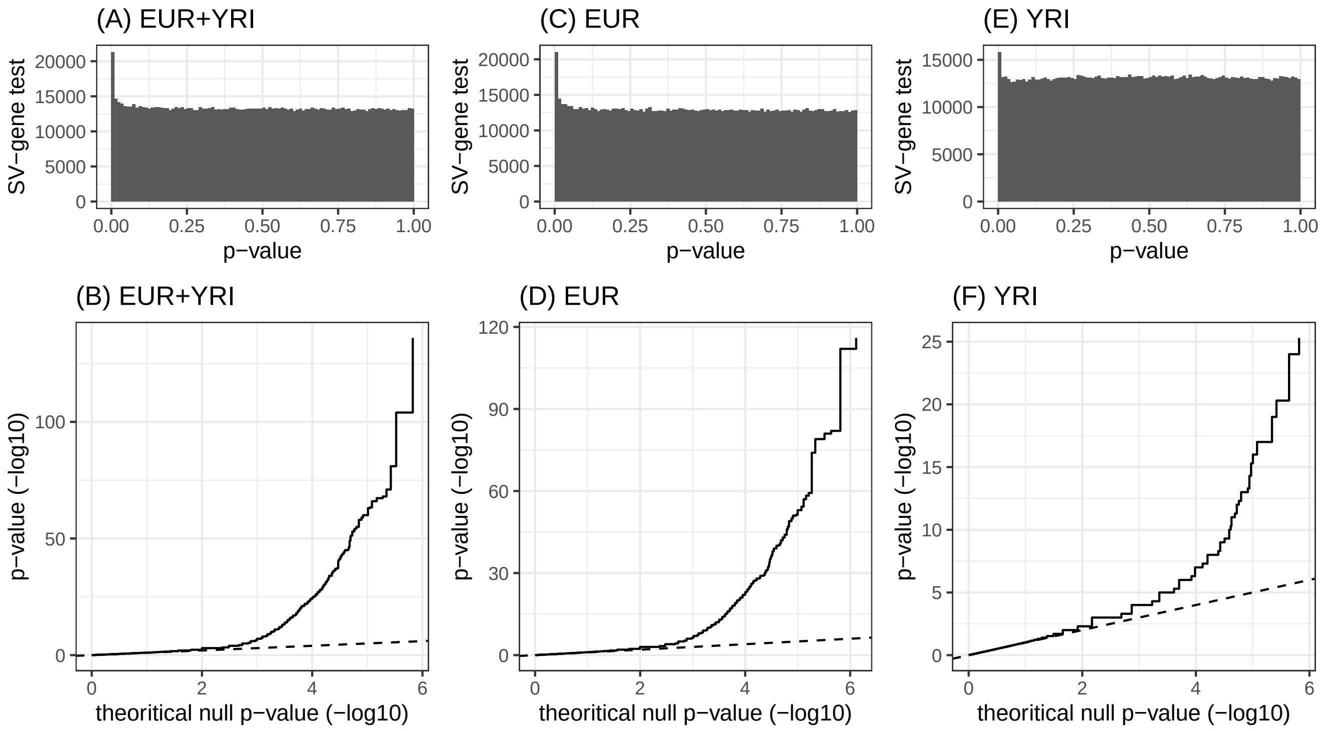

**Figure S16.** **P-value distribution of the SV vs gene expression tests in the GEUVADIS dataset.** Three analysis were performed: joint European and Yoruba populations (A-B), European populations only (C-D), and the Yoruba population (YRI) only (E-F). (A,C,E) show the p-value distribution, uniform except for the peak of low p-values. (B,D,F) show the QQ plots for the three analysis.

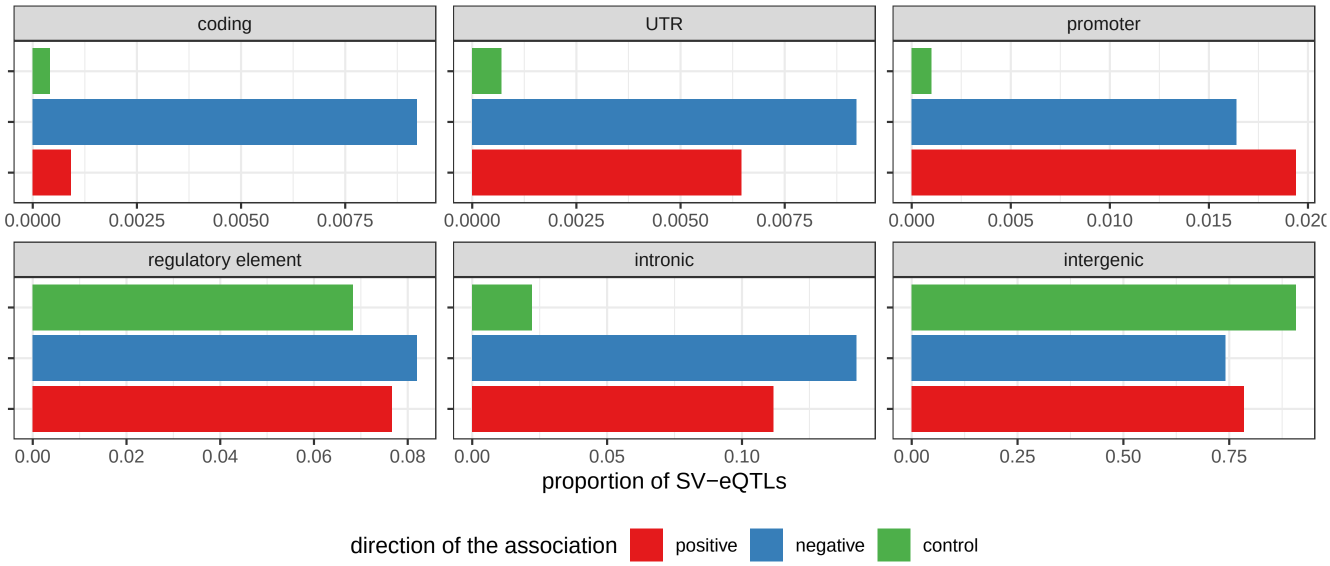

**Figure S17.** **Location of the SV-eQTLs around the associated gene** The x-axis reports the proportion of SV-eQTLs overlapping each annotation group (panels). A SV-eQTL is either positively (red) or negatively (blue) associated with gene expression, i.e. each additional SV allele correlates with increased or decreased gene expression respectively. The control bar (green) represents the expected distribution of common SVs around control genes. Control genes were chosen to match the size distribution of genes involved in eQTLs. *UTR: untranslated region.*
